# Molecular and ecological determinants of effective reassortment in orthohantaviruses

**DOI:** 10.64898/2026.06.10.731004

**Authors:** Ricardo Rivero, David Simons, Lambodhar Damodaran, Irene Karegi, Sarah Gurev, Daniel J. Becker, Dan L. Warren, Nicola F. Müller, David A. Rasmussen, Stephanie N. Seifert

## Abstract

A central unsolved problem in RNA virus evolution is understanding why some viral reassortants establish and persist while others do not. To answer this question, we reconstructed reassortant histories across 553 viral genomes from seven orthohantavirus species between 1983 and 2026 using phylogenetic reconciliation and molecular dating. We found that the frequency of retained reassortants varied among orthohantaviruses. For example, reassortment ranged from absent in Andes virus (0/61) to frequent in Dobrava-Belgrade (12/18), Sin Nombre (20/34), Seoul (47/101), Puumala (28/69), and Tula (27/71) viruses, showing that effective reassortment is not a genus-wide constant. Our Bayesian hierarchical models identified local host overlap as the strongest ecological factor associated with viral reassortment (posterior mean log-odds = 0.297, 94% highest-density interval [HDI] 0.039–0.561; increasing to 0.356, 94% HDI 0.118–0.622 among subnationally georeferenced genomes), while cross-segment linkage and terminal RNA structure serve as a molecular filter. We found that the probability of reassortment establishment is highest when ecological opportunity is paired with molecular permissiveness, with their interaction term inferred as the strongest signal in our establishment models (posterior mean log-odds = 0.866, 94% HDI 0.004–1.826, posterior probability = 0.97). These results suggest that reassortment in orthohantaviruses is a sequentially filtered evolutionary process in which divergent lineages must first meet in a host through ecological overlap, exchange segments that are molecularly compatible, and do so within a lineage background permissive to establishment in the host population.

## 1 Introduction

Negative-sense RNA viruses experience dampened rates of homologous recombination, with implications for their ability to effectively purge deleterious mutations or combine beneficial mutations^1^. However, viruses with a negative-sense genome broken into genome segments can generate reassortant genotypes via segment exchange in a single mutational step^2, 3^. Most reassortant genotypes do not persist in host populations as only a minority of possible reassortants are detected in natural populations. Here we ask which biological conditions favor establishment of viral reassortants in host populations.

Orthohantaviruses are an excellent system in which to address this question. They have a tripartite negative-sense RNA genome consisting of the S (nucleocapsid protein), M (glycoprotein precursor) and L (RNA-dependent RNA polymerase; RdRp) segments and they are maintained in small-mammal reservoirs, primarily rodents^4^. Several orthohantaviruses cause zoonotic disease following spillover into humans, including the Hantavirus Pulmonary Syndrome-associated orthohantaviruses in the Americas and the Hemorrhagic Fever with Renal Syndrome-associated orthohantaviruses throughout Eurasia. As such, full genome sequences span decades of surveillance with cosmopolitan distributions. Reassortment has been documented in several orthohantaviruses^3^, providing both the biological variation and the data density needed to model reassortment effectively. Strong host-virus phylogenetic structure had been considered a hallmark of the genus, yet strict codivergence with associated host species is not observed^5–7^. Instead, host switching, geographic overlap among reservoir hosts, and reassortment have all contributed to diversification in orthohantaviruses^3, 5, 6, 8^.

Reassortment has been documented in both natural and experimental hantavirus systems. Natural reassortants have been reported in several orthohantavirus species, including Sin Nombre virus (SNV; *Orthohantavirus sinnombreense*), Puumala virus (PUUV; *Orthohantavirus puumalaense*), Dobrava-Belgrade virus (DOBV; *Orthohantavirus dobravaense*), Hantaan virus (HTNV; *Orthohantavirus hantanense*), and Seoul virus (SEOV; *Orthohantavirus seoulense*)^3, 9–15^. Most well-supported events involve variants within the same species^3, 9, 12, 13, 15, 16^, although reassortants between HTNV and SEOV have been reported, showing that inter-species reassortment can occur^17^. In contrast, evidence for viral reassortment in Andes virus (ANDV; *Orthohantavirus andesense*) is sparse^18, 19^. In PUUV, longitudinal studies of bank vole populations found frequent reassortment among co-circulating viral lineages^15, 20, 21^, while studies of DOBV identified cases where the evolutionary histories of different genome segments did not match among host-associated genotypes^9^. Experimental coinfections further show that viable hantavirus reassortants can be recovered, but not all segment combinations appear equally, suggesting variable compatibility^3, 22, 23^. Notably, across these experiments, recovered single-segment reassortants have generally involved M-segment exchange while S and L remain paired. This pattern suggests that the nucleocapsid-polymerase-RNA replication module imposes stronger compatibility constraints than the glycoprotein segment^3, 22, 24–27^.

Together, these observations suggest a sequential filter for establishment of reassortants where host ecology and behavior must first bring compatible viral lineages into contact to give rise to coinfected host cells. Because orthohantaviruses are maintained in small-mammal reservoirs, many with multiple possible host taxa^8^, we hypothesized that opportunities for coinfection depend on host overlap, local infection prevalence, and the spatial structure of host communities. Investigating host contributions to reassortment is made possible by our recent effort to synthesize all published surveillance data on arenaviruses and orthohantaviruses into a single database^8^. Within species, they also depend on whether distinct viral lineages co-occur in the same host populations. Second, exchanged segments must remain functionally compatible within the reassortant genomic background. Once host cells are coinfected, viral segments must be compatible for virion assembly and packaging and maintain functional 5’ and 3’ UTR sequences compatible with the RdRp. Importantly, these filters may not operate identically across hantavirus species; even if a host community provides opportunities for mixed infection, few reassortants may persist if the segment backgrounds are incompatible.

This study evaluates two linked hypotheses. The ecological-opportunity hypothesis posits that reassortment is more effective when host overlap, reservoir co-occurrence, and local transmission context increase the likelihood of mixed infection. The molecular-filter hypothesis asserts that, even when coinfection is possible, only certain segment constellations persist because reassortment is limited by cross-segment coadaptation, linkage structure, terminal regulatory architecture, and species-specific genome backgrounds. Accordingly, the analysis focuses on effective reassortment, defined as exchanged segment constellations that persist sufficiently to be detected in descendant genomes, identifying the ecological and molecular conditions that enable reassortant genomes to persist.

## 2 Results

### 2.1 Effective reassortment varied markedly among orthohantavirus species

We assembled a harmonized orthohantavirus dataset spanning 1983–2026 across seven focal species: ANDV, DOBV, HTNV, PUUV, SEOV, SNV, and Tula virus (TULV; *Orthohantavirus tulaense*). The harmonized variant set contained 2,131 records; requiring S, M, and L segment representation and at least 70% sequence completeness reduced this to 638 variants, and time-calibration filtering produced 553 complete genomes for segment-tree reconciliation (Fig. 1a).

**Figure 1:**
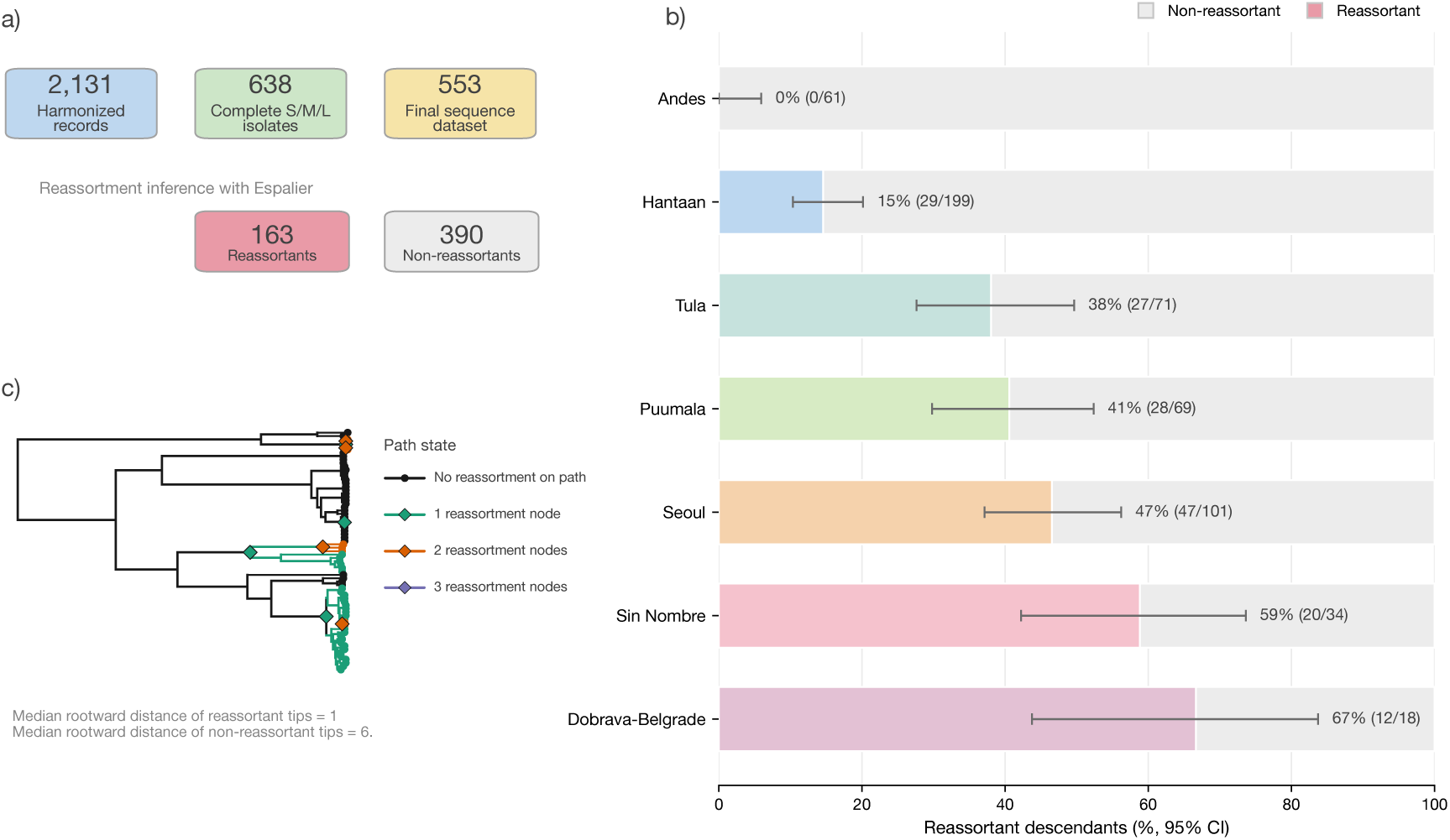
**a,** Assembly and label-generation workflow. Harmonized records were filtered to complete S/M/L isolates and then to the final 553-genome sequence dataset; Espalier-based reassortment inference assigned 163 reassortants and 390 non-reassortants. **b,** Species-level retained reassortment fractions and counts (*k/n*) among the 553 genomes. Bars show the percentage of reassortant (colored, with 95% Wilson binomial confidence intervals) and non-reassortant (gray) descendants for each focal orthohantavirus species. **c,** Tip-to-root reassortment-path validation on the time-calibrated Tula virus S-segment phylogeny (*n* = 71). Colored diamonds and branches show the cumulative number of rootward reassortment encounters along each path (0, 1, 2, or 3 nodes), with reassortant status restricted to direct descendants of reassortment nodes (*d ≤* 2 internal nodes).

We reconstructed reassortment ancestry from time-calibrated segment trees using Espalier^28^, an ancestral recombination graph (ARG)-based phylogenetic reconciliation framework that identifies statistically supported reassortment events while reducing the influence of segment-tree discordance attributable to phylogenetic uncertainty. We then defined reassortment-positive tips as those descending from inferred reassortment nodes. To avoid inflating reassortment counts from large descendant clades, we applied a denesting step in which each tip was traced towards the root and labeled positive only when the nearest retained reassortment node was its direct ancestral event. Tips separated from a reassortment node by more than one intervening internal node were counted as negative. The final explanatory dataset contained 163 reassortants and 390 non-reassortants (Fig. 1a). Path-based validation confirmed that positive labels reflected nearby retained reassortment ancestry: across the 553 genomes, 113 reassortant tips lay immediately below an inferred reassortment node (*d* = 1) and 50 were separated by a single intervening node (*d* = 2), whereas non-reassortant tips either had no ancestral reassortment node (119*/*553) or encountered only deep ancestral events separated by a median of 6 internal nodes (Fig. 1c; Supplementary Table 8).

Retained reassortment varied sharply among species. It was undetected in the analyzed ANDV genomes (0/61; 0.0%, 95% confidence interval [CI] 0.0–5.9%), low in HTNV (29/199; 14.6%, 95% CI 10.3–20.1%), and common in DOBV (12/18; 66.7%, 95% CI 43.7–83.7%), SNV (20/34; 58.8%, 95% CI 42.2–73.6%), SEOV (47/101; 46.5%, 95% CI 37.1–56.2%), PUUV (28/69; 40.6%, 95% CI 29.8–52.4%), and TULV (27/71; 38.0%, 95% CI 27.6–49.7%) (Fig. 1b; Supplementary Table 8). These fractions are descriptive summaries of the sampled genomes, as sample size, host composition, temporal span, and phylogenetic depth vary among species (Supplementary Table 3; Supplementary Fig. 3). Normalizing the 73 de-nested minimal reassortment origins by mean segment total branch length (in substitutions per site) accounted for differences in evolutionary time sampled and post-origin clonal expansion, yielding origin rates ranging from 0.00 origins per substitution per site in ANDV to 4.52 in DOBV, 5.55 in SNV, 7.26 in TULV, 8.31 in HTNV, 9.08 in PUUV, and 15.37 in SEOV (Supplementary Table 8). Together, these differences show that the studied orthohantavirus cohorts span reassortment-restrictive and reassortment-permissive regimes, consistent with retained reassortment being shaped by species-specific ecological or molecular filters (Fig. 1).

### 2.2 Ecological opportunity and molecular compatibility defined distinct reassortment filters

The species-level differences in retained reassortment suggested that ecological opportunity and molecular compatibility act as separable filters. Segment-level Procrustean Approach to Cophylogeny (PACo) tests detected non-random host-virus association for S, M, and L histories (all permutation *p <* 0.001; Supplementary Table 1, Supplementary Fig. 10), but residual spread and segment-specific link patterns did not support strict codivergence across all segment histories. Instead, segment-specific mismatches were consistent with partial codivergence, host switching, reservoir sharing and reticulate viral ancestry. This supports an ecological-opportunity framework in which reassortment is most likely when related viral lineages have been brought into overlapping host populations or geographic spaces.

Local host overlap and local prevalence captured different aspects of this opportunity. Local host overlap summarized the predicted co-occurrence of plausible hantavirus hosts near sampled genomes (Fig. 2a, b), whereas local prevalence summarized nearby evidence for active hantavirus circulation^8^. High prevalence in one dominant reservoir may amplify a single viral background without increasing exposure to divergent segment lineages; local host overlap more directly represents the setting in which distinct viral lineages can be brought into contact (Fig. 2c).

**Figure 2:**
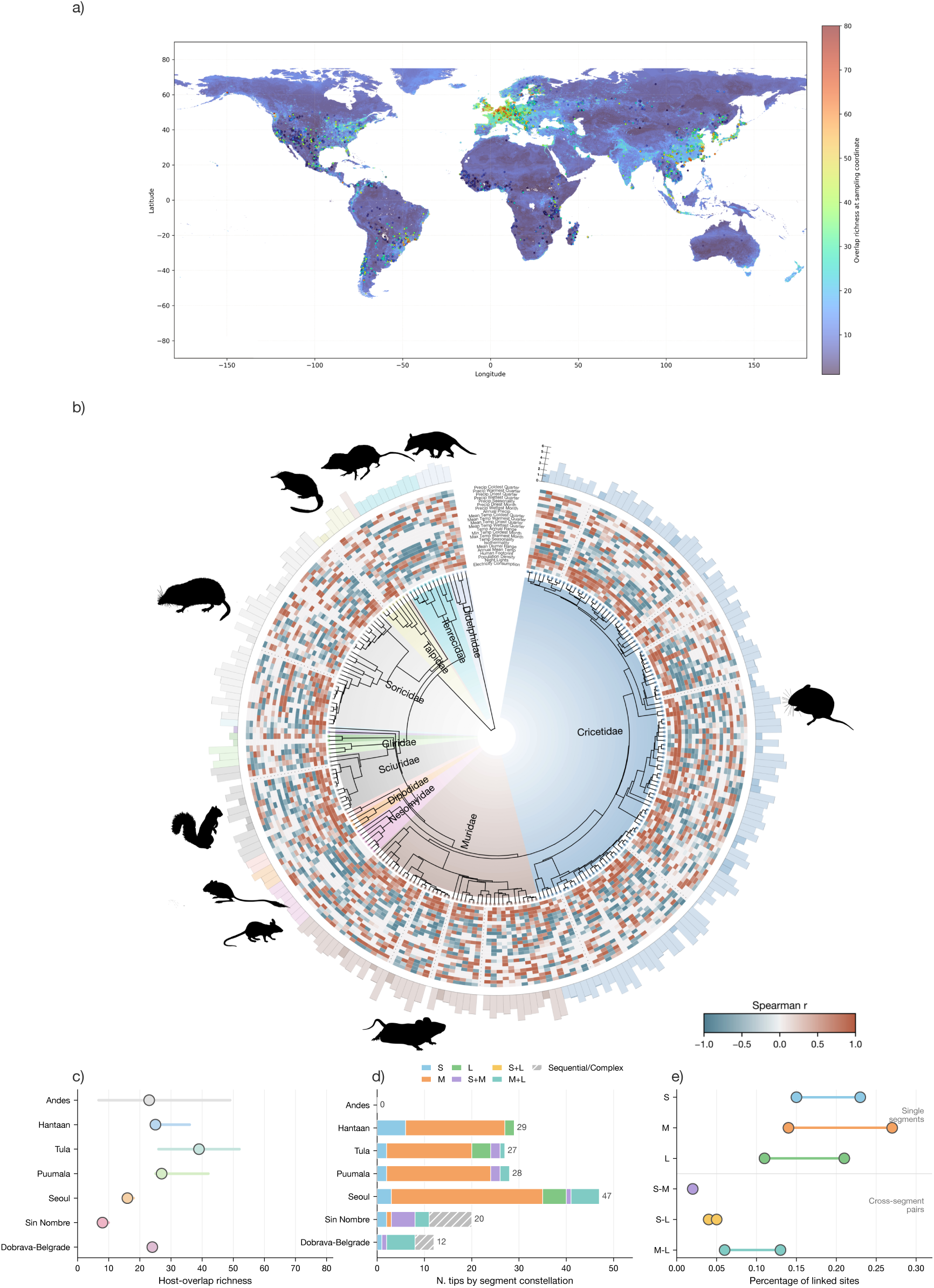
**a,** Global predicted host-overlap richness 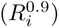 and virus sampling localities at 30-arc-second resolution, summarizing macroecological opportunity for host sharing. **b,** Mammalian host phylogeny annotated with environmental covariates and host-SDM partial-dependence response summaries; heatmap cells show unitless Spearman correlations (*r*, from *−*1 to +1) between predictor values and host-SDM partial-dependence responses across species, and outer radial bars show the number of documented viral/surveillance links per host taxon. **c,** Local host-overlap richness (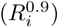 by virus species (points: medians; horizontal bars: interquartile ranges). **d,** Segment constellations among reassortant genomes (*n* = 163) by species. **e,** Range (minimum to maximum percentage across species) of polymorphic nucleotide site pairs with significant link-age disequilibrium (Cramér’s *V ≥* 0.1) within single segments and across segment pairs.

The molecular filter addressed compatibility after exchange. Hantavirus nucleocapsid protein binds terminal viral RNA panhandle structures and participates in replication and transcription, making terminal RNA sequence and structure plausible contributors to segment compatibility^24–27^. Cross-segment linkage disequilibrium (LD) measured whether variable nucleotide states on different genome segments occurred together across sampled genomes; this was interpreted as a proxy for segment-background coupling. Across all seven species, branch-length-normalized reassortment origin rates declined strongly with mean cross-segment coding LD (Pearson *r* = *−*0.916*, p* = 0.0037; Spearman *ρ* = *−*0.893*, p* = 0.0068), and this inverse relationship held across the six reassortment-positive species when ANDV was excluded (*r* = *−*0.842*, p* = 0.035; Spearman *ρ* = *−*0.829*, p* = 0.042), indicating that the association is not driven solely by the absence of observed reassortment in ANDV (Supplementary Table 8). Codon-level selection scans and a 10,000-permutation phylogenetic epistasis screen further indicated that this cross-segment LD signal reflected broader segment-background coupling, with no evidence of unusually pervasive site-wise purifying selection or individual amino-acid pair epistasis (Supplementary Fig. 7; Supplementary Table 4). UTR structural divergence measured differences among predicted terminal RNA folds, providing a proxy for compatibility differences in the regulatory regions contacted by viral replication machinery.

Cross-segment LD was highly localized. Variable LD links occupied only 0.15–0.23% of S-segment sites, 0.14–0.27% of M-segment sites and 0.11–0.21% of L-segment sites. Between segments, detectable linkage was concentrated in pairs involving the polymerase segment, spanning both M–L (0.06–0.13% of site pairs) and S–L (0.04–0.05%), whereas S–M links contributed little species-level signal (*≤*0.02%), consistent with cross-segment coupling being anchored through the L segment across both the replication module (S and L) and the glycoprotein (M) (Fig. 2e). Segment-combination diagnostics further supported a non-random compatibility filter. HTNV, PUUV, SEOV, and TULV were dominated by single-segment reassortants, especially M-only, whereas DOBV and SNV were dominated by multi-segment constellations (Fig. 2d; Supplementary Fig. 2). These patterns indicate that ecological opportunity alone is insufficient: retained reassortment also depends on lineage-specific molecular compatibility of the exchanged segment constellation (Fig. 2).

### 2.3 Predictive signal was strong within species but weakly transferable across species

We next used a supervised predictive model suite to test whether ecological, molecular, and lineage-context variables distinguished reassortant from non-reassortant genomes, and whether signals of reassortment were species-specific or generalized across species. Feature coverage is summarized in Supplementary Figs. 5 and 6, and covariate definitions with nonmissing fractions are reported in External Supplementary Table 7. Within the sampled species space, integrated predictors classified retained reassortment accurately, with mean cross-validated AUC = 0.907 (SD = 0.024) and average precision = 0.837 (SD = 0.045) (Fig. 3a). After repeated predictions were collapsed to a single probability per sequence, out-of-fold AUC and average precision were 0.905 and 0.830, respectively (Fig. 3b). At the balanced-accuracy threshold, the model classified 176/553 sequences as reassortant (129/163 true positives [79.1% sensitivity] and 47/390 false positives [87.9% specificity]), closely tracking the observed 163/553 reassortants (Fig. 3c).

**Figure 3:**
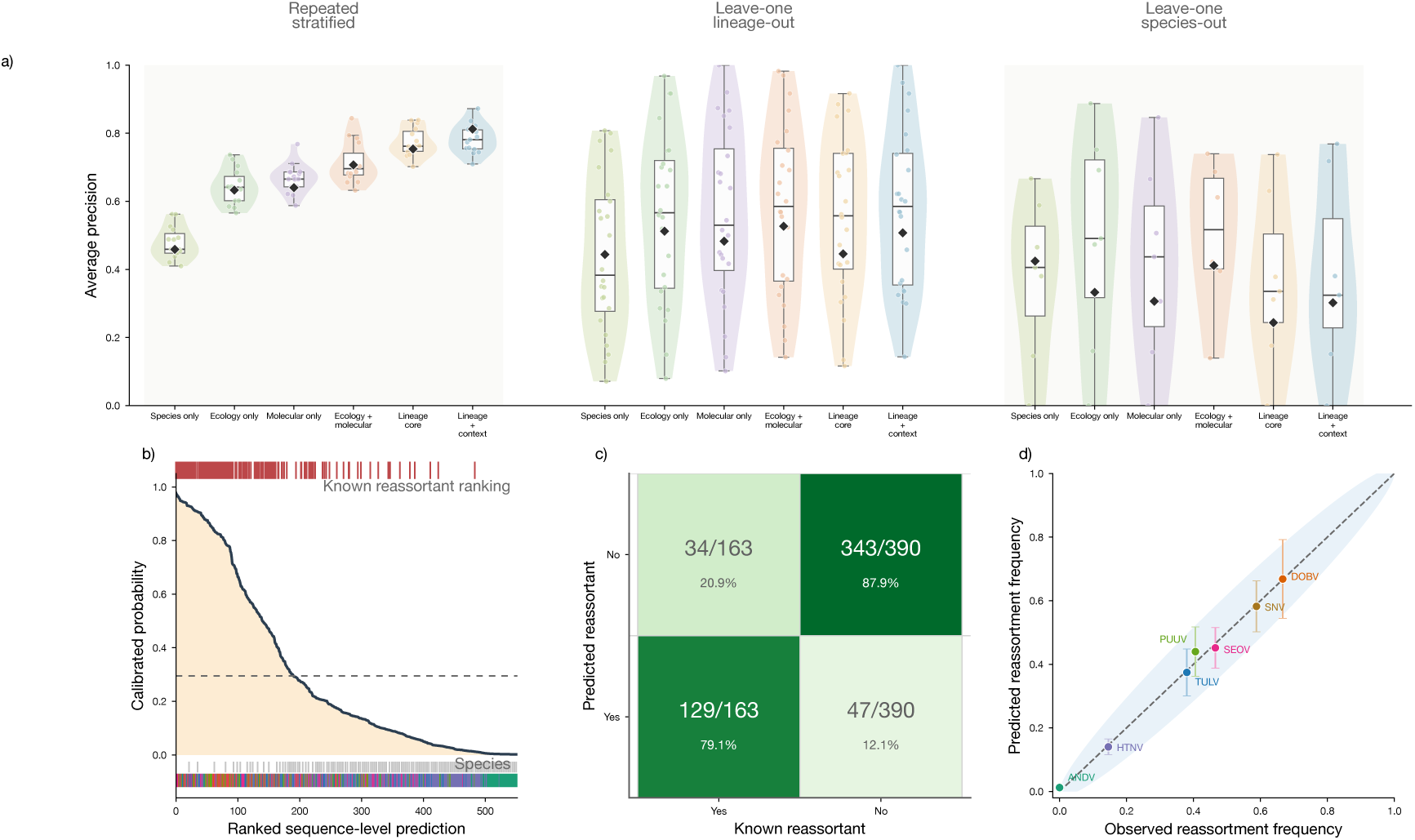
**a,** Out-of-fold average precision across feature sets and validation regimes, comparing repeated-stratified, leave-one-lineage-out, and leave-one-species-out prediction (violins and boxplots show fold distributions; black diamonds mark pooled out-of-fold average precision). **b,** Ranked out-of-fold calibrated reassortment probabilities for sequence-level predictions (*n* = 553, dashed line at the classification threshold), with known reassortants (top red rug) and species identity (bottom color strip) shown as marginal annotations. **c,** Thresholded sequence-level classification counts comparing predicted and known reassortment labels. **d,** Species-level observed versus mean predicted reassortment frequencies (*±*1.96 SE error bars and 95% binomial confidence band around the 1:1 diagonal), showing calibration across the seven focal orthohantavirus species.

Prediction errors were biologically structured. High-reassortment systems such as SEOV, DOBV, and SNV were generally recovered as reassortment-permissive; ANDV remained reassortment-negative, and under stricter specificity thresholds the main underprediction occurred in HTNV (where 29/199 sequences were observed reassortants), consistent with HTNV having a large non-reassortant background and a localized reassortment signal (Fig. 3d).

Predictive signal weakened when models were required to transfer across unsampled lineages within species (pooled average precision *≈* 0.45–0.53) but did not collapse (Fig. 3a; Supplementary Fig. 4). Across-species transfer made ecological and molecular variables substantially weaker predictors of retained reassortment, indicating that retained reassortment is not governed by a single orthohantavirus-wide rule. Instead, species differ in reservoir ecology, segment-compatibility regimes, sampling histories, and segment constellations that persist after exchange. Thus, ecological and molecular variables distinguish reassortant genomes within sampled species, but weak cross-species transfer supports a lineage-conditioned model of effective reassortment (Fig. 3a).

### 2.4 Local host overlap was the clearest ecological correlate of retained reassortment

Sequence-level explanation profiles separated reassortant and non-reassortant genomes across host ecology, molecular structure, tree discordance, and sampling/control features (Fig. 4a). Local host overlap was the clearest ecological correlate of retained reassortment after accounting for baseline differences among virus species (Fig. 4d; Supplementary Table 9). A one-standard-deviation increase in local host-overlap opportunity was associated with higher reassortment probability, with posterior mean log-odds effect 0.297 (odds ratio 1.35) and 94% highest-density interval (HDI) 0.039–0.561 (odds-ratio interval 1.04–1.75) (Fig. 4c). The ecology-only model gave the same qualitative result, with posterior mean 0.289 (odds ratio 1.34), 94% HDI 0.044–0.533, and 98.9% posterior mass above zero. Removing spatial and host-attribution noise strengthened this relationship: excluding sequences georeferenced only to country centroids increased the posterior mean effect of local host overlap to 0.356 (odds ratio 1.43, 94% HDI 0.118–0.622, posterior probability positive = 0.996; *n* = 512), and restricting the analysis to genomes sequenced directly from wildlife reservoirs (excluding human clinical cases) increased the effect to 0.340 (odds ratio 1.40, 94% HDI 0.090–0.642, posterior probability positive = 0.994; *n* = 449). The positive association with local host overlap also persisted after deduplicating repeated sampling to a single genome per locality, year, and host species (posterior mean 0.393, posterior probability positive = 0.950; *n* = 173), collapsing each reassortment origin clade to a single representative tip (posterior mean 0.224 across random draws), or varying the rootward path-distance threshold used to define reassortant tips (*d* = 1: posterior mean 0.285, 94% HDI 0.060–0.526; *d ≤* 3: posterior mean 0.238; Supplementary Table 10).

**Figure 4:**
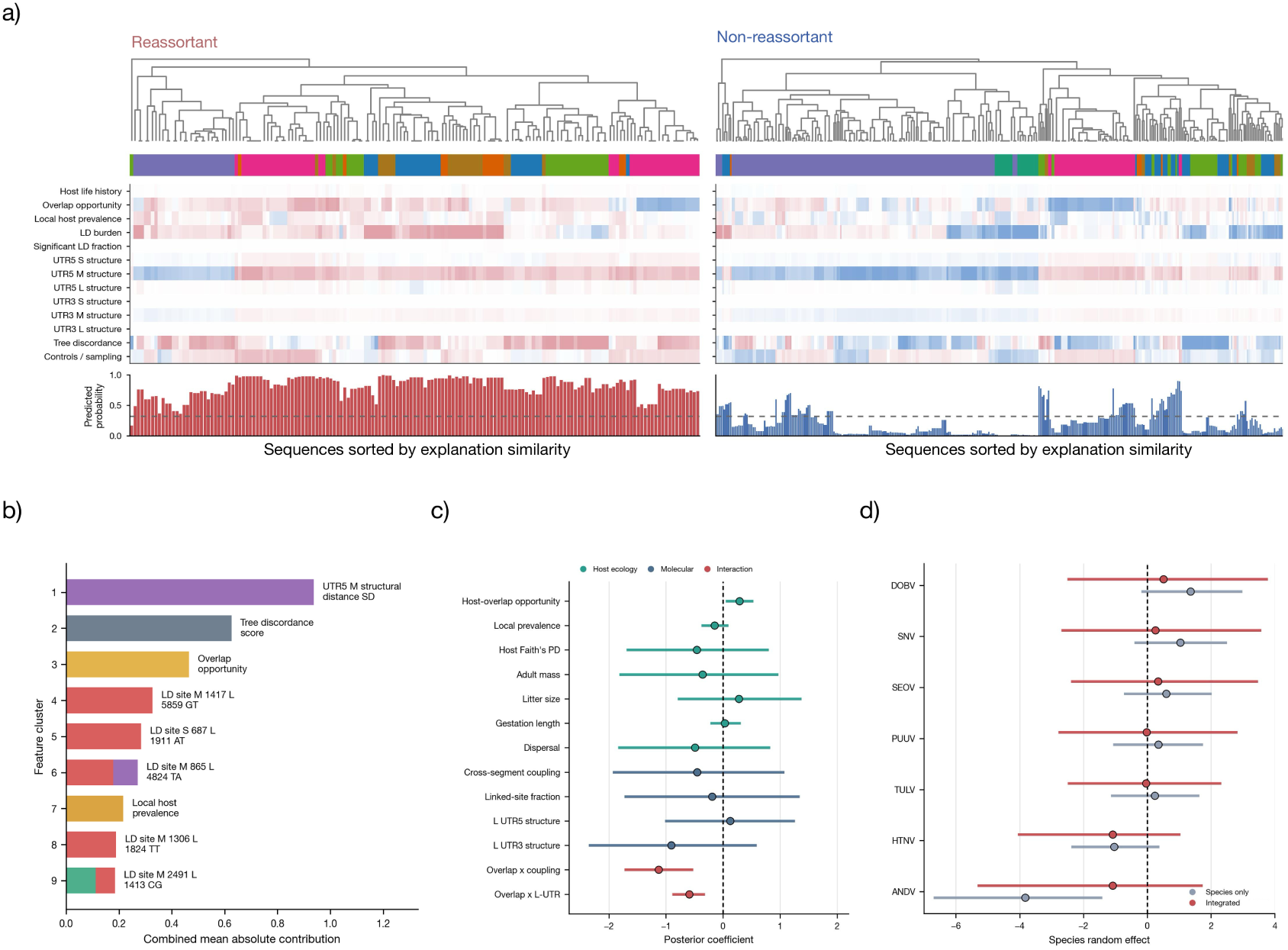
**a,** Sequence-level SHAP explanation profiles (log-odds scale) for reassortant (*n* = 163) and non-reassortant (*n* = 390) genomes, sorted by explanation similarity (top dendrograms) and aligned to predicted reassortment probability (bottom bars). **b,** Ranked feature-contribution clusters from the predictive model, summarized by combined mean absolute SHAP contribution on the log-odds scale. **c,** Bayesian hierarchical posterior regression coefficients (points: posterior means; horizontal bars: 94% highest-density intervals [HDIs] on the log-odds scale) for standardized host ecology, molecular, and ecology-by-molecular interaction terms. **d,** Posterior species random intercepts (means and 94% HDIs on the log-odds scale) from the species-only and integrated Bayesian models, showing how integrated predictors absorb species-level baseline reassortment differences toward zero.

By contrast, local reservoir prevalence did not behave like a simple positive opportunity variable. Its coefficient trended slightly negatively in both ecology-only and integrated models, with 94% HDIs crossing zero across spatial pooling radii from 10 to 1,000 km (Fig. 4c; Supplementary Tables 9 and 10). Thus, retained reassortment was more closely associated with local co-occurrence of plausible reservoir communities than with infection intensity in a single host population. Broad host summaries, including host phylogenetic diversity and life-history traits, were weaker and more uncertain predictors of retained reassortment, suggesting that these variables are too coarse to replace local contact structure.

### 2.5 Molecular compatibility modified the effect of ecological opportunity

Molecular predictors gave a complementary picture. The most interpretable tendency was a negative association between cross-segment coupling and effective reassortment. In the integrated model, cross-segment linkage had a posterior mean log-odds effect of *−*0.456 to *−*0.682 (odds ratio 0.51–0.63), although its 94% HDI remained broad because baseline species differences were absorbed by the species random intercepts; the molecular-only model retained the same negative direction (Fig. 4c; Supplementary Table 9). Feature-contribution summaries ranked terminal-region structure, tree discordance, host-overlap opportunity, and individual LD site states among the strongest explanatory components (Fig. 4b). This is consistent with a compatibility-filter model in which viral backgrounds with stronger pre-existing segment coupling are less tolerant of exchanged segment constellations.

UTR-derived variables were more complex. No individual UTR5 or UTR3 structural-distance main effect had a 94% HDI excluding zero, but incorporating ecological and molecular summaries substantially contracted posterior species-level random intercepts toward zero relative to the species-only baseline, suggesting that terminal-region structure and cross-segment linkage capture part of the between-species molecular background (Fig. 4d; Supplementary Table 9). This effect was not universal, consistent with the expectation that UTR compatibility, nucleoprotein-RNA recognition, packaging constraints, and segment ancestry differ among lineages.

The strongest support for the molecular-filter model came from interactions between ecological opportunity and molecular background. Allowing local host overlap to interact with cross-segment coupling or L-segment 5’ UTR structure improved leave-one-out information criterion (LOOIC) relative to both the species-only and additive models (ΔLOOIC = *−*14.17 and *−*13.49; Supplementary Table 2). In both cases, the interaction term was negative on the log-odds scale, and its 94% HDI excluded zero: *−*1.127 (odds ratio 0.32, 94% HDI *−*1.731 to *−* 0.522; or *−*0.876, odds ratio 0.42, 94% HDI *−*1.280 to *−* 0.472 when LD was estimated within subnational locality cohorts) for host overlap by cross-segment coupling, and *−*0.593 (odds ratio 0.55, 94% HDI *−*0.893 to *−*0.315) for host overlap by L-segment 5’ UTR structure (Fig. 4c; Supplementary Fig. 8; Supplementary Tables 9 and 10). These interactions indicate that host overlap is most predictive in molecular backgrounds where exchanged segments can persist (Fig. 5b). Effective reassortment therefore depends on ecological opportunity filtered through lineage-specific molecular compatibility, consistent with sequential filtering in which their joint occurrence predicts reassortment retention (Fig. 4c).

**Figure 5:**
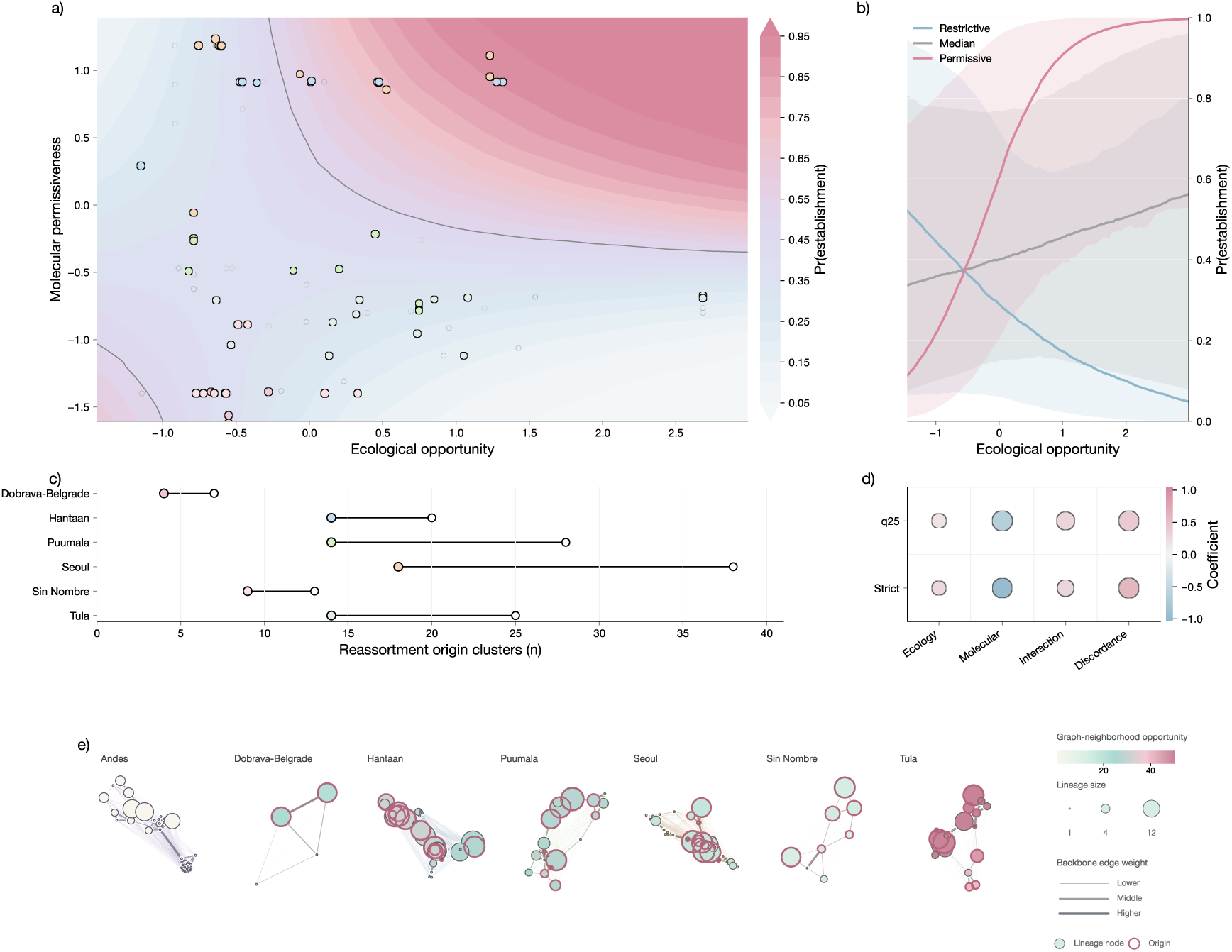
**a,** Posterior mean establishment probability surface across standardized ecological opportunity and molecular permissiveness, with retained minimal reassortment origins (*n* = 73, filled species-colored circles scaled by log descendant clade size) and matched non-reassortant internal clades (*n* = 202, open gray circles) overlaid; solid contour marks Pr = 0.50. **b,** Conditional establishment probability curves (medians and 94% credible ribbons) across ecological opportunity under restrictive (15th percentile), median (50th percentile), and permissive (85th percentile) molecular backgrounds. **c,** Reassortment origin clusters per species before (candidate ARG origin clusters, *n* = 131, open circles) and after (retained minimal origins, *n* = 73, filled circles) topological denesting. **d,** Rarefaction sensitivity of establishment coefficient signs (circle color: mean standardized coefficient; circle size: directional sign stability) under first-quartile and strict minimum species-sampling targets. **e,** Species-specific lineage graphs (143 nodes) showing retained origins (pink borders), lineage size (node area), graph-neighborhood opportunity (node color), and backbone edge weights.

### 2.6 Ecological opportunity and molecular permissiveness jointly predicted reassortment establishment

We next used establishment and expansion models to ask whether retained reassortment origins differed in their probability of evolutionary establishment and descendant-clade growth. We treated ARG-supported reassortment origins as empirical introductions and compared 73 de-nested reassortment origins with 202 matched non-reassortant internal nodes from the same species and nearby lineage context (Fig. 5c, e; Supplementary Fig. 9). The establishment model included ecological opportunity, molecular permissiveness, their interaction, tree discordance, graph-neighborhood opportunity, and sampling effort (Supplementary Table 9).

The interaction between ecological opportunity and molecular permissiveness was the strongest establishment result (all establishment coefficients on the log-odds scale). Ecological opportunity alone was positive but uncertain, with a posterior mean of 0.509 (odds ratio 1.66) and 94% HDI *−*0.334–1.430. Molecular permissiveness alone was also positive but uncertain, with a posterior mean of 0.631 (odds ratio 1.88) and 94% HDI *−*0.577–1.867. By contrast, their interaction was positive and concentrated away from zero, with a posterior mean of 0.866 (odds ratio 2.38), 94% HDI 0.004–1.826, and posterior probability positive 0.966 (Fig. 5a, b; Supplementary Table 9).

Tree discordance was also positive, as expected for ARG-supported origins arising in locally discordant segment histories. Graph-neighborhood opportunity alone was not positive, and sampling effort trended negative after matching and adjustment, indicating that the establishment signal was not driven by better-sampled or more connected lineage neighborhoods.

The expansion component showed weaker support (Fig. 5c; Supplementary Table 9). In the negative-binomial model of descendant clade size (log-rate scale), the ecological-opportunity by molecular-permissiveness interaction remained positive in direction (posterior mean 0.192, 94% HDI *−*0.161–0.520), but its 94% HDI crossed zero. Rarefaction sensitivity supported the establishment result: when descendant clade sizes were downsampled to common species-sampling targets, the establishment interaction remained positive in 86.5% of first-quartile target iterations and 77.5% of strict minimum-sampling iterations (Fig. 5d). Thus, retained reassortment origins were most consistently associated with the joint conditions under which establishment becomes possible, whereas post-establishment expansion was more sampling sensitive.

## 3 Discussion

Segmented RNA viruses can exchange genome segments in a single coinfection event, but those exchanges become evolutionarily consequential only if the resulting constellation persists in host populations. In orthohantaviruses, host association, local reservoir overlap, and segment compatibility vary substantially across species. Our results support a sequential-filter model in which reassortment and effective reassortment are distinct: segment exchange may occur during coinfection, but only some exchanged constellations persist long enough to leave sampled descendants. Across seven orthohantavirus species, retained reassortment ranged from undetected in the analyzed ANDV genomes to common in DOBV, SNV, SEOV, PUUV, and TULV, showing that effective reassortment is a species- and lineage-context-dependent outcome.

This finding fits a broader view of orthohantavirus evolution in which host association is strong but strict codivergence is incomplete. Orthohantaviruses often show reservoir-associated phylogenetic structure, yet host switching, reservoir sharing, geographic overlap, and reassortment have all contributed to their diversification^3, 5, 6, 10^. Natural reassortment has been reported in SNV, DOBV, PUUV, and HTNV, and experimental systems show that coinfection can generate viable reassortants^3, 9, 12, 15, 16, 22, 23^. Our results extend that literature by separating possible reassortment from retained reassortment and identifying which exchanged constellations persist long enough to be sampled.

The ecological results identify host overlap as the clearest opportunity axis. Local host overlap was positively associated with reassortment, an effect that strengthened when analyses were restricted to wildlife reservoir isolates and subnationally georeferenced localities, whereas local prevalence was not. High prevalence within a single reservoir population may increase transmission of a single viral background without increasing the likelihood that divergent segment lineages meet. In that setting, denser sampling can add many closely related genomes to the dataset without adding new opportunities to detect effective reassortment. Effective reassortment instead depends on spatial and temporal overlap among genetically distinct viral lineages within host communities, localities, or contact interfaces where divergent infections can co-occur in the same host species, host population, or spillover bridge. This also explains why broad host traits and host phylogenetic diversity were weaker predictors of retained reassortment. Life-history summaries can describe reservoir biology, but they are several steps removed from the cell-level requirement for mixed infection.

The molecular results indicate a second filter acting after ecological contact. Experimental work in orthohantaviruses has repeatedly shown that reassortment is possible but non-random^3, 22, 23^. In DOBV, PUUV, and ANDV/SNV systems, recovered reassortants often exchanged the M segment while retaining S and L from the same parental background, suggesting stronger coupling among the nucleocapsid, polymerase, and viral RNA terminal structures than between the glycoprotein segment and the replication module. Our segment-combination results support this pattern across HTNV, PUUV, SEOV, and TULV, in which M-only reassortants were common, consistent with M-segment exchange altering glycoprotein-mediated host-interface traits while preserving the S-L replication module. However, DOBV and SNV contained many multi-segment constellations, indicating that S-L pairing is not universal and that compatibility constraints are lineage-specific.

The absence of detected reassortants in the 61 ANDV genomes analyzed here is noteworthy, particularly given recent evidence that ANDV circulates across multiple host taxa, including non-rodent reservoirs such as bats^8, 29^. This expanded host range contrasts with the narrow Sigmodontine-restricted distribution traditionally associated with ANDV. If ANDV maintains ecologically distinct transmission chains across divergent mammalian taxa, the opportunities for within-host coinfection with divergent lineages may be structured differently than in orthohantaviruses with a more overlapping reservoir ecology. Whether this multi-host system limits reassortment opportunities by partitioning distinct viral lineages into separate host species remains largely unexplored, although recent whole-genome surveillance of nine wild Chilean *Oligoryzomys longicaudatus* isolates reports local co-circulation of divergent ANDV lineages and potential reassortment^18^.

Two sampling constraints also inform our ANDV result: primary reconciliation required complete S, M, and L triplets (although auditing partial-genome records identified only one additional S+M-complete *≥*70% ANDV isolate lacking an L segment), and 48 of the 61 complete ANDV genomes derived from human clinical cases (Supplementary Table 3), including clonal outbreak clusters and cell-culture passages (deduplicating passages, duplicates, and outbreak clusters to 17 independent ANDV sensu stricto lineages still yielded 0/17 reassortants; Supplementary Information). However, human-biased sampling alone does not explain the absence of detected reassortment in our ANDV cohort: in SNV, 18 of 34 complete genomes also derived from human clinical cases, yet 20 of 34 carried retained reassortment ancestry, whereas ANDV had none in the 61-genome cohort. ANDV may therefore represent a lineage in which divergent viral backgrounds meet less often in reassortment-relevant host or cellular contexts, or in which exchanged constellations are less likely to persist^30, 31^.

The UTR and LD results should be interpreted in the context of this species-specific compatibility filter. Hantavirus nucleocapsid protein recognizes viral RNA panhandle structures, and terminal RNA sequences are involved in genome recognition, encapsidation, replication, and transcription^24–27, 32^, making segment terminal regions plausible features for compatibility. While cross-segment linkage in natural populations can reflect both epistatic selection and shared clonal genealogy when reassortment is rare, the persistence of a negative relationship between coding linkage and branch-length-normalized origin rates across the six reassortment-positive species, together with the independent constraint imposed by L-segment 5’ UTR structure, supports a functional compatibility filter alongside demographic history. In our data, individual UTR coefficients were not universal additive predictors, but UTR summaries reduced species-level variance and interacted negatively with ecological opportunity. Similarly, localized cross-segment LD suggested that compatibility is maintained among specific segment backgrounds. Overall, these findings indicate that molecular compatibility is a distributed property of the segment constellation across coding and terminal regions.

The weak cross-species transfer highlights the importance of species identity in these analyses. Within species, lineages share reservoir ecology, sampling context, segment ancestry, and compatibility architecture, allowing ecological and molecular features to transfer partially. However, these assumptions do not hold across species. Across the sampled cohorts, ANDV, HTNV, SEOV, SNV, DOBV, PUUV, and TULV span distinct retained-reassortment profiles. In some cohorts, ecological opportunity is broad but strongly filtered; in others, retained reassortment is concentrated in specific lineage neighborhoods; and in others, compatibility is permissive enough to allow multiple segment constellations to persist. Thus, the weak predictive performance across species indicates that effective reassortment is conditioned by lineage context.

The fixation-establishment analysis brings these components together. Following the logic of evolutionary graph theory, the fate of a new variant depends on both its properties and the structured population in which it appears^33^. We used ARG-supported reassortment origins as empirical introductions, lineage graphs as structured ecological opportunity space, and descendant clade size as payoff. The strongest signal was the interaction between ecological opportunity and molecular permissiveness. Retained origins were most strongly associated with the joint occurrence of opportunity and permissiveness.

This result is consistent with the sequential-filter hypothesis. Ecological overlap is associated with where reassortment can be introduced, but introduction alone does not predict evolutionary retention. Molecular permissiveness is associated with whether an exchanged constellation can survive the compatibility filter. Lineage history is associated with the regime in which that event is tested. Effective reassortment in orthohantaviruses may therefore be best understood as a conditional evolutionary outcome: divergent viral genomes must meet, exchange, and remain viable in a lineage background that permits establishment.

Several limitations follow from reconstructing microevolutionary processes from macro-scale sequence and occurrence data^34, 35^. Our host-overlap and prevalence variables are species distribution model (SDM)- and record-derived proxies for the unobserved state underlying reassortment: coinfection of individual hosts or cells with divergent viral lineages. Reassortment detection is also conditional on genomic sampling, because unsampled reassortant lineages might remain invisible and clonal expansions can add many nearly identical genomes without increasing the number of distinct segment-exchange opportunities^36^. Finally, our UTR and LD summaries serve as sequence-level proxies for nucleocapsid-RNA binding, packaging, polymerase, and glycoprotein interactions. Together, these constraints focus the analysis on the ecological and molecular factors that govern retained reassortment in natural populations.

Several microecological processes remain unresolved and should motivate future work. Our study used host distributions, prevalence records, and sequence-derived ancestry, but it could not directly observe mixed infection within individual hosts, within-host viral load dynamics, superinfection exclusion, tissue tropism, transmission timing, or the demographic structure of reservoir populations. Age, sex, reproductive status, territorial behavior, seasonal density pulses, maternal antibodies, aggressive encounters, and transient spillover among sympatric rodents could all influence the probability that divergent lineages coinfect the same animal or cell. Future longitudinal field studies that combine dense mark-recapture sampling, whole-genome sequencing from individual hosts, viral-load quantification, and host community surveillance could test the microecological mechanisms that sit between landscape-level opportunity and retained reassortment. Resolving these filters would strengthen pandemic preparedness by helping predict where divergent lineages can successfully exchange segments, providing a mechanistic basis for anticipating novel orthohantaviruses with altered host ranges or increased potential.

## 4 Methods

### 4.1 Study design and analysis overview

We designed the study to keep reassortment detection separate from reassortment explanation. Reassortment labels were inferred upstream from time-calibrated S, M, and L segment phylogenies reconciled with Espalier^28^, without using the ecological, host, molecular, or lineage predictors later tested as explanatory variables. Downstream analyses then asked which ecological settings, host contexts, molecular compatibility features, and lineage backgrounds were associated with effective reassortment, defined here as segment exchange that persisted long enough to be recovered from reconciled phylogenies and assigned to sampled descendants.

The study tested two linked hypotheses. The ecological-opportunity hypothesis predicts more effective reassortment where host overlap, reservoir co-occurrence, and local transmission context increase the probability of mixed infection. The molecular-filter hypothesis predicts that, even where coinfection is plausible, only some segment constellations persist because reassortment is constrained by cross-segment coadaptation, linkage structure, terminal regulatory architecture, and lineage-specific genome backgrounds. We therefore interpreted all statistical models as models of retained reassortment.

The primary sequence dataset included 553 true-labeled sequence-level observations: 163 reassortant and 390 non-reassortant. These observations came from a harmonized seven-species orthohantavirus dataset spanning ANDV, DOBV, HTNV, PUUV, SEOV, SNV, and TULV. The initial harmonized variant table contained 2,131 candidate records; requiring complete S, M, and L segment representation reduced this set to 638 variants, and root-to-tip interquartile-distance (IQD) molecular-clock filtering^37^ during time calibration produced the final 553 genome packages used for the primary explanatory analyses. We excluded incomplete and unlabeled genome triplets from the main inferential models.

### 4.2 Sequence, variant, and metadata assembly

Virus sequence and metadata tables were assembled from NCBI/GenBank-derived species records and harmonized across segment, variant, host, locality, country, and sampling-year fields. Segment records were reconciled to variant-level S/M/L triplets using normalized variant names and metadata crosswalks. Records were filtered for segment completeness (*≥*70%), unambiguous species assignment, and sufficient metadata to support downstream phylogenetic and ecological annotation. When multiple records represented the same variant-segment combination, the retained record was selected to maximize sequence completeness and consistency with species-specific segment alignments.

The final explanatory dataset was built at the sequence level because the modeling dataset contained both variant-level metadata and segment-specific molecular features. Variant-level true labels were propagated to sequence-level rows only after S, M, and L triplets had been reconciled. This preserved the link between the response variable and phylogenetic reassortment ancestry while allowing segment-specific predictors for discordance, LD, and UTR architecture.

### 4.3 Segment phylogenies and direct reassortment labels

For each focal orthohantavirus species, per-segment FASTA files were harmonized, aligned with MAFFT^38^, and filtered to enforce comparable tip sets across S, M, and L. Segment-specific phylogenies were then estimated under a GTR+G substitution model using IQ-TREE 3^39^ and time-calibrated using TreeTime^37^ with segment-specific substitution rates and root-to-tip clock filtering removing tips that deviated by more than 3 interquartile distances (IQDs) from the temporal regression line in any segment. Common variant coverage was enforced before ancestral graph reconciliation. Local discordance among segment histories was reconciled with Espalier^28^, which attempts to reconcile discordance among segment trees likely arising from phylogenetic error/uncertainty while retaining statistically well-supported discordances likely due to actual reassortment events.

More specifically, Espalier compares two segment-specific trees and produces a maximum agreement forest (MAF) by pruning all subtrees that have internally consistent topologies but are in discordant positions relative to the larger topology of the two segment trees. Each subtree in the MAF therefore represents a potential reassortant lineage. Espalier uses likelihood-ratio testing internally (evaluating RAxML-NG site log-likelihoods under a prior recombination rate of *γ* = 0.0002 and retaining alternative subtree attachments when exp(*ℓ*_alt_ *− ℓ*_ref_) *>* 0.1) to evaluate whether a discordant local history is better supported by a two-tree model than by a single-tree model, and then reconciles the pairwise segment comparisons into a species-level ancestral recombination graph (ARG; Supplementary Figs. 1–2).

True reassortment labels were generated by traversing the reconciled Espalier outputs and identifying sampled tips descending from retained reassortment events across S, M, or L histories (without using any external non-phylogenetic information). For segment-combination diagnostics (Fig. 2d; Supplementary Fig. 2), reassortant tips were classified by the segment(s) carrying discordant ancestry along their reconciled root-to-tip path (S, M, L, S+M, S+L, M+L, or sequential/complex when all three segments accumulated discordant ancestry across sequential nodes). Requiring descent from a reconciled reassortment node is stricter than generic segment-tree discordance, which can arise from deep ancestral events, and yielded binary labels at the variant and sequence levels.

We validated labels with a topology-based reassortment-path analysis. For each sampled tip, we traversed the reconstructed tree path from tip to root and recorded whether a reassortment node was encountered. For tips that encountered such a node, we calculated the node distance to the nearest reassortment ancestor, labeling tips reassortment-positive when the nearest reassortment ancestor lay within *d ≤* 2 internal nodes in the primary analysis (evaluated alongside *d* = 1 and *d ≤* 3 sensitivity thresholds; Fig. 1c; Supplementary Table 8).

Tree-discordance quantities were treated cautiously because they are close to the label-generation process. Tip-level tree discordance was calculated from segment-tree patristic distances. For tip *i*, segment pair (*a, b*), and all other shared tips *j*, we computed the correlation between distance vectors *d_i,j_^a^* and *d_i,j_^b^*. Segment-pair discordance was

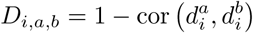

where *d_i_^a^* and *d_i_^b^* are vectors of patristic distances from tip *i* to all other shared tips in segment trees *a* and *b*, respectively, and the final tip-level score was

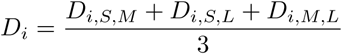

using finite segment-pair values. Because *D_i_* is derived from the same segment phylogenies used by Espalier to infer reassortment ancestry, it was excluded from all primary ecological, molecular, and interaction Bayesian hierarchical models, evaluated only in a dedicated tree-discordance sensitivity model (Supplementary Tables 2 and 9), and included solely as an adjustment covariate in the origin-level establishment and expansion regressions.

### 4.4 Host context, cophylogeny, and ecological opportunity

Host and reservoir context was assembled from ArHa host, sequence, and pathogen tables^8^; harmonized host trait resources including COMBINE^40^, PanTHERIA^41^, and EltonTraits^42^; GBIF-supported taxonomic reconciliation; and branch-length mammalian host phylogenies. Sequence host labels were normalized against ArHa and trait-table names using exact matching, synonym rescue, fuzzy matching, and accepted-name rescue where available. Host phylogenetic diversity was summarized with Faith’s phylogenetic diversity^43^,

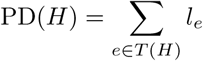

where *T* (*H*) is the minimal subtree connecting the host set *H*, and *l_e_* is the length of branch *e*.

Complementary cophylogenetic analyses placed reassortment in host evolutionary context. Host and virus trees were pruned to shared taxa and compared using PACo and ParaFit-style tests with Cailliez-corrected cophenetic distances and permutation tests where implemented^44–46^. These analyses evaluated whether segment histories followed strict host-virus codivergence or showed segment-specific host-virus mismatch consistent with host switching, reservoir sharing, or reticulate history. Segment-level PACo statistics are reported in Supplementary Table 1 (Supplementary Fig. 10). Cophylogenetic statistics provided evolutionary context for the species comparisons.

Local ecological opportunity was defined at the sequence sampling coordinates (or the subnational locality or country centroid when exact trapping coordinates were unavailable; 512/553 genomes carried subnational coordinates). Local overlap opportunity summarized the predicted sympatric richness of modeled hantavirus reservoir hosts in the 30-arc-second grid cell containing the sequence locality; viral genomes sequenced from human clinical cases (*Homo sapiens*) or carrying unresolved host labels were assigned the predicted small-mammal reservoir overlap of the locality where infection or sampling occurred. Local reservoir prevalence was estimated from ArHa small-mammal surveillance records within a 25-km radius of each sequence’s coordinates (or inverse-distance-weighted across records within 1,000 km when no trap record fell within 25 km, with sensitivity checks across 10–1,000 km radii), matching both the virus and resolved wildlife host species when available or pooling across all small-mammal surveillance records for that virus species when the genome derived from a human clinical case. Host life-history traits were joined from the resolved wildlife reservoir species or imputed to the species-cohort median with a host-provenance indicator for clinical isolates.

### 4.5 Species distribution models and host overlap surfaces

Host-side species distribution models (SDMs) were used to generate macroecological context and host-overlap surfaces. For each host species with sufficient occurrence support, models were fit on a common 30-arc-second global predictor grid using filtered occurrence records, IUCN range polygons buffered by 0.5 degrees, WorldClim bioclimatic variables, and anthropogenic covariates including human footprint, night-time lights, and population density^47–49^. Calibration domains were species-specific and constrained by occurrence and range information.

Within each calibration domain, pseudo-absences were sampled at a 1:1 ratio with presences. Predictors with more than 30% missingness or zero variance were removed, and strong collinearity was pruned at an absolute pairwise correlation threshold of 0.95 while preserving a minimum of 10 predictors when available. Tuned XGBoost and LightGBM classifiers were trained after median imputation, with hyperparameters optimized via Bayesian search with spatial cross-validation, using approximately 150-km blocks where feasible. The two learners were combined as an equal-weight ensemble, and out-of-fold isotonic regression was used to calibrate projected suitability values. Tuned SDM hyperparameters and feature-selection counts are reported in External Supplementary Table 5.

Calibrated host suitability surfaces were projected to the shared grid and stacked across modeled host species. For each grid cell *i*, continuous host-overlap richness was calculated as the sum of calibrated suitability values across species, 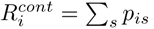, where *p_is_* is the calibrated projected suitability for species s. This metric represents probability-weighted host richness. Thresholded overlap richness was calculated as 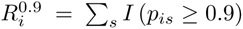, the number of modeled host species with high predicted suitability in that cell. We used 0.9 as the final feature threshold to retain only high-confidence suitable habitat and reduce the influence of diffuse low-suitability predictions. Cells not covered by any projected host surface were treated as missing rather than zero overlap. Sequence localities were linked back to the overlap surfaces to derive local ecological-opportunity variables for virus-side models. For interpretive context, one-dimensional partial-dependence curves were generated for retained predictors across host species, and response direction was summarized with Spearman correlations between predictor values and partial-dependence values.

### 4.6 Molecular compatibility and genomic-filter predictors

The molecular predictor block approximated compatibility constraints acting after segment exchange. Cross-segment linkage disequilibrium was extracted from aligned multi-segment sequence data and summarized by species and segment pair. For pairs of variable sites on different genome segments, contingency tables were built from observed nucleotide states. Cramér’s V was calculated as

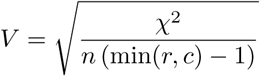

where *χ*^2^ is the Pearson chi-squared statistic, n is the number of genomes, and r and c are the numbers of observed states at the two sites. Normalized mutual information was calculated as mutual information divided by the geometric mean of the two site entropies. Species-level summaries included mean and median Cramér’s V, mean normalized mutual information, number of cross-segment site pairs, and the proportion of site pairs with Cramér’s V *≥* 0.1. We interpreted higher mean cross-segment linkage as a sequence-level proxy for segment-background coupling.

UTR features were included because terminal untranslated regions can affect replication, transcription, packaging, and segment compatibility. Transcript-oriented UTRs were extracted from NCBI/Entrez records using CDS coordinates from GFF3 annotations where available, and GenBank CDS features otherwise. For positive-strand annotations, the 5’ UTR was defined upstream of the primary CDS; for negative-strand annotations, the corresponding downstream sequence was reverse-complemented into transcript orientation. UTRs shorter than 15 nt were excluded. DNA sequences were converted to RNA programmatically. Secondary structures were predicted as minimum-free-energy folds with ViennaRNA RNAfold^50^, and structural dissimilarity between two dot-bracket structures was calculated as base-pair distance, equivalent to the size of the symmetric difference between paired-position sets.

We evaluated two UTR representations. The primary predictive analysis retained direct terminal UTR features, including segment-specific 5’ and 3’ UTR length, recovery, and disparity summaries. Broader species-level 3’ UTR summaries were evaluated in a UTR-augmented sensitivity branch. Because this branch did not improve rank-based discrimination over the simpler lineage-core explanatory bundle, the broader 3’ UTR summaries were excluded from the primary dataset.

### 4.7 Final dataset assembly and preprocessing

The final explanatory table joined direct reassortment labels, sequence metadata, host traits, local ecological variables, ArHa linkage fields, LD summaries, UTR features, segment-discordance summaries, species-level mechanism features, and provenance indicators. Continuous predictors were centered and scaled within the relevant modeling workflow. Unless otherwise stated, standardized predictors were calculated as

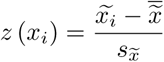

where missing values in *x* were first replaced by the observed median to obtain *x̃*. Missingness indicators were retained when the absence of a value could reflect biological or sampling structure. Categorical predictors were one-hot encoded or handled with model-specific categorical encodings depending on the learner.

Three model-dataset pairs were retained because they answered different biological validation questions. The repeated-stratified predictive analysis tested whether ecological, molecular, lineage, and context variables distinguished reassortant from non-reassortant genomes within the sampled species space. The within-species lineage-transfer model tested whether those signals transferred to unsampled evolutionary neighborhoods within species. The cross-species transfer model tested whether reassortment-associated features generalized to species not seen during training. We interpreted strong repeated-stratified performance, reduced lineage-transfer performance, and weak across-species transfer as evidence that retained reassortment is partly predictable but strongly lineage-conditioned.

### 4.8 Predictive benchmarking and validation regimes

Predictive benchmarking was performed across predefined ecology, molecular, context, control, and integrated feature sets. The model suite included regularized logistic regression, elastic net, group lasso, random forest, XGBoost, LightGBM, and CatBoost^51–54^. Each model was evaluated with preprocessing matched to its requirements, including scaling and imputation for linear models, categorical handling for tree-based models where supported, and fixed feature-block definitions to keep comparisons interpretable. Tree-based learners were included because they can accommodate non-linear ecological effects, categorical LD states, and interactions among host, geography, tree discordance, and molecular compatibility variables without forcing a single linear mechanism.

We evaluated predictive performance under three validation regimes. Repeated stratified cross-validation measured within-dataset predictive signal while preserving overall reassortant/non-reassortant class balance. Within-species lineage-transfer validation held out evolutionary neighborhoods within species, preventing near-neighbor tips from the same local lineage context from appearing in both training and test folds. For this scheme, refined species trees were used to define candidate lineage units within each species. Pairwise patristic distances among tips were clustered with average-linkage hierarchical clustering, and candidate clusters were retained only when they contained at least three tips. Oversized lineage units were split recursively with the same tree-distance procedure, or partitioned when tree-based splitting was not possible, so no held-out group exceeded 30% of the species sample. Candidate units were then packed into held-out evaluation groups; when a species contained both reassortant and non-reassortant observations, packing preferentially formed local, size-constrained groups containing both label classes where possible. Each evaluation group was left out once, and predictions for that lineage neighborhood were generated from models trained on the remaining lineages and species. Across-species transfer validation held out one species at a time and asked whether reassortment signal learned in the remaining species generalized to an unseen species.

Performance was summarized with out-of-fold average precision, ROC AUC, balanced accuracy, Brier score, observed reassortment frequency, and predicted reassortment frequency. Average precision was emphasized in lineage-transfer and species-transfer settings because it measures whether reassortant observations rank above non-reassortants under class imbalance. Probability scores were also thresholded to summarize binary reassortant calls, but thresholded classification was interpreted cautiously because calibration and prevalence differed among species. The validation regimes were not treated as interchangeable: strong repeated-stratified performance but weaker lineage- or species-aware performance was interpreted as evidence that reassortment signal is partly lineage-conditioned.

### 4.9 Feature attribution to prioritize Bayesian covariates

To connect predictive benchmarking with the inferential Bayesian models, we first used SHAP-style feature attribution to identify which variables contributed most strongly to reassortment predictions in the integrated predictive model^55^. Feature contributions were calculated on the logit scale and summarized across observations as mean absolute contributions, both for individual variables and for grouped ecological, host, molecular, tree-discordance, geographic/context, and sampling-control modules.

This attribution step was used to prioritize covariates for the Bayesian models. Variables that repeatedly carried strong predictive signal, including local host-overlap opportunity, cross-segment linkage, terminal UTR structure, tree-discordance measures, and relevant context/control terms, were carried forward into the Bayesian hierarchical framework. The Bayesian models then estimated effect direction, uncertainty, species-level baseline variation, and predeclared ecology-by-molecular interactions for the predictors most supported by the predictive explanation analysis.

### 4.10 Bayesian hierarchical modeling of opportunity and compatibility

The main inferential analysis used Bayesian hierarchical logistic regression with the binary direct reassortment label as the response and species random intercepts for baseline differences among virus species. The Bayesian block-comparison matrix used the 553-observation manuscript sequence cohort and included 163 reassortant positives. For observation i in species *s*[*i*],

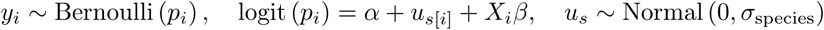

Here, *u_s_* denotes the species-level random intercept and *σ*_species_ is the among-species standard deviation. Models were fit as a predefined block series: species-only baseline, ecology block, ecology interaction block, molecular block, molecular-plus-UTR3 block, additive integrated block, integrated-plus-UTR3 block, tree-discordance sensitivity, and predeclared interaction models. The ecology block included local overlap opportunity, local reservoir prevalence, host phylogenetic diversity, and selected host life-history or trait-availability terms. The molecular block included mean cross-segment Cramér’s V, the significant-pair fraction, and species-level UTR5 structural-distance summaries; UTR3 summaries were added in the UTR3 sensitivity blocks. The additive integrated model combined ecology and molecular predictors to estimate whether the two groups contributed separable signal after accounting for species background. Interaction models tested overlap opportunity by LD burden, overlap opportunity by L-segment UTR5 structural distance, and host phylogenetic diversity by LD burden.

Common effects used weakly regularizing Normal(0, 1) priors, and intercepts used Normal(0, 1.5) priors. The hierarchical block models were sampled with four chains, 3,000 tuning iterations, and 3,000 post-warmup draws per chain, with target acceptance 0.97 unless otherwise stated. Posterior convergence was assessed with R-hat, effective sample size, divergent transitions, and Pareto-k diagnostics for Pareto-smoothed importance-sampling leave-one-out cross-validation (PSIS-LOO). Model fit was compared with leave-one-out expected log predictive density and leave-one-out information criterion, calculated as LOOIC = *−*2 *×* ELPD_LOO_. Species-variance absorption was summarized by comparing posterior *σ*_species_ against the species-only baseline.

### 4.11 Fixation-inspired establishment and expansion analysis

ARG-supported reassortment origins were treated as candidate introductions of reassortment-derived geno-types, lineage graphs represented structured ecological opportunity, and descendant clade size was used as an empirical payoff proxy. The analysis did not simulate a full Moran process because the true infection-to-infection replacement graph is unobserved.

Candidate reassortment-origin clusters were extracted from ARG and local-tree outputs. Because the same reassortment signal can appear at nested ancestral levels, origin calls were de-nested within species. Candidate clusters were sorted by reassortant descendant-tip count, total descendant count, ARG time, and cluster identifier. We retained the smallest ARG-supported origin cluster containing the reassortant tips and removed larger clusters that were strict supersets of an already retained origin. This reduced 131 candidate ARG-origin clusters to 73 retained minimal origins.

For each retained origin, we computed descendant clade size, excess descendant clade size, temporal span, sampled host spread, sampled country and site spread, geographic span, and mean ecological and molecular context. Excess descendant clade size was

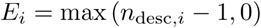

because every detected origin necessarily has at least one sampled descendant.

Matched background observations were sampled from non-reassortant internal S-tree nodes. Eligible background nodes were internal clades with at least two descendant tips and no reassortant descendant tips. Up to three controls were selected for each retained origin from the same species and nearby lineage context, prioritizing shared dominant lineage, similar descendant clade size, similar mean sampling year, similar host and country composition, comparable lineage sampling effort, and comparable terminal UTR recovery. This produced 202 matched non-reassortant background rows.

A species-specific lineage graph represented ecological neighborhood opportunity. Nodes corresponded to species-tree lineage assignments. For lineages a and b, edge weights combined country overlap, host overlap, temporal proximity, and geographic proximity:

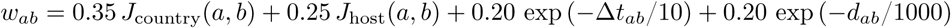

where *J* denotes Jaccard similarity, Δ*t_ab_* is the absolute difference in mean sampling year, and *d_ab_* is the haversine distance in kilometers. For lineage *a*, the graph-neighborhood opportunity was:

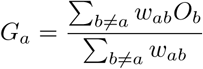

where *O_b_* is mean local overlap opportunity in neighboring lineage *b*. If a lineage had no positive-weight edges, its own overlap score was used. The graph contained 143 lineage nodes and 2,380 weighted edges.

Compact strategy predictors were built from standardized components. Ecological opportunity was

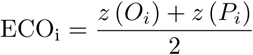

where *O_i_*is the local overlap opportunity and *P_i_* is the local reservoir prevalence.

LD compatibility was the sign-reversed standardized mean Cramér’s V. Species-level UTR compatibility was the sign-reversed mean of standardized UTR5 and UTR3 structural distance. The terminal UTR regime combined lower 5’ disparity and higher 3’ disparity:

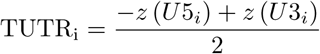

where *U* 5*_i_* and *U* 3*_i_* are the 5’-UTR and 3’-UTR inter-segment length-disparity summaries, respectively. Molecular permissiveness was

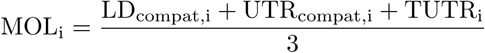

The central sequential-filter predictor was the standardized product

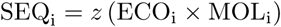

The establishment component was a Bayesian Bernoulli mixed model comparing retained origins with matched background nodes:

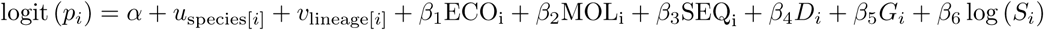

where *D_i_* is tree discordance, *G_i_* is graph-neighborhood opportunity, and *S_i_* is sampling effort. Random intercepts were included for species and dominant lineage.

The expansion component was a Bayesian negative-binomial model fit only to retained origins:

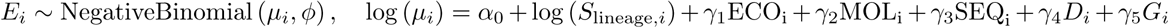

The log lineage sampling effort term was used as an offset. Establishment models used weakly regularizing priors with Normal(0, 1.5) on the intercept and Normal(0, 1) on common effects. The expansion model used Normal(0, 1.5) on the intercept, Normal(0, 0.75) on regression coefficients, and Exponential(1) on the negative-binomial overdispersion parameter. The establishment model was sampled with four chains, 900 tuning iterations, and 900 post-warmup draws per chain; the count model used four chains, 1,600 tuning iterations, and 900 post-warmup draws per chain.

### 4.12 Rarefaction sensitivity for sampling bias

Because establishment and descendant clade size can be affected by uneven sequencing intensity among species, we performed rarefaction sensitivity for the fixation-inspired model. For each retained origin, descendant counts were repeatedly rarefied to common species-level sampling targets using a hypergeometric sampling scheme:

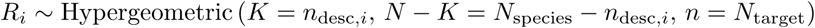

where *N*_species_ is species-level sampling effort and *N*_target_ is the rarefaction target. Two targets were used: the strict minimum species sampling effort and the first-quartile species sampling effort. Origins with fewer than two sampled descendant tips after rarefaction were treated as unobserved under that sampling regime and excluded from the rarefied positive-event set.

For each rarefied dataset, fast regularized sensitivity models estimated coefficient sign stability. The establishment component used L2-regularized logistic regression with balanced class weights, and the expansion component used an L2-regularized Poisson model among retained rarefied origins. The rarefaction summary was the proportion of iterations in which each coefficient retained a positive sign.

### 4.13 Software and reproducibility

Analyses were implemented in Python and R through reproducible repository workflows. Phylogenetic reconciliation and ARG-derived labels were generated with a pipeline based in Espalier. Predictive modeling used scikit-learn-compatible pipelines with XGBoost, LightGBM, CatBoost, random forests, and regularized linear estimators. Bayesian models were fit with PyMC, Bambi, and ArviZ. Graph summaries used NetworkX, and manuscript figures were produced with matplotlib and seaborn^56–61^.

## Declarations

## Acknowledgments

This research was supported by funding to Verena (viralemergence.org) from the U.S. National Science Foundation (NSF DBI 2515340 to S.N.S and D.J.B., supporting D.S. and R.R.). S.N.S. is partly supported by the Centers for Disease Control and Prevention Center for Forecasting and Outbreak Analytics (cooperative agreement CDC-RFA-FT-23-0069). R.R. was partly supported by the Poncin Scholarship Fund from the Poncin Trust under award number GF007810. D.S. receives salary support from the National Science Foundation-National Institutes of Health-National Institutes of Food and Agriculture Ecology and Evolution of Infectious Disease Award (Grant #2208034). This research used resources of the Center for Institutional Research Computing at Washington State University and was supported by the National Science Foundation under Award No. CNS-2216108. We thank all laboratories and researchers who deposited novel Andes virus genome sequences in Pathoplexus. We confirm that we have adhered to the Data Use Terms of Pathoplexus.

## Competing Interests

The authors declare no competing interests.

## Data Availability

Data used in this study is available as described in the methods section and include COMBINE: a coalesced mammal database of intrinsic and extrinsic traits[40], PanTHERIA: a species-level database of life history, ecology, and geography of extant and recently extinct mammals[41], EltonTraits 1.0: Species-level foraging attributes of the world’s birds and mammals[42], and ArHa [8]. Sequence-accession metadata are provided in External Supplementary Table 6 and are accessible through GenBank (https://www.ncbi.nlm.nih.gov/genbank/). Mammal occurence data can be found on GBIF (https://www.gbif.org/), ANDV sequences not available through GenBank can be downloaded from Pathoplexus (https://doi.org/10.62599/PP_SS_2344.1).

## Code Availability

Code to reproduce these analyses is available through GitHub (https://github.com/RicardoRH96/hantavirus-reassortme and Zenodo (https://doi.org/10.5281/zenodo.20573223).

## Author contributions

R.R., D.S., and S.N.S. designed the research; R.R. performed research; D.J.B., I.A.K., S.G., L.D., D.L.W., N.F.M., and D.R. contributed analytical tools; R.R. analyzed data; R.R. and S.N.S prepared the original draft manuscript; all authors revised and approved the final manuscript.

## Supplementary Information

- Supplementary Figs. 1–2 provide ARG-derived reassortment-label diagnostics, including tip-to-root path tracing and segment-combination patterns among reassortant genomes.
- Supplementary Figs. 3–7 provide sampling-discovery, cross-validation calibration, dataset-sensitivity, feature-coverage, and coding-region FEL/linkage-disequilibrium figures.
- Supplementary Figs. 8–10 show Bayesian ecology-by-molecular interaction surfaces, species-specific lineage graph context, and the hantavirus-only host-virus cophylogeny.
- Supplementary Tables 1–4 report segment-level PACo cophylogeny statistics, Bayesian hierarchical LOOIC diagnostics, host-species representation and temporal spans by virus species, and 10,000-permutation epistatic-interaction candidates with FDR, corrected p-values, and statistical-support status.
- External Supplementary Tables 5–7 in supplementary_tables_external.xlsx report host-SDM hyperparameters and feature-selection counts, the final 553-genome S/M/L accession triplets with NCBI submission metadata, and the covariate dictionary with nonmissing fractions.
- Supplementary Tables 8–10 report branch-length-normalized and tip-normalized reassortment origin rates with rootward path-distance breakdowns, complete posterior parameter estimates for all Bayesian hierarchical and establishment/expansion models, and sensitivity analyses across deduplication, host provenance, coordinate precision, and spatial pooling scales.

## Supplementary Information

This supplemental includes supporting figures, supplementary tables, and an external three-sheet workbook. Supplementary Tables 1–4 report segment-level PACo cophylogeny statistics, Bayesian model-comparison diagnostics, host-species representation by virus species, and the 10,000-permutation epistasis interaction table. External Supplementary Tables 5–7 are provided in supplementary_ tables_external.xlsx and report host species distribution model hyperparameters, the final S/M/L accession triplets with NCBI Entrez direct-submission metadata, and a covariate dictionary. Supplementary Tables 8–10 report branch-length-normalized and tip-normalized reassortment rates with rootward path distance threshold breakdowns, complete posterior parameter estimates for all 11 sequence-level Bayesian hierarchical models and origin-level establishment and expansion models, and sensitivity analyses across clonal tip deduplication, outbreak censoring, host provenance, geo-referencing resolution, and spatial interpolation scales.

## Supplementary Methods

### Coding-region FEL selection scans

To assess whether cross-segment linkage disequilibrium (LD) reflected site-wise coding constraint, we ran fixed-effects likelihood (FEL) selection scans on S, M, and L coding alignments for each virus species using HyPhy 2.5^1^ and the FEL framework^2^. Segment alignments and trees were pruned to the common S/M/L isolate set within each species before analysis. Reading frames were selected by minimizing stop codons, and codon columns containing stop codons or more than 50% gaps were excluded.

FEL was run on all branches with synonymous rate variation enabled, no multiple-hit correction, and a nominal site-level threshold of *p ≤* 0.1. Sites were classified as FEL-negative when the nonsynonymous rate estimate was lower than the synonymous rate estimate and the FEL test passed this threshold. Sites were classified as FEL-positive when the nonsynonymous rate estimate was higher than the synonymous rate estimate and passed the same threshold. For the overview summary, site-level q-values were calculated using Benjamini-Hochberg correction^3^. We also mapped model-linked coding LD endpoint sites to their corresponding codons and recorded whether each endpoint fell in a FEL-negative, FEL-positive, or nonsignificant codon.

### Intersegment epistasis screen

We implemented a pragmatic Kryazhimskiy/Neverov-style phylogenetic screen for coordinated substitutions across genome segments^4–6^. This class of methods tests whether substitutions at a trailing site occur unusually soon after substitutions at a leading site on the phylogeny. The rationale is that, under positive epistasis, a mutation at one site can make a subsequent mutation at another site more favorable, producing an excess of rapid consecutive substitutions relative to a permutation null^4^. Neverov and colleagues extended this framework to inter-gene viral evolution with distinct segment histories and highlighted the need to distinguish candidate epistatic clustering from broader episodic selection or hitchhiking effects^5, 6^.

In our adaptation, candidate sites were all segment sites with internal nonsynonymous mutation events, and candidate pairs were all intersegment nonsynonymous site pairs within each virus species. For each ordered segment pair, one segment was treated as the leading segment and the other as the trailing segment. Leading-segment mutation events were projected onto the trailing-segment tree by their shared descendant-tip set, and each lead-trail site pair was scored as the sum of exponentially decayed proximities between projected leading events and descendant trailing events. Two distance modes were evaluated: calendar time, using an exponential scale of 5 years, and branch divergence, using an exponential scale of 0.01 substitutions per site.

For each species, ordered segment pair, and distance mode, empirical pair-level *p*-values were estimated with 10,000 permutations. The permutation scheme preserved the number of events per site while shuffling trailing-segment events across eligible internal branches. Empirical *p*-values were calculated as (*r* +1)*/*(10, 000 +1), where *r* is the number of permuted scores greater than or equal to the observed score. Analysis-level FDR was estimated from 400 permutation draws treated as null datasets, using the expected number of false positives divided by the observed number of nominal positives^7^. The corrected *p*-value reported in Supplementary Table 4 is the permutation-derived probability that a null dataset produced at least as many nominal positive pairs as the observed analysis.

A concrete amino-acid interaction was retained only if it passed all selected-interaction gates: pair-level *p ≤* 0.01, analysis-level FDR *≤* 0.20, hitchhiking-robust positive-fraction *p ≤* 0.10, and at least two non-virtual projected leading events. This gate was designed to distinguish recurrent site-pair clustering from single-event hitchhiking and from broad enrichment that remained consistent with the permutation null.

## Supplementary Results

### FEL scans separate coding constraint from cross-segment linkage

Across 21 species-by-segment FEL scans, 26,788 codons were tested. Of these, 18,341 were classified as FEL-negative and 62 as FEL-positive. FEL-negative fractions were highest in L, intermediate in M, and lowest in S: 11,027/15,241 L codons, 5,629/8,520 M codons, and 1,685/3,027 S codons were FEL-negative (Supplementary Fig. 7). This pattern is consistent with broad purifying constraint across hantavirus coding regions, especially in the polymerase segment.

The FEL results differed from the species-level LD pattern. Andes virus had the highest coding-region LD burden in the model table but the lowest FEL-negative fraction in the common-tip scan: 945/3,830 codons were FEL-negative, compared with 61.1–87.0% in the other virus species. Andes virus also had no FEL-positive sites. Among the 54 coding LD endpoint sites used in the model-linked endpoint test, only 11 were FEL-negative in Andes virus, compared with 37–49 in the other virus species, and no model-linked endpoint was FEL-positive in any species. These results indicate that Andes virus preserves linked coding combinations without showing an unusually high fraction of individually FEL-negative codons. This interpretation remains conservative because Andes virus had fewer variable codons than the other species, reducing power to detect site-level selection.

### Nominal epistasis candidates were not supported after correction

The full intersegment epistasis screen evaluated 215,995 intersegment nonsynonymous candidate pairs, producing 86,900 mutation-event rows and 1,295,970 ordered pair-mode scores. At the primary *α* = 0.01 threshold, 1,535 nominal positives were observed across ordered species-segment-mode analyses, but the permutation-estimated expected false-positive count was 1,740.1. At *α* = 0.05, 13,265 nominal positives were observed, with 13,950.7 expected false positives. These totals indicate that most nominal pair-level signals were consistent with the simultaneous null.

The strongest nominal analyses did not pass the selected-interaction gate. The closest cases were Tula virus L-to-S in divergence and time modes, with 21 and 22 observed positives, analysis-level FDR values of 0.321 and 0.324, and corrected *p*-values of 0.077 and 0.095, respectively. These analyses passed the hitchhiking-robust positive-fraction screen but failed the prespecified FDR threshold. The next-lowest FDR analysis, Dobrava-Belgrade virus S-to-L in divergence mode, had one nominal positive pair and an FDR of 0.235, also above the selected-interaction threshold. Therefore, no concrete amino-acid pair received statistical support under the 10,000-permutation selected-interaction gate.

Accordingly, the final epistasis feature dataset kept species-level permutation summaries and numeric selected-pair burden summaries, but no selected amino-acid or nucleotide pair-state columns. Supplementary Table 4 reports the strongest nominal candidates to document why they were discarded after FDR and corrected-*p* evaluation.

### Sensitivity to phylogenetic depth, clonal expansion, path-distance thresholds, host provenance, and spatial scale

We evaluated whether species differences in retained reassortment and the estimated effects of ecological opportunity and molecular compatibility were robust to differences in phylogenetic depth, post-origin clonal expansion, rootward path-distance cutoffs, clinical versus wildlife host provenance, coordinate precision, and spatial pooling scales across the 553-genome dataset (Supplementary Tables 8–10):

1. **Phylogenetic branch-length and origin-level normalization (Supplementary Table 8):** Dividing de-nested minimal reassortment origins (*n* = 73) by mean segment total branch length (*L̄*_tot_, in substitutions per site) accounted for differences in evolutionary time sampled and clonal expansion across species, yielding origin rates of 0.00 origins per substitution per site in Andes virus (*L̄*_tot_ = 0.30, one-sided 95% Poisson upper bound 10.15), 4.52 in Dobrava-Belgrade virus (*L̄*_tot_ = 0.88), 5.55 in Sin Nombre virus (*L̄*_tot_ = 1.62), 7.26 in Tula virus (*L̄*_tot_ = 1.93), 8.31 in Hantaan virus (*L̄*_tot_ = 1.69), 9.08 in Puumala virus (*L̄*_tot_ = 1.54), and 15.37 in Seoul virus (*L̄*_tot_ = 1.17). Across all seven species, branch-length-normalized origin rates correlated negatively with species-level coding linkage disequilibrium (Pearson *r* = *−*0.916*, p* = 0.0037; Spearman *ρ* = *−*0.893*, p* = 0.0068), and this inverse relationship held across the six reassortment-positive species when Andes virus was excluded (*r* = *−*0.842*, p* = 0.035; Spearman *ρ* = *−*0.829*, p* = 0.042). Among the six reassortment-positive species, sample size *n* correlated negatively with both tip-level reassortment frequency (*r* = *−*0.930*, p* = 0.0071) and origins per tip (*r* = *−*0.957*, p* = 0.0027), indicating that higher reassortment frequencies were not driven by denser sampling.
2. 2. **Andes virus metadata curation, outbreak and passage deduplication, and segment-completeness check:** Because the 61-genome Andes virus cohort included clonal human-to-human transmission clusters (28 Epuyen 2018–2019 cases, three Argentina 2014 cases, and four MV Hondius 2026 travel-associated cases), serial cell-culture and hamster passages, re-sequenced reference isolates, a RefSeq duplicate (NC_003466–68 duplicating AF291702–04), and GenBank records lacking a collection_date qualifier, we verified sensitivity after correcting collection dates, removing passages, duplicates, and the second longitudinal semen sample (ANDV-CH-LS-2022), and collapsing each transmission cluster to a single representative (*n* = 19 independent natural lineages, or *n* = 17 ANDV sensu stricto after excluding two *∼*16–20% divergent lineages, Arg-h484 and Brazil/VARS-22-01). All 17/17 ANDV sensu stricto genomes passed TreeTime 3-IQD molecular-clock filtering across S, M, and L, and Espalier again inferred 0/17 reassortants (reconciled SPR = 0 and RF = 0 across all three segment pairs). Mean cross-segment coding Cramér’s *V* remained high after deduplication (0.483 at *n* = 19 and 0.369 at *n* = 17, compared with 0.430 across all 61 genomes). Auditing the candidate harmonized dataset also identified only one additional S+M-complete (*≥*70%) Andes virus isolate lacking an L segment, indicating that requiring all three segments did not exclude a substantial pool of partially sequenced Andes virus reassortants.
3. 3. **Rootward path-distance threshold sensitivity (Supplementary Tables 8 and 10):** Varying the maximum rootward internal-node distance threshold used to define retained reassortment from *d* = 1 (immediate child of a reassortment node; 113/553 = 20.4% positive) to the primary *d ≤* 2 threshold (allowing at most one intervening coalescent bifurcation; 163/553 = 29.5% positive) and a relaxed *d ≤* 3 threshold (193/553 = 34.9% positive) preserved the positive association with local host overlap (*β* = 0.285, Pr(*β >* 0) = 0.989 at *d* = 1; *β* = 0.305, Pr(*β >* 0) = 0.989 at *d ≤* 2; *β* = 0.238, Pr(*β >* 0) = 0.973 at *d ≤* 3) and the negative direction of cross-segment linkage (*β* = *−*0.518*, −*0.647, and *−*0.557, respectively). Non-reassortant tips that encountered ancestral reassortment nodes deeper in the phylogeny (*d >* 3; 241/553 = 43.6%) were separated from those historical events by a median of 7.0 intervening coalescent nodes (interquartile range 5.0–12.0), whereas 119/553 tips (21.5%, including all 61 Andes virus tips and all 14 non-reassortant Sin Nombre virus tips) encountered no ancestral reassortment node anywhere along their root-to-tip path (*d* = *∞*).
4. 4. **Clonal clade collapse and outbreak deduplication (Supplementary Table 10):** Collapsing each reassortment origin clade to a single randomly sampled representative tip across eight independent draws (*n* = 494, 104 positives; mean *β*_overlap_ = 0.224, SD = 0.033, positive in 8/8 draws; mean *β*_LD_ = *−*0.661, SD = 0.030), selecting a single deterministic representative tip per origin (*β*_overlap_ = 0.268, Pr(*β >* 0) = 0.946), deduplicating repeated local sampling to a single genome per locality, year, and host species (*n* = 173, 61 positives; *β*_overlap_ = 0.393, Pr(*β >* 0) = 0.950), and capping lineage-by-response cluster size at five tips (*n* = 476, 150 positives; *β*_overlap_ = 0.198, Pr(*β >* 0) = 0.964) all preserved the positive effect of local host overlap and the negative effect of cross-segment linkage.
5. 5. **Wildlife reservoir versus clinical host provenance and coordinate resolution (Supplementary Table 10):** Restricting the hierarchical regression to genomes sequenced directly from wildlife reservoirs (*n* = 449, 146 positives) increased the positive effect of local host overlap (*β* = 0.340, 94% HDI 0.090–0.642, Pr(*β >* 0) = 0.994), as did excluding sequences georeferenced only to country centroids (*n* = 512, 151 positives; *β* = 0.356, 94% HDI 0.118–0.622, Pr(*β >* 0) = 0.996) or including a binary covariate for human clinical host provenance (*β*_overlap_ = 0.293, Pr(*β >* 0) = 0.986). Similarly, when cross-segment coding linkage and terminal UTR summaries were estimated within subnational locality cohorts, the negative interactions between local host overlap and cross-segment linkage (*β* = *−*0.876, 94% HDI *−*1.280 to *−* 0.472, Pr(*β <* 0) = 1.000) and between local host overlap and L-segment 5’ UTR structural divergence (*β* = *−*0.591, 94% HDI *−*0.881 to *−* 0.299, Pr(*β <* 0) = 1.000) remained strongly supported.
6. 6. **Habitat suitability thresholds and local prevalence spatial pooling scales (Supplementary Table 10):** Across 36,576 small-mammal capture sites in the SDM evaluation grid, stacked reservoir richness at the primary 0.90 suitability threshold 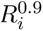 correlated strongly with richness at thresholds of 0.80 (*r* = 0.851) and 0.70 (*r* = 0.806) as well as unthresholded probability-weighted continuous richness 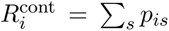 (*r* = 0.784, Spearman *ρ* = 0.895*, p <* 10^−100^). Varying the local prevalence exact-pooling radius (10, 25, or 50 km), maximum inverse-distance-weighted fallback radius (250, 500, or 1,000 km), and distance-decay power (1 or 2) consistently yielded near-zero prevalence coefficients (*β*_prev_ *∈* [*−*0.177*, −*0.008]) while leaving the positive effect of local host overlap unchanged (*β*_overlap_*∈* [0.276, 0.306], all Pr(*β >* 0) *≥* 0.986). Log-transforming 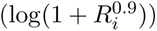 or winsorizing local host overlap at the 90th percentile likewise preserved the positive overlap effect (*β* = 0.248, Pr(*β >* 0) = 0.976 and *β* = 0.309, Pr(*β >* 0) = 0.992, respectively).

---

**Supplementary Fig. 1:**
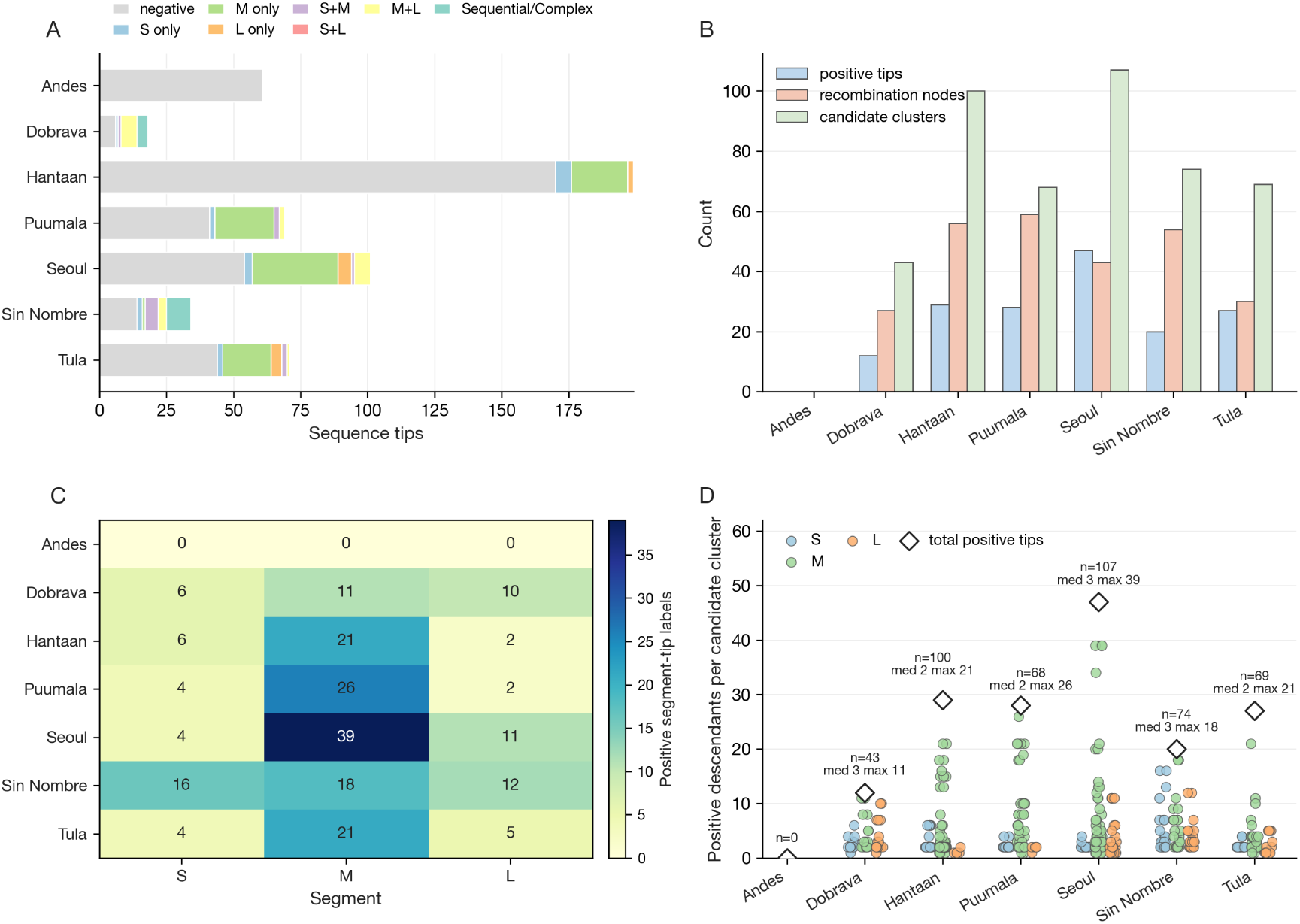
Tip-to-root diagnostics for phylogenetic reassortment labels. A, final tip labels by species and segment combination; B, positive tips, internal reassortment nodes, and candidate descendant clusters; C, segment-specific positive labels; D, positive descendants per candidate cluster.

**Supplementary Fig. 2:**
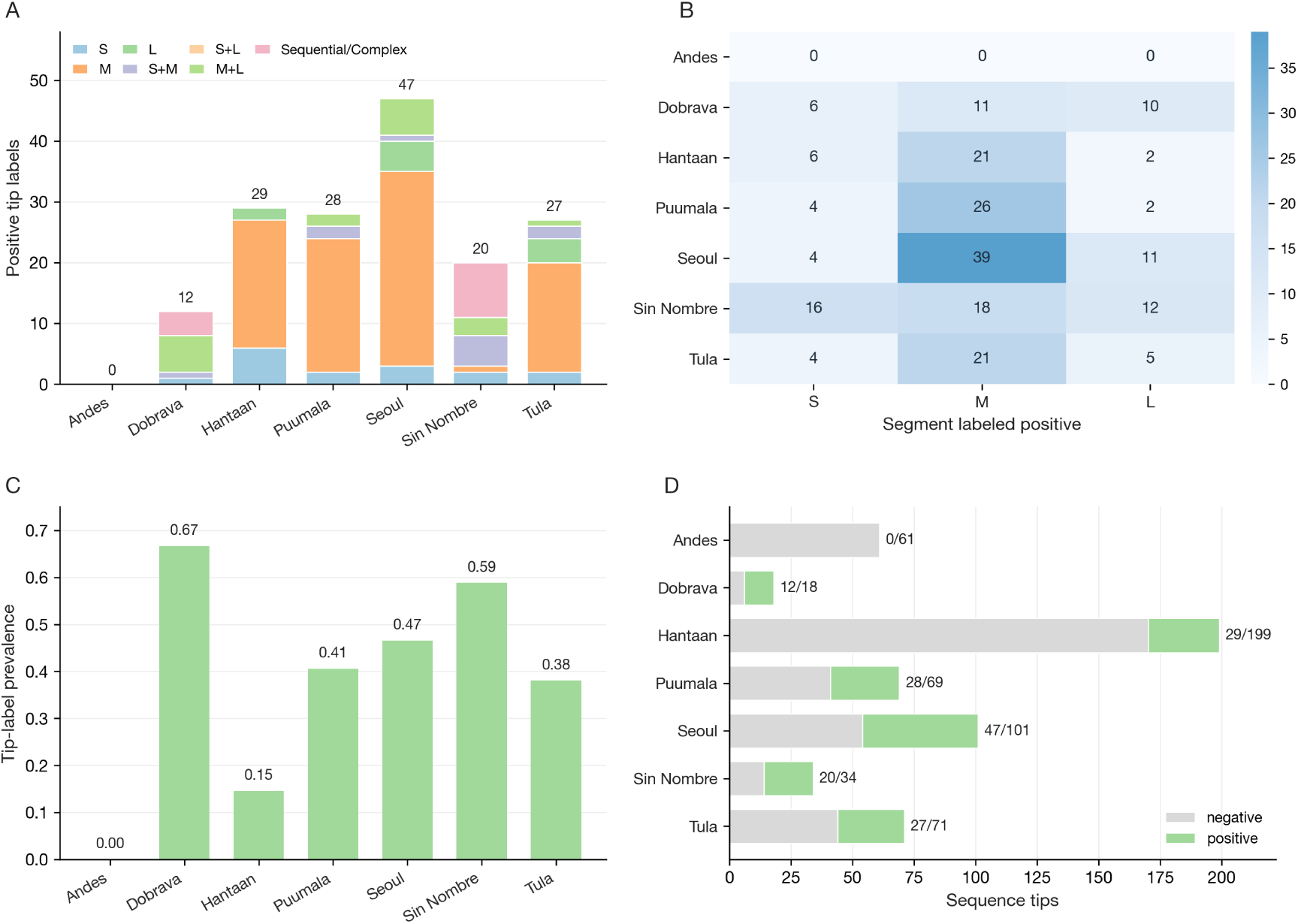
Segment-combination diagnostics for genomes labeled as reassortant. A, positive tips partitioned by inferred S, M, and L segment combinations; B, segment-wise positive-label counts; C, positive-label prevalence by species; D, full labeled-tip counts split by reassortant status.

**Supplementary Fig. 3:**
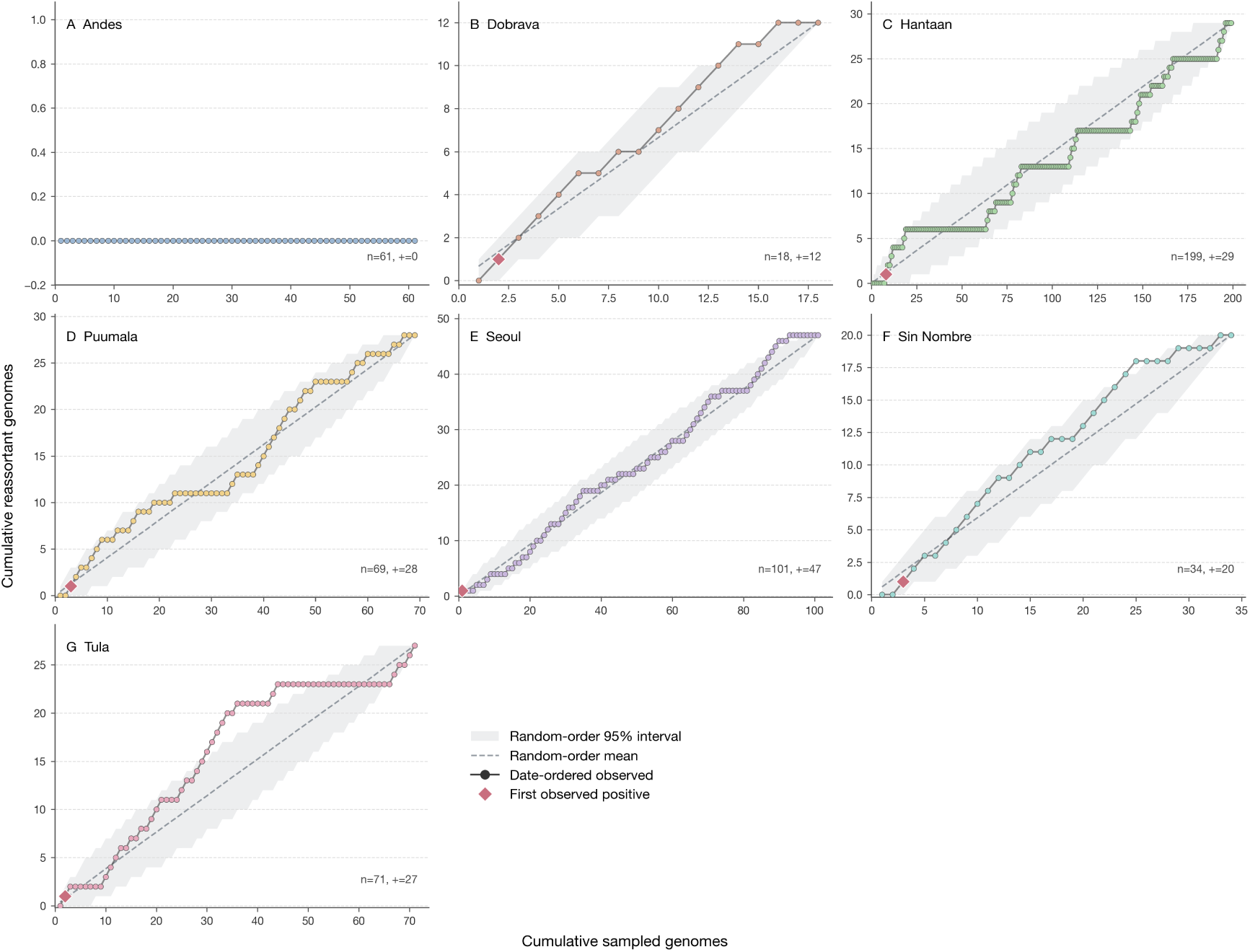
Date-ordered sampling-discovery curves for the final genome set. Species panels show cumulative reassortant labels after ordering genomes by sampling date; shaded envelopes and dashed lines show random-order expectations, and diamonds mark the first positive genome.

**Supplementary Fig. 4:**
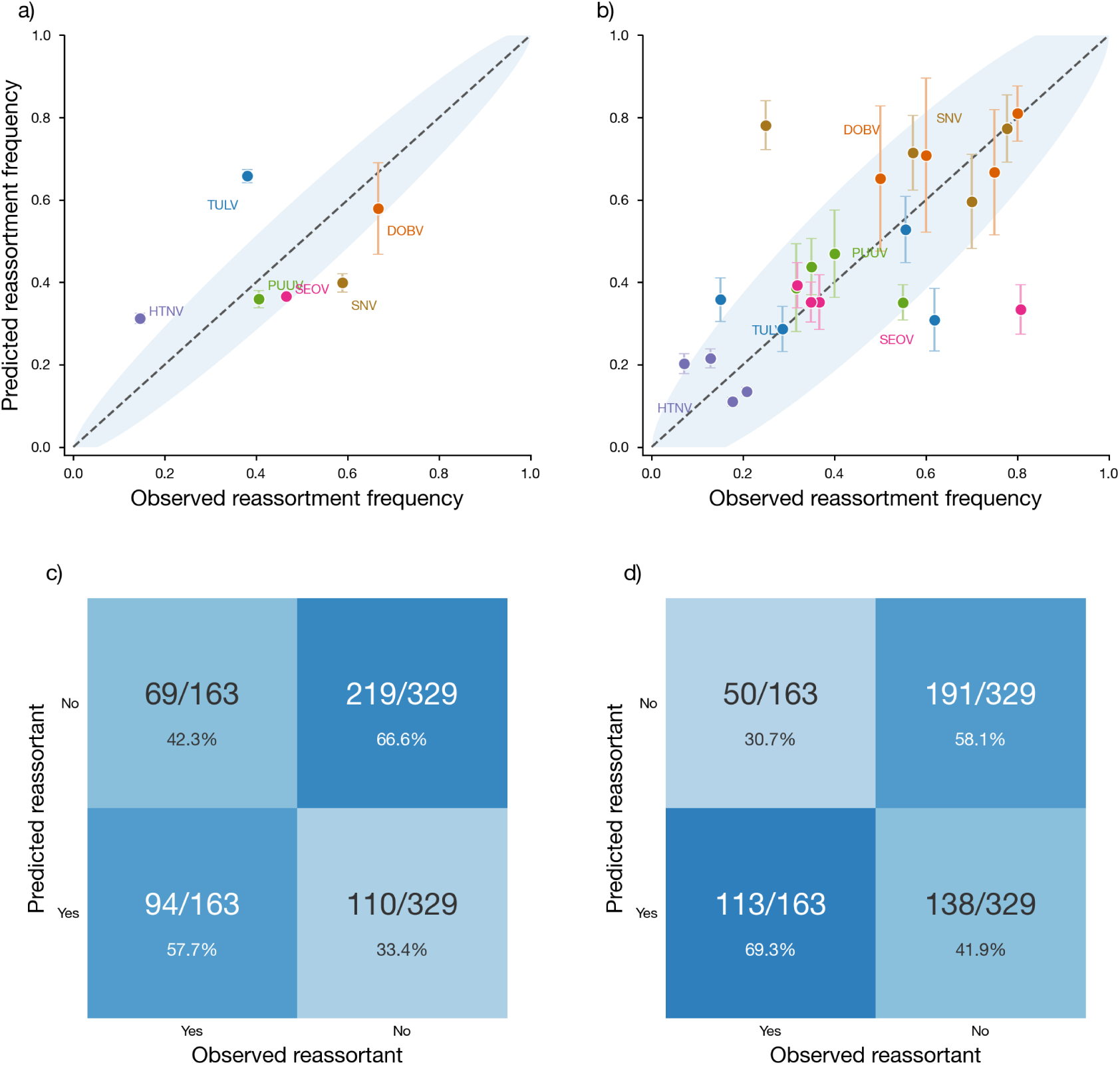
Observed versus predicted reassortment frequencies in the validation analysis. Each point shows the prediction made by the final model after being trained with two different validation schemes. **a)** leave-one-species-out validation. **b)** leave-one-clade-out validation. In both panels, the dashed line shows perfect agreement; the envelope shows the 95% confidence interval for a binomial distribution with the observed frequency as the mean and the number of genomes in the validation set as the number of trials; and points are colored by virus species, with error bars. **c)** and **d)** show the same validation results but expressed as a confusion matrix.

**Supplementary Fig. 5:**
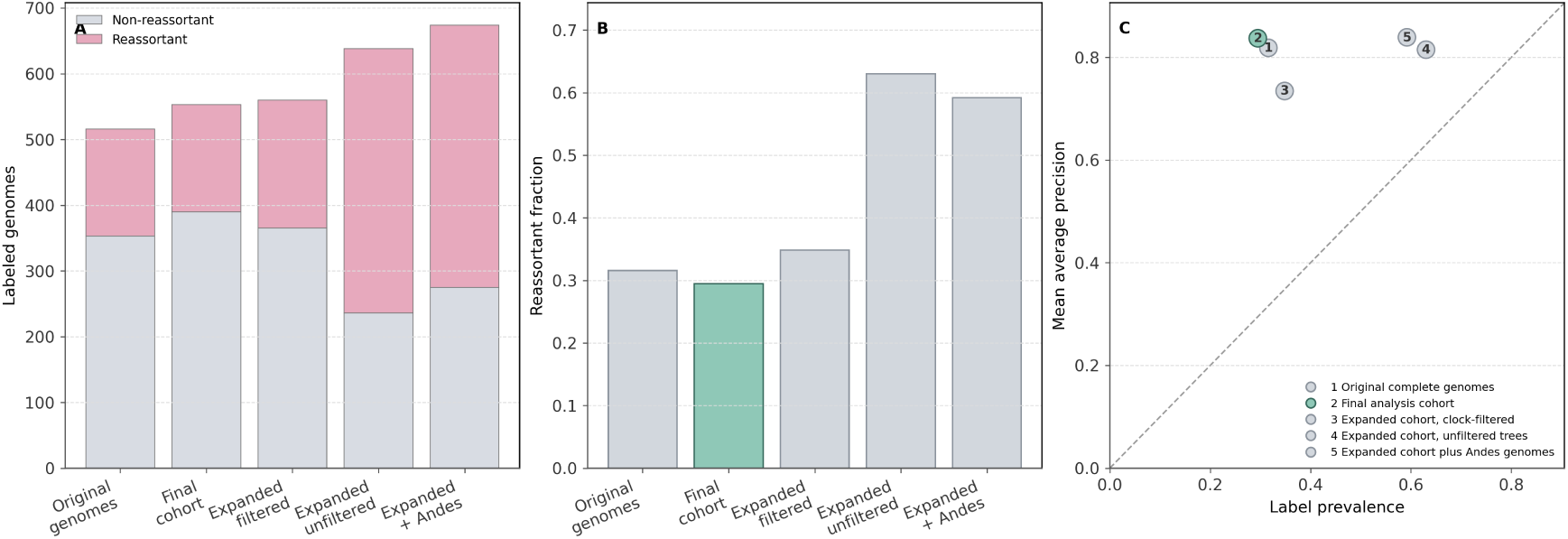
Sensitivity of reassortment-label distribution and predictive performance to genome-sampling scope. A, labeled genomes in each analysis cohort separated by reassortant status; B, reassortant-label prevalence; C, mean average precision plotted against prevalence as the class-balance baseline.

**Supplementary Fig. 6:**
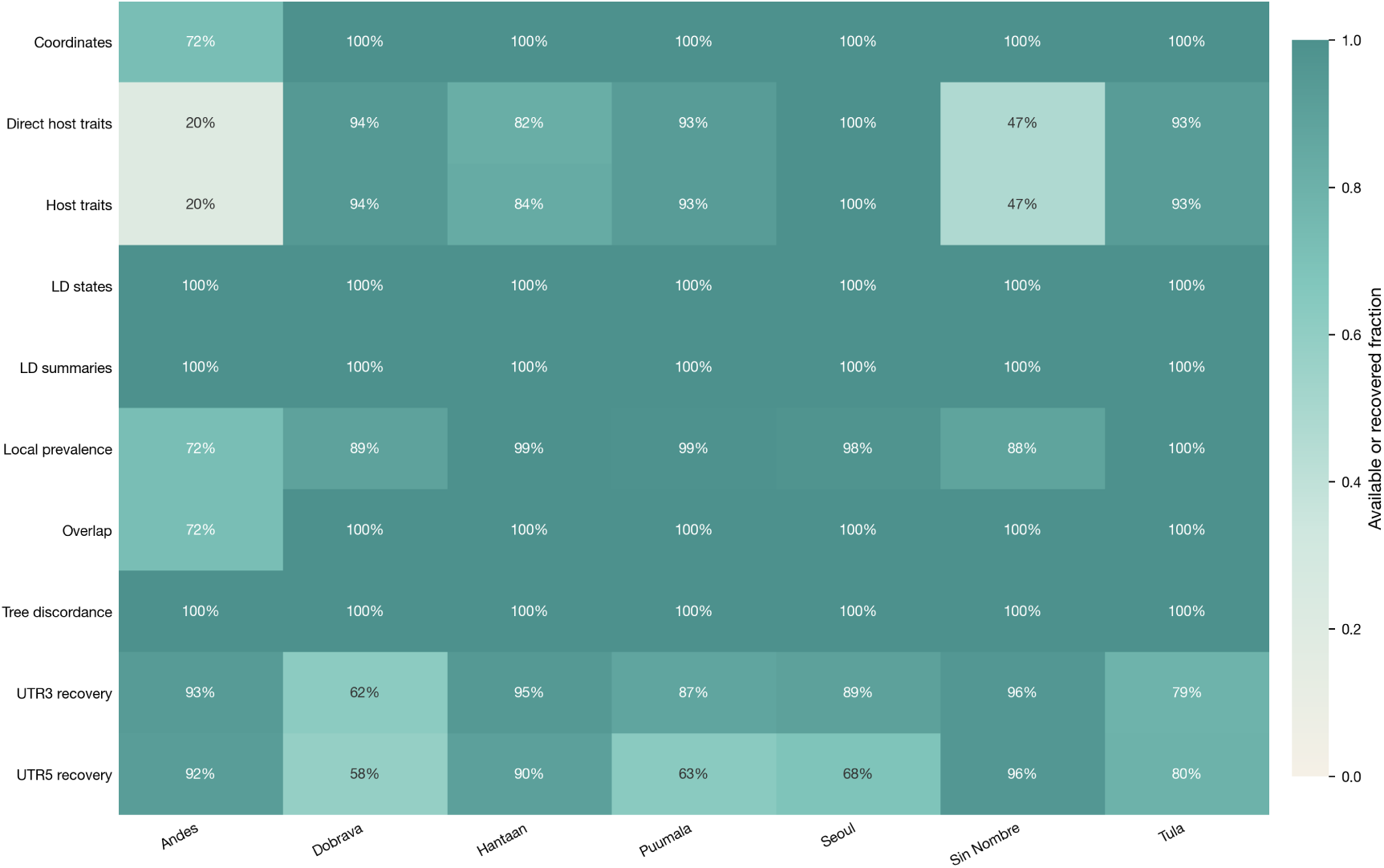
Feature-coverage for the final analysis cohort. Cells show the fraction of genomes per virus species with available or recovered coordinate, local-ecology, host-trait, tree-discordance, linkage-disequilibrium, and UTR features.

**Supplementary Fig. 7:**
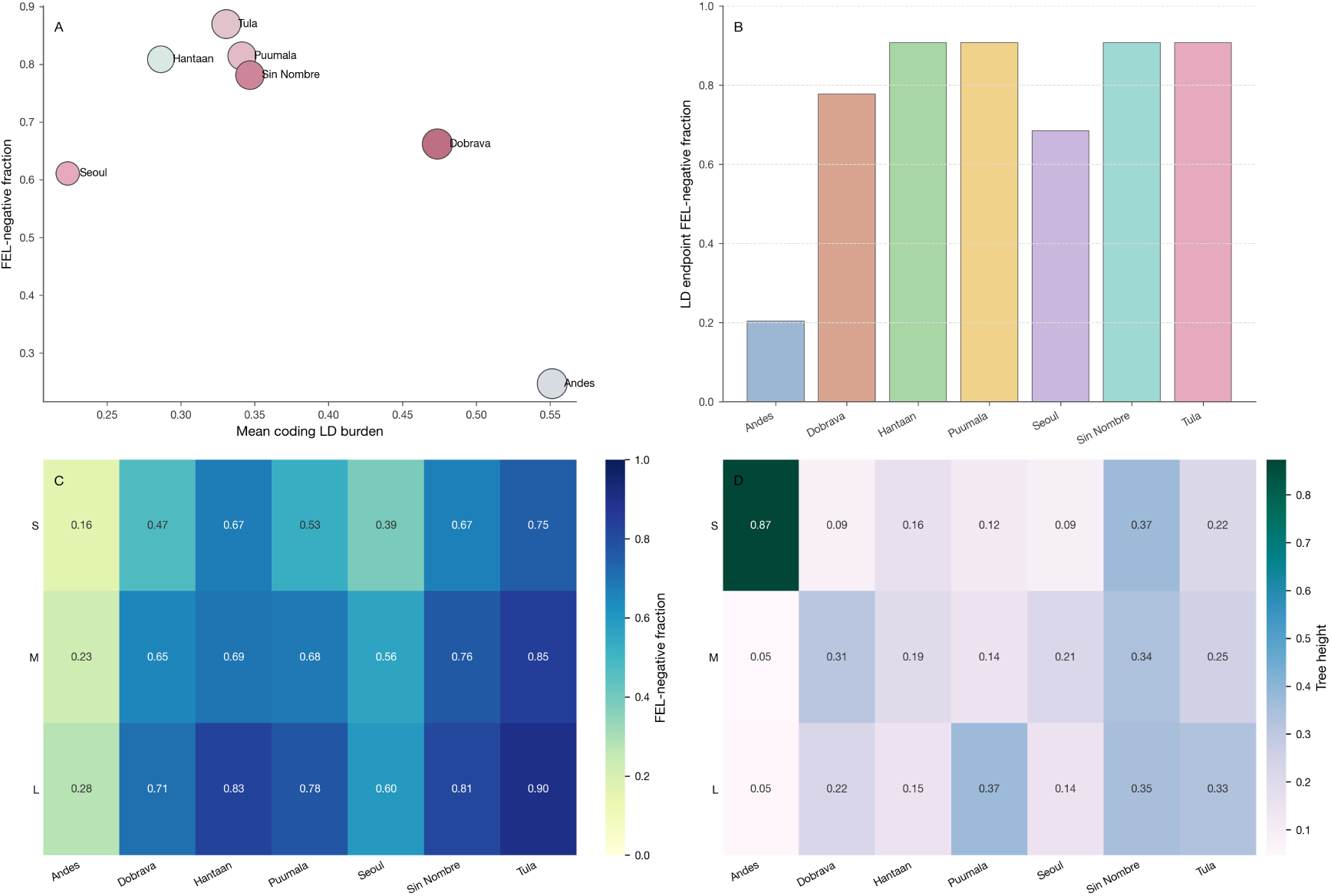
Coding-region FEL and linkage-disequilibrium. A, mean coding LD burden versus FEL-negative fraction by species; B, FEL-negative fraction among model-linked LD endpoints; C, segment-by-species FEL-negative fractions; D, segment-tree heights.

**Supplementary Fig. 8:**
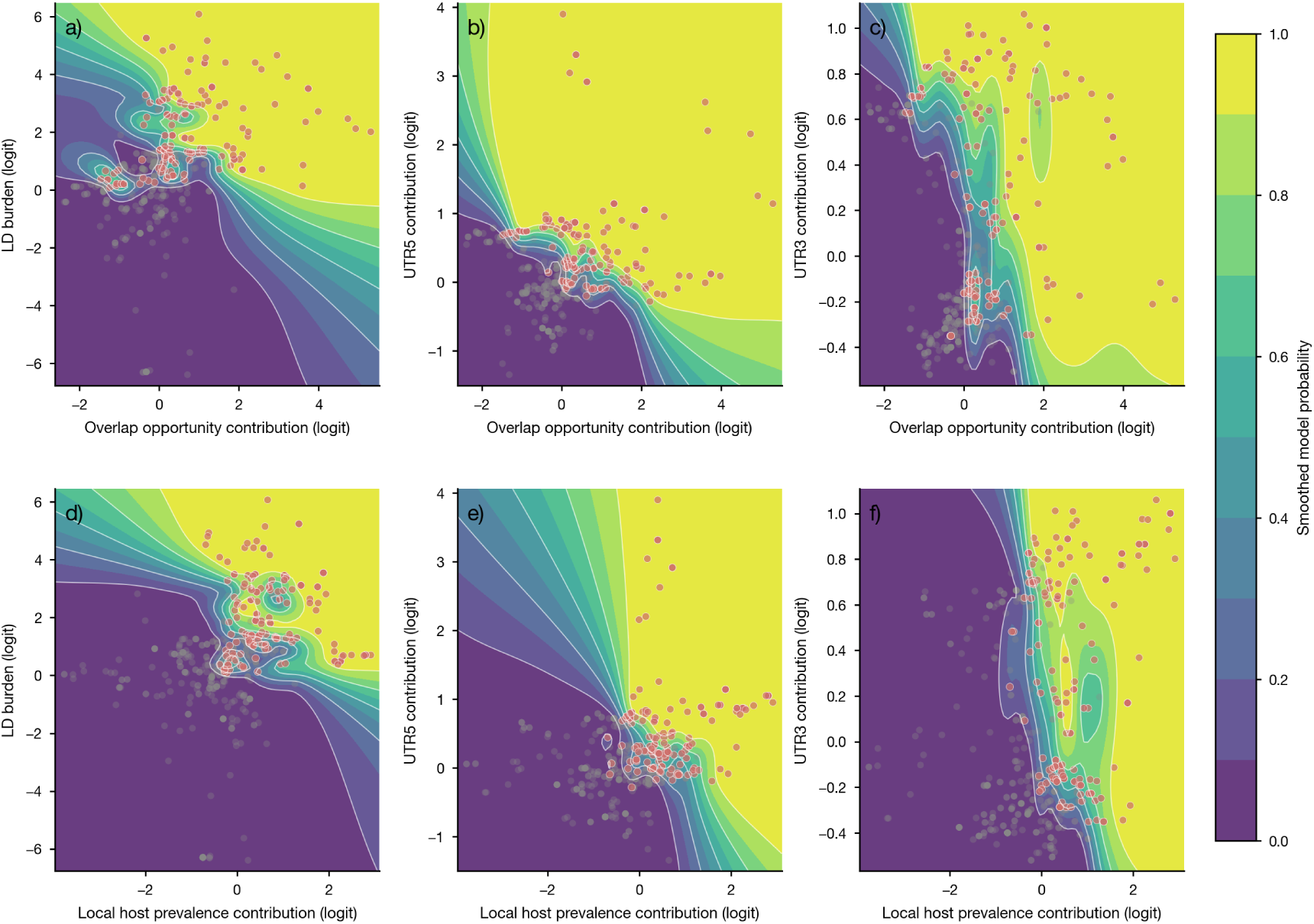
Bayesian interaction surfaces for ecological opportunity and molecular context in the final analysis cohort. A–C, local host-overlap contribution crossed with LD burden, UTR5, and UTR3 contributions; D–F, local host-prevalence contribution crossed with the same molecular predictors.

**Supplementary Fig. 9:**
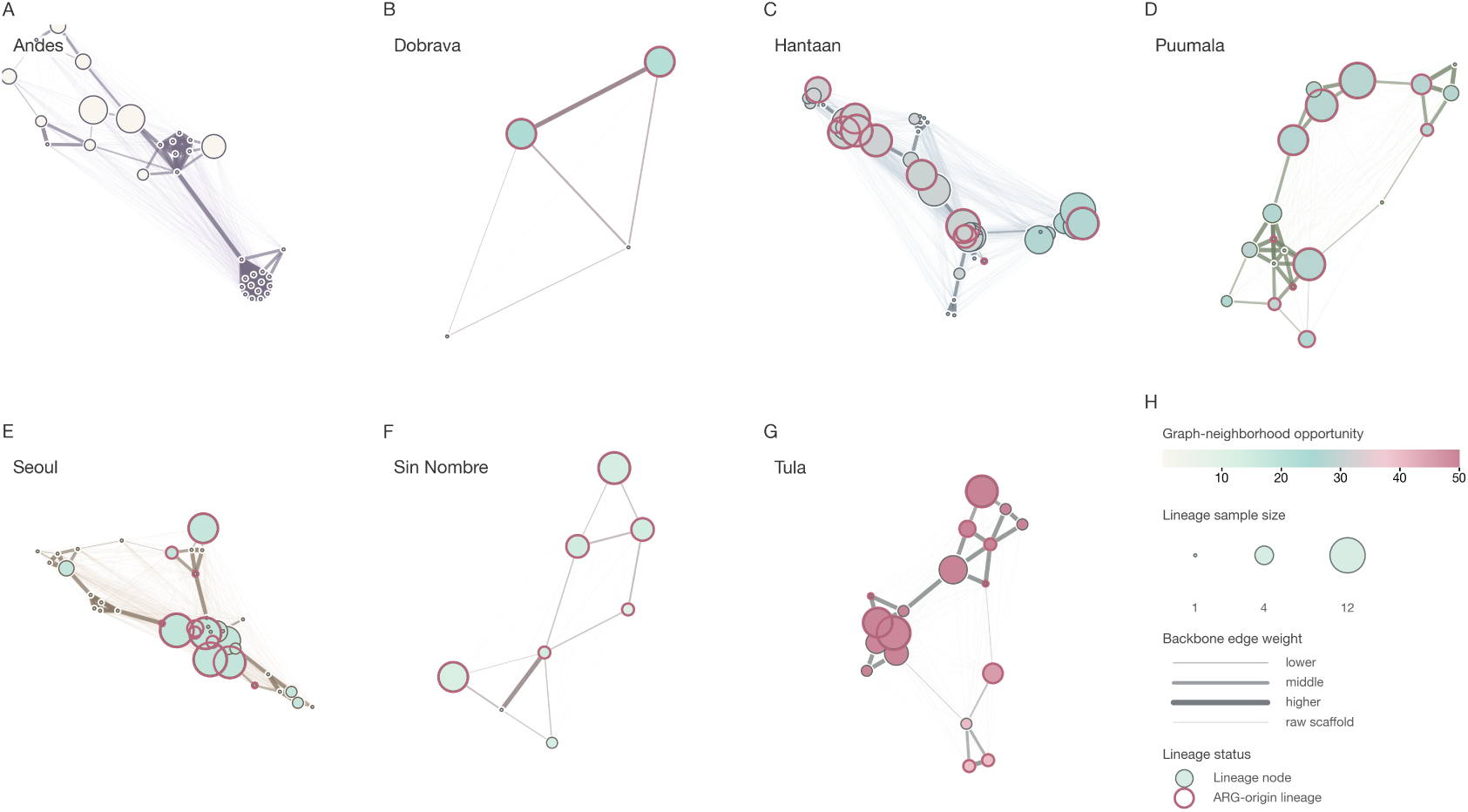
Species-specific lineage graphs used in the establishment analysis. A–G, lineage networks for each virus species, with node size showing lineage sample size, node color showing graph-neighborhood opportunity, and pink outlines marking lineages containing inferred reassortment origins; H, visual encoding key.

**Supplementary Fig. 10:**
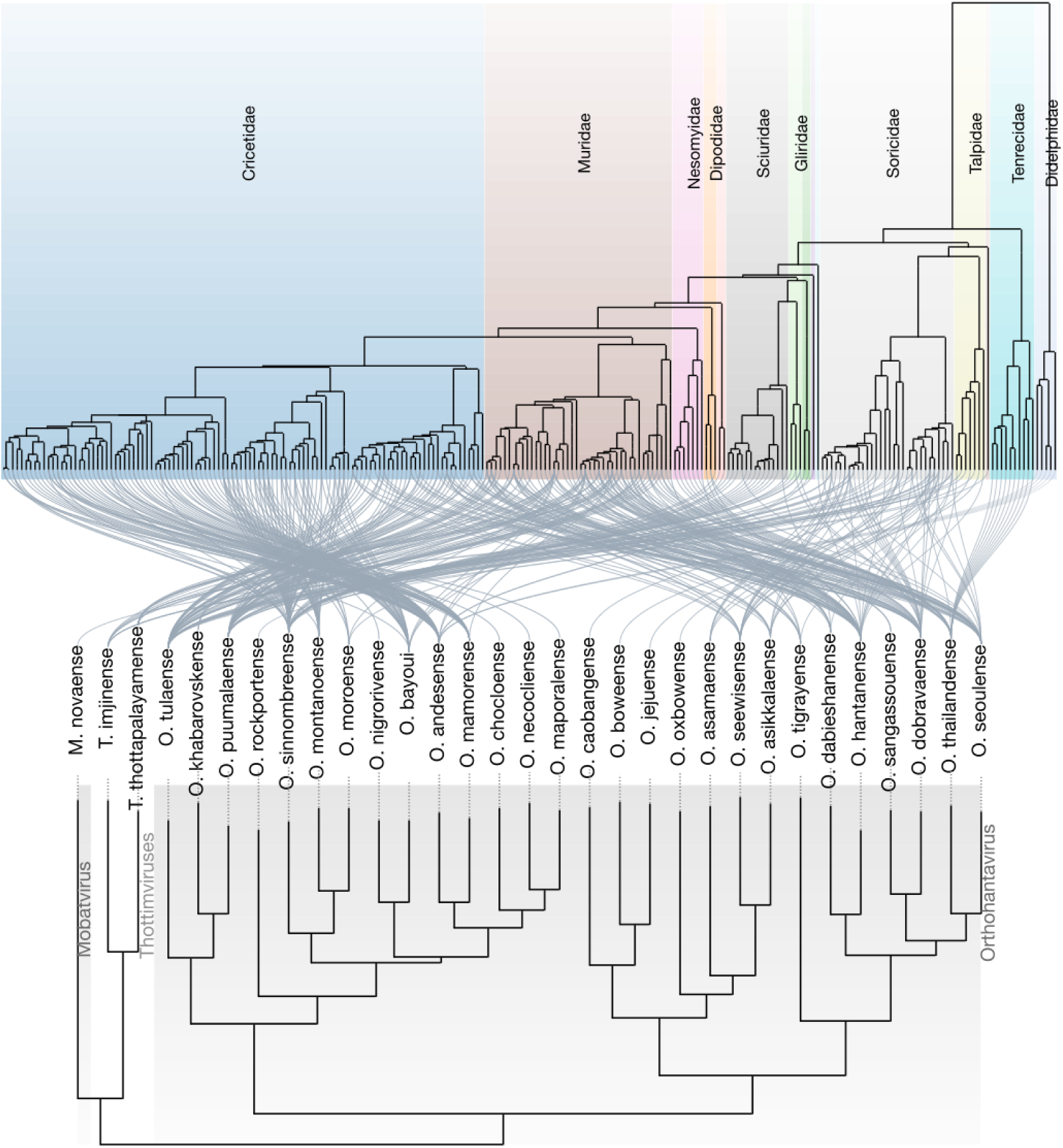
Hantavirus-only host-virus cophylogeny for the evolutionary context used in this study. Links connect orthohantavirus species to mammalian host lineages, with host-family colors showing how reservoir associations are distributed across the host phylogeny.

## Supplementary Tables

External Supplementary Tables 5–7 are provided in supplementary_tables_external.xlsx: Supplementary Table 5 contains host species distribution model hyperparameters and feature-selection counts, Supplementary Table 6 contains the final S/M/L accession triplets and Entrez metadata, and Supplementary Table 7 contains the covariate dictionary with nonmissing fractions. Supplementary Tables 8–10 below provide branch-length-normalized reassortment rates and rootward distance breakdowns, complete posterior parameter estimates for all hierarchical and establishment/expansion models, and sensitivity refits across analytical robustness checks.

**Supplementary Table 1:**
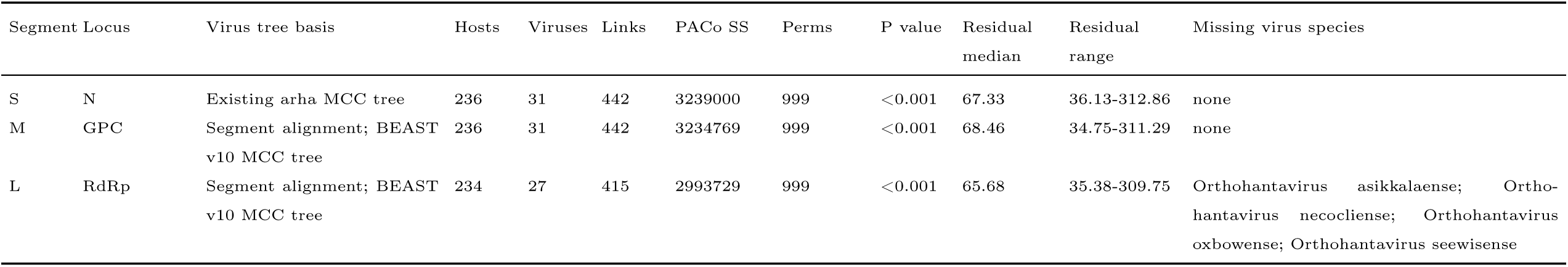
Segment-level PACo host-virus cophylogeny statistics for S, M, and L segment histories. PACo SS is the Procrustes sum of squares from the host-virus fit; smaller residuals indicate closer host-virus phylogenetic association for the included links.

| Segment | Locus | Virus tree basis | Hosts | Viruses | Links | PACo SS | Perms | P value | Residual median | Residual range | Missing virus species |
|---|---|---|---|---|---|---|---|---|---|---|---|
| S | N | Existing arha MCC tree | 236 | 31 | 442 | 3239000 | 999 | <0.001 | 67.33 | 36.13-312.86 | none |
| M | GPC | Segment alignment; BEAST v10 MCC tree | 236 | 31 | 442 | 3234769 | 999 | <0.001 | 68.46 | 34.75-311.29 | none |
| L | RdRp | Segment alignment; BEAST v10 MCC tree | 234 | 27 | 415 | 2993729 | 999 | <0.001 | 65.68 | 35.38-309.75 | Orthohantavirus asikkalaense; Ortho-hantavirus necocliense; Orthohantavirus oxbowense; Orthohantavirus seewisense |

**Supplementary Table 2:**
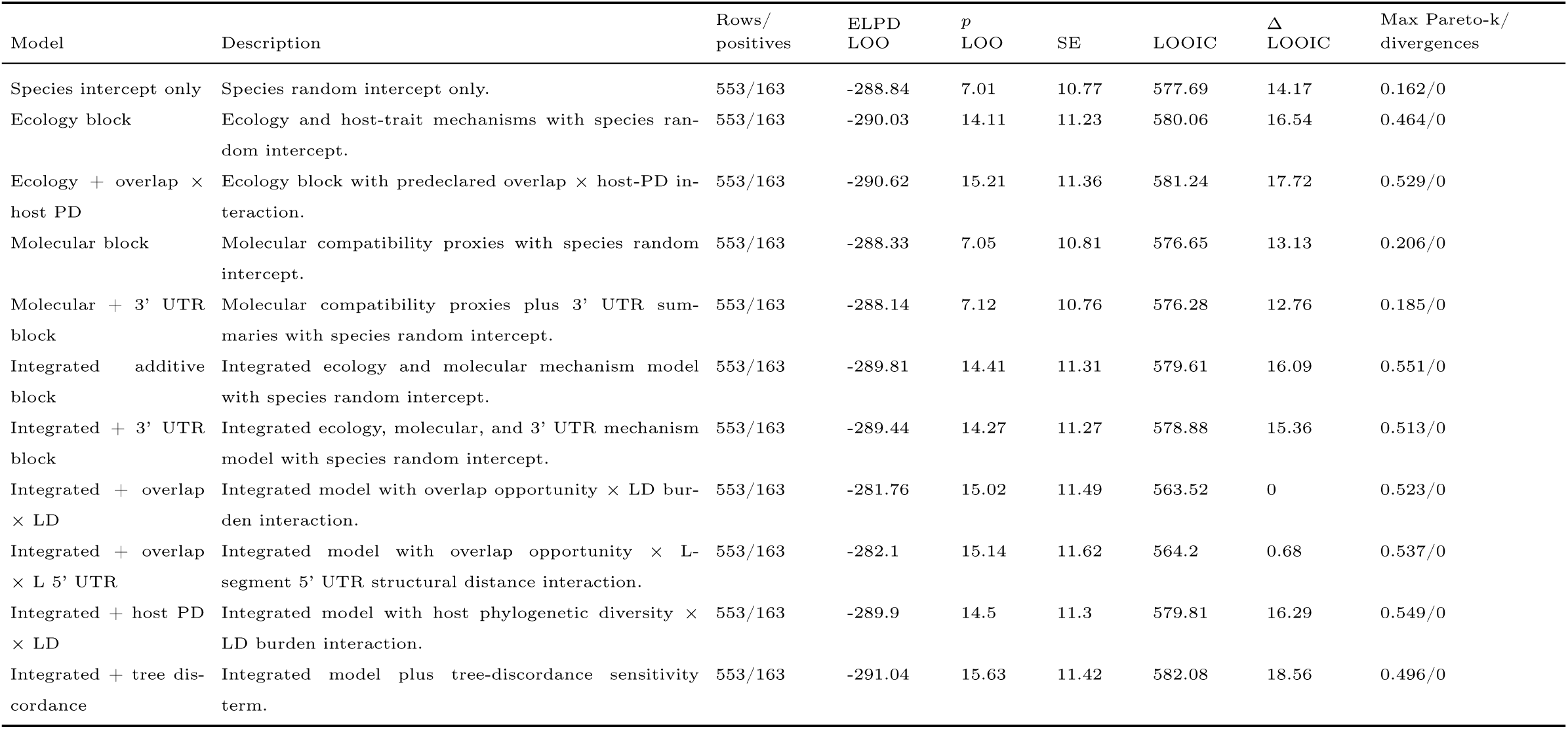
Bayesian hierarchical model-comparison diagnostics for the sequence cohort. LOOIC was calculated as -2 times the leave-one-out expected log predictive density; lower values indicate better expected predictive fit.

| Model | Description | Rows/<br>positives | ELPD<br>LOO | *p*<br>LOO | SE | LOOIC | Δ<br>LOOIC | Max Pareto-k/<br>divergences |
|---|---|---|---|---|---|---|---|---|
| Species intercept only | Species random intercept only. | 553/163 | -288.84 | 7.01 | 10.77 | 577.69 | 14.17 | 0.162/0 |
| Ecology block | Ecology and host-trait mechanisms with species ran-dom intercept. | 553/163 | -290.03 | 14.11 | 11.23 | 580.06 | 16.54 | 0.464/0 |
| Ecology + overlap × host PD | Ecology block with predeclared overlap × host-PD in-teraction. | 553/163 | -290.62 | 15.21 | 11.36 | 581.24 | 17.72 | 0.529/0 |
| Molecular block | Molecular compatibility proxies with species random intercept. | 553/163 | -288.33 | 7.05 | 10.81 | 576.65 | 13.13 | 0.206/0 |
| Molecular + 3' UTR block | Molecular compatibility proxies plus 3' UTR sum-maries with species random intercept. | 553/163 | -288.14 | 7.12 | 10.76 | 576.28 | 12.76 | 0.185/0 |
| Integrated additive block | Integrated ecology and molecular mechanism model with species random intercept. | 553/163 | -289.81 | 14.41 | 11.31 | 579.61 | 16.09 | 0.551/0 |
| Integrated + 3' UTR block | Integrated ecology, molecular, and 3' UTR mechanism model with species random intercept. | 553/163 | -289.44 | 14.27 | 11.27 | 578.88 | 15.36 | 0.513/0 |
| Integrated + overlap × LD | Integrated model with overlap opportunity × LD bur-den interaction. | 553/163 | -281.76 | 15.02 | 11.49 | 563.52 | 0 | 0.523/0 |
| Integrated + overlap × L 5' UTR | Integrated model with overlap opportunity × L-segment 5' UTR structural distance interaction. | 553/163 | -282.1 | 15.14 | 11.62 | 564.2 | 0.68 | 0.537/0 |
| Integrated + host PD × LD | Integrated model with host phylogenetic diversity × LD burden interaction. | 553/163 | -289.9 | 14.5 | 11.3 | 579.81 | 16.29 | 0.549/0 |
| Integrated + tree dis-cordance | Integrated model plus tree-discordance sensitivity term. | 553/163 | -291.04 | 15.63 | 11.42 | 582.08 | 18.56 | 0.496/0 |

**Supplementary Table 3:**
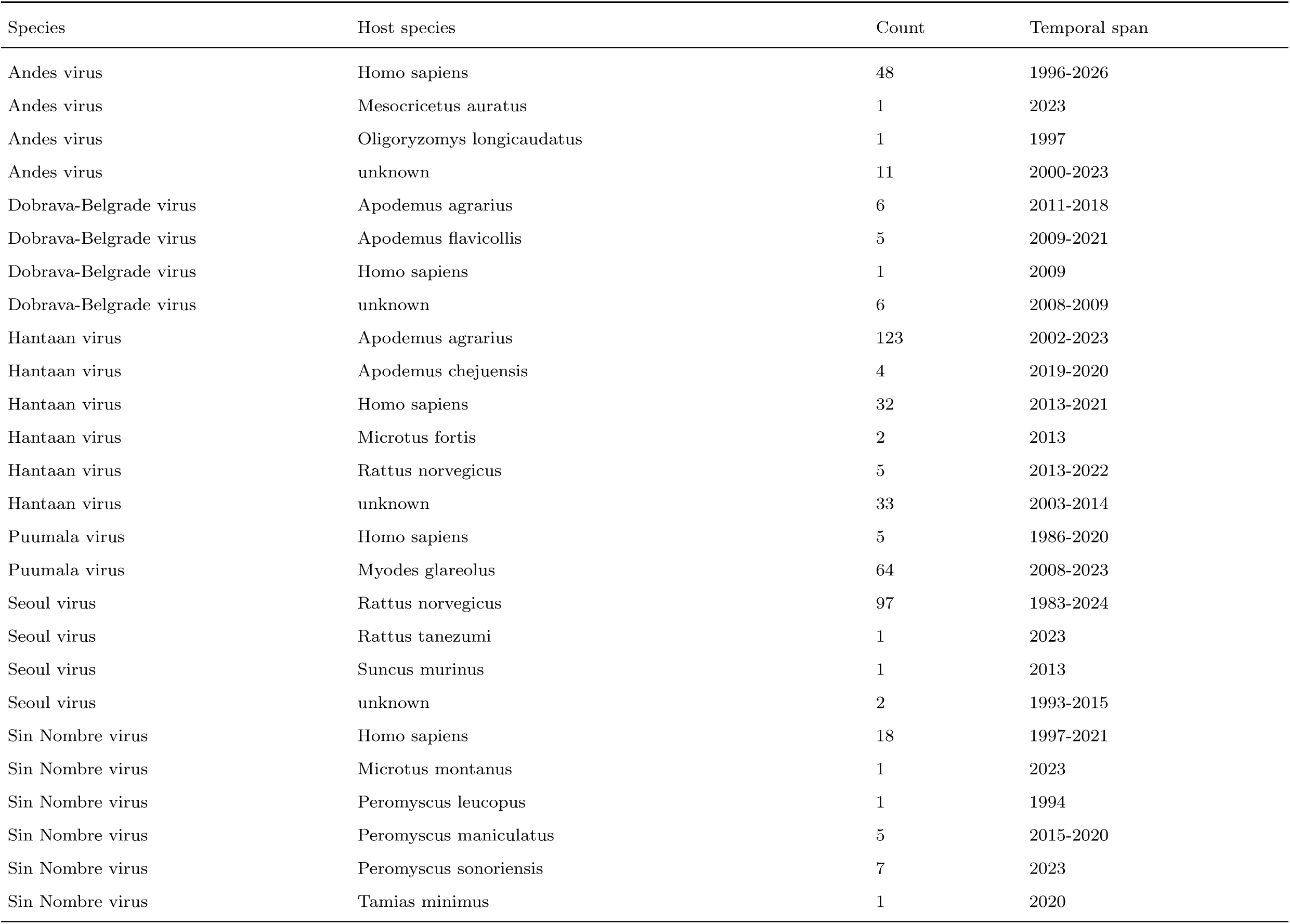
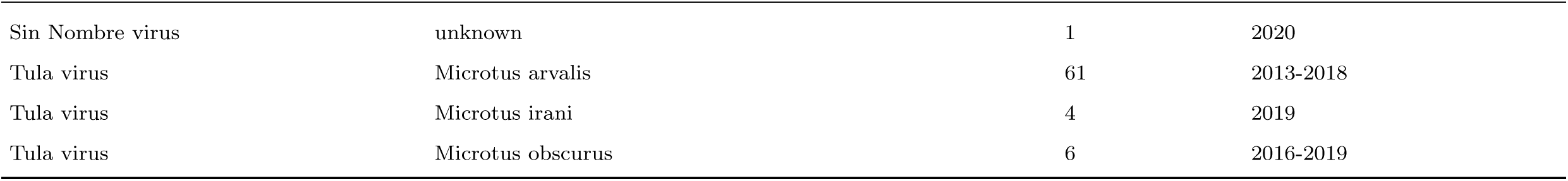
Host species representation and temporal span in the final dataset.

**Supplementary Table 4:**
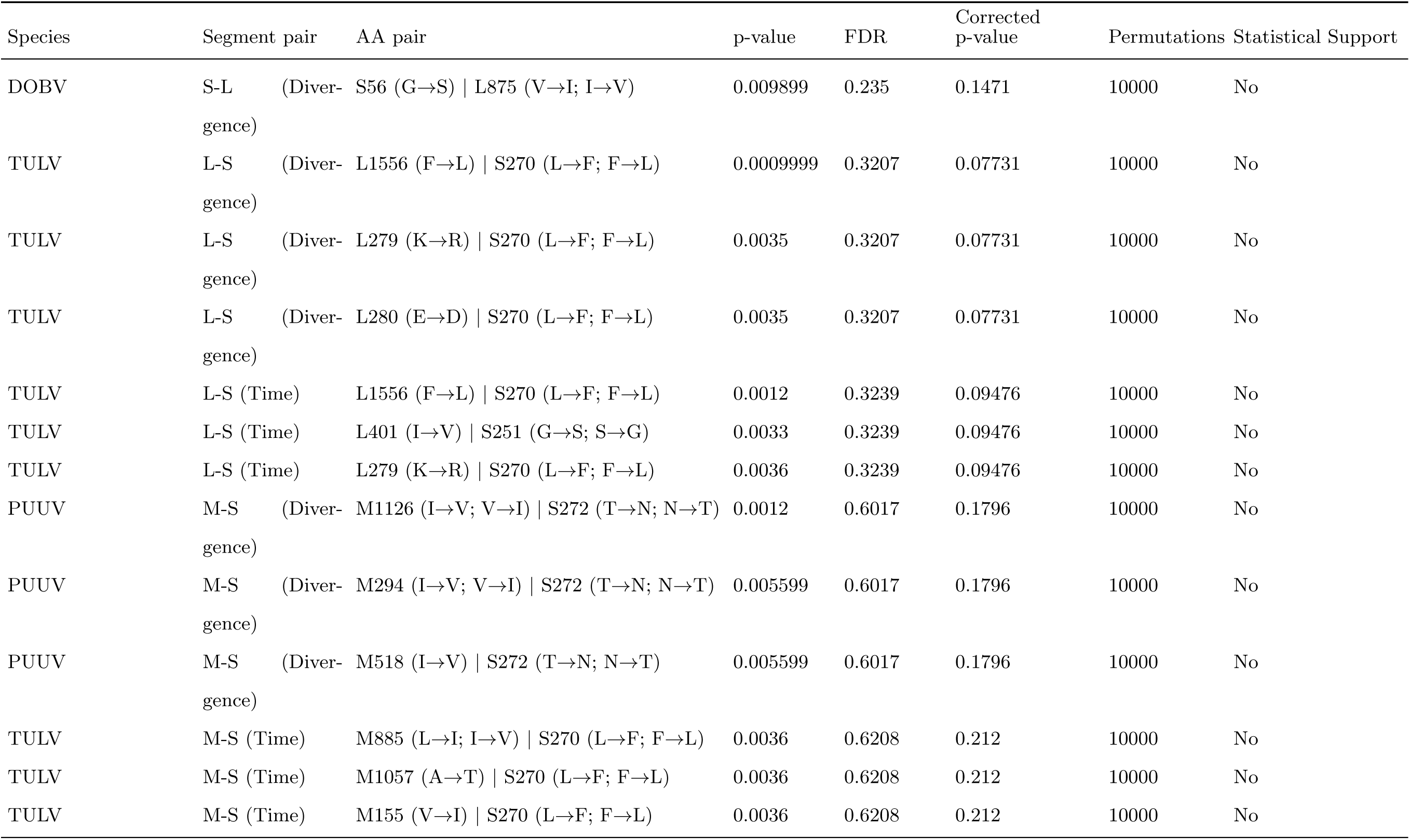
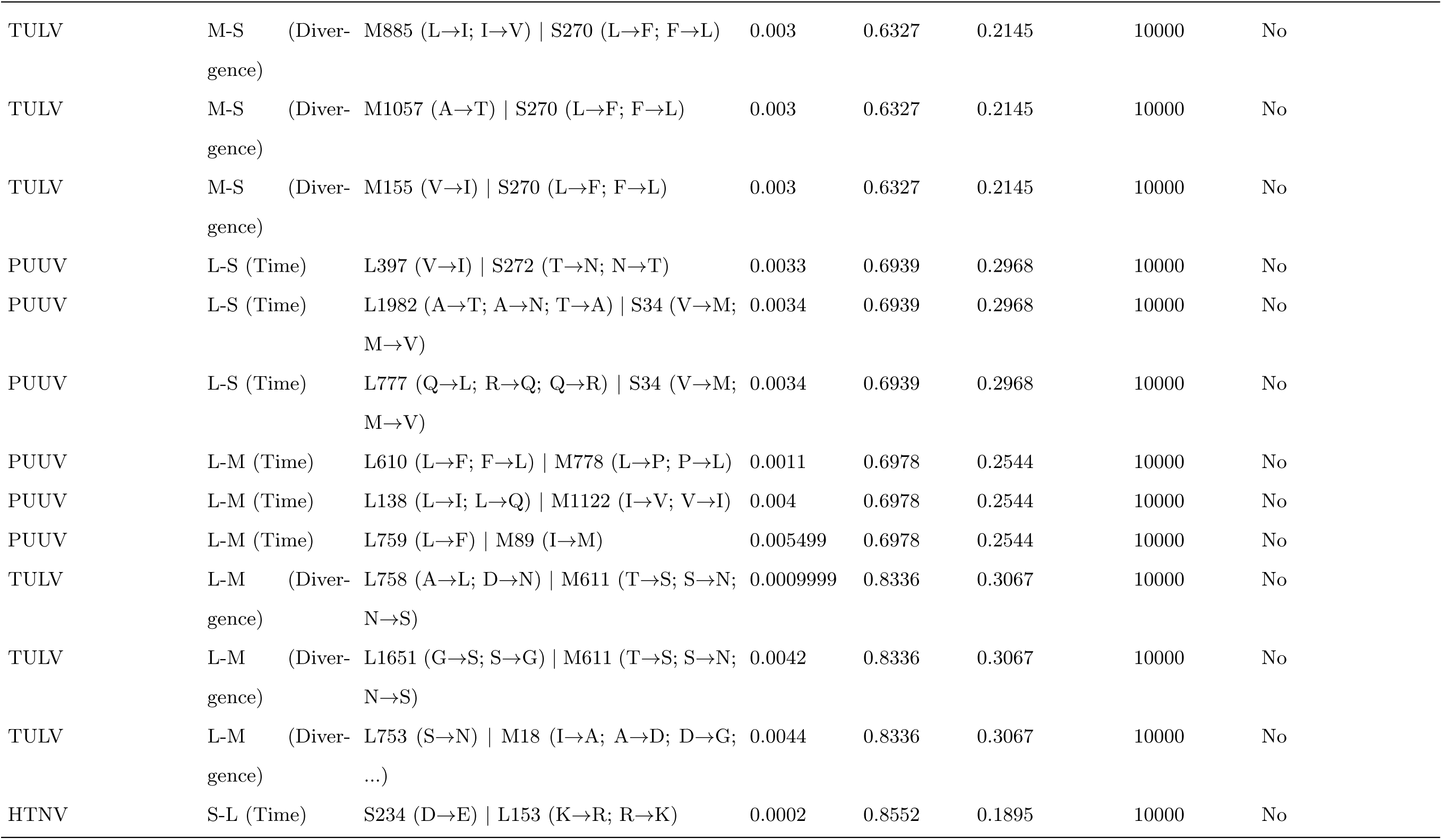
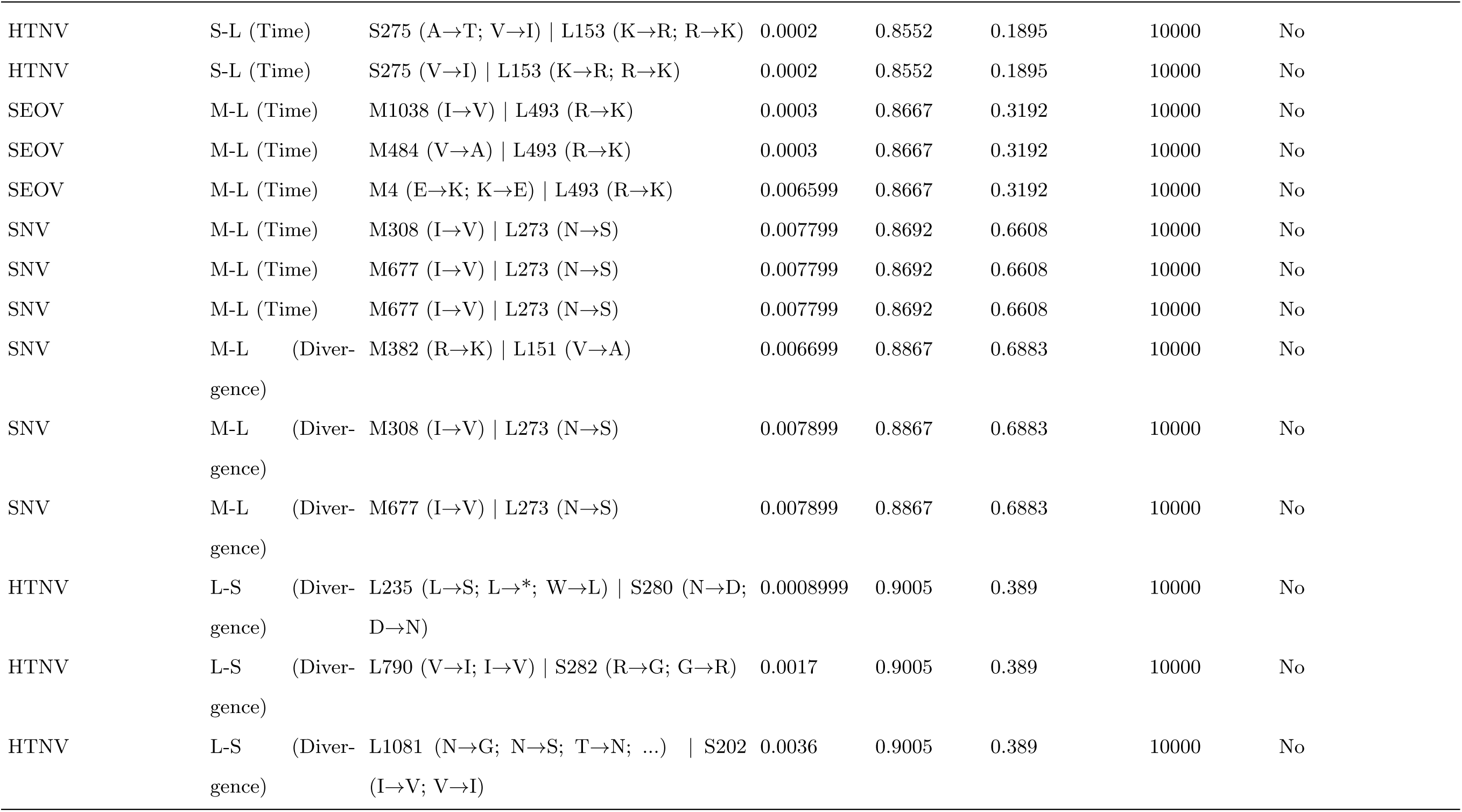
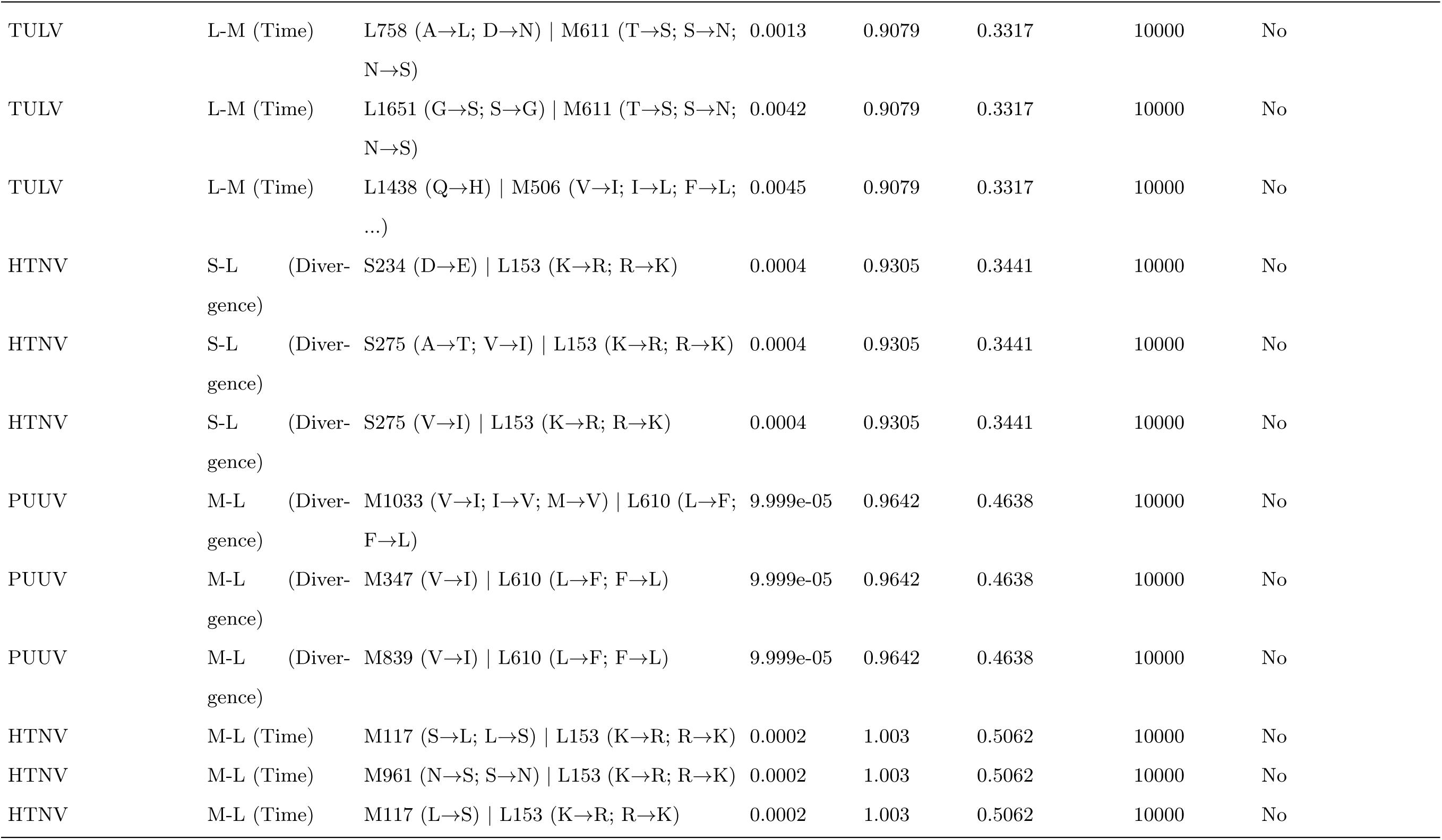
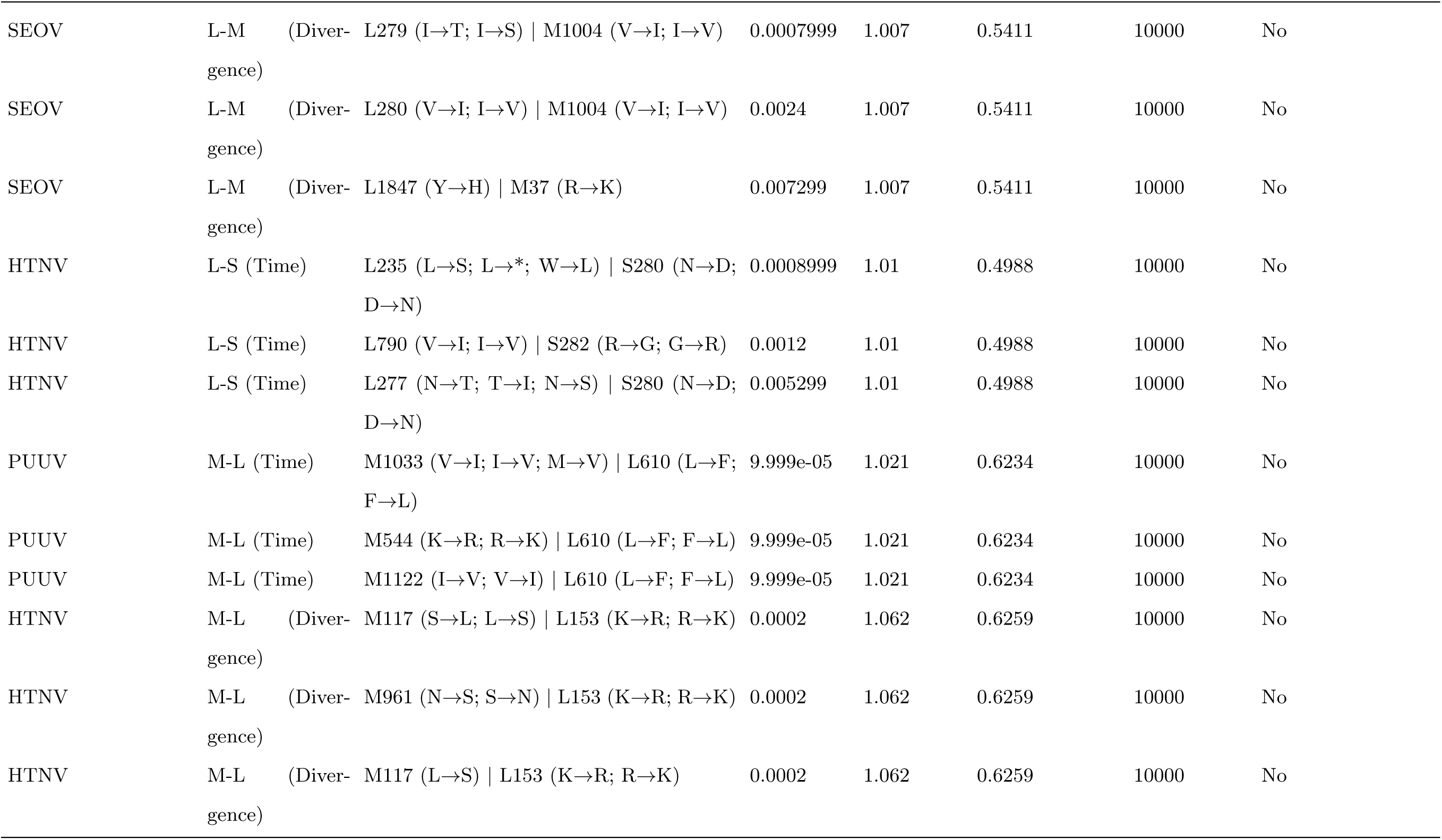
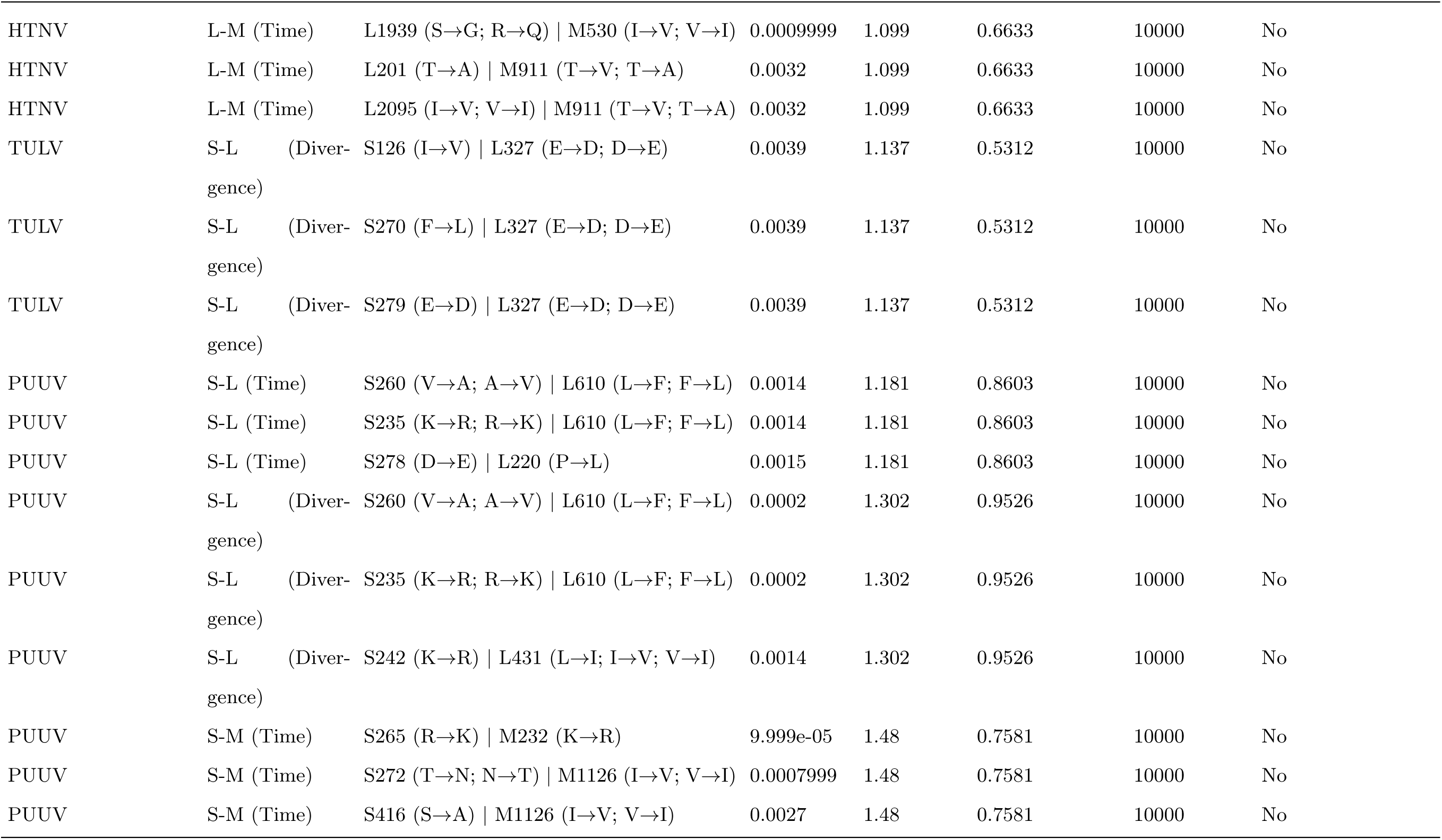
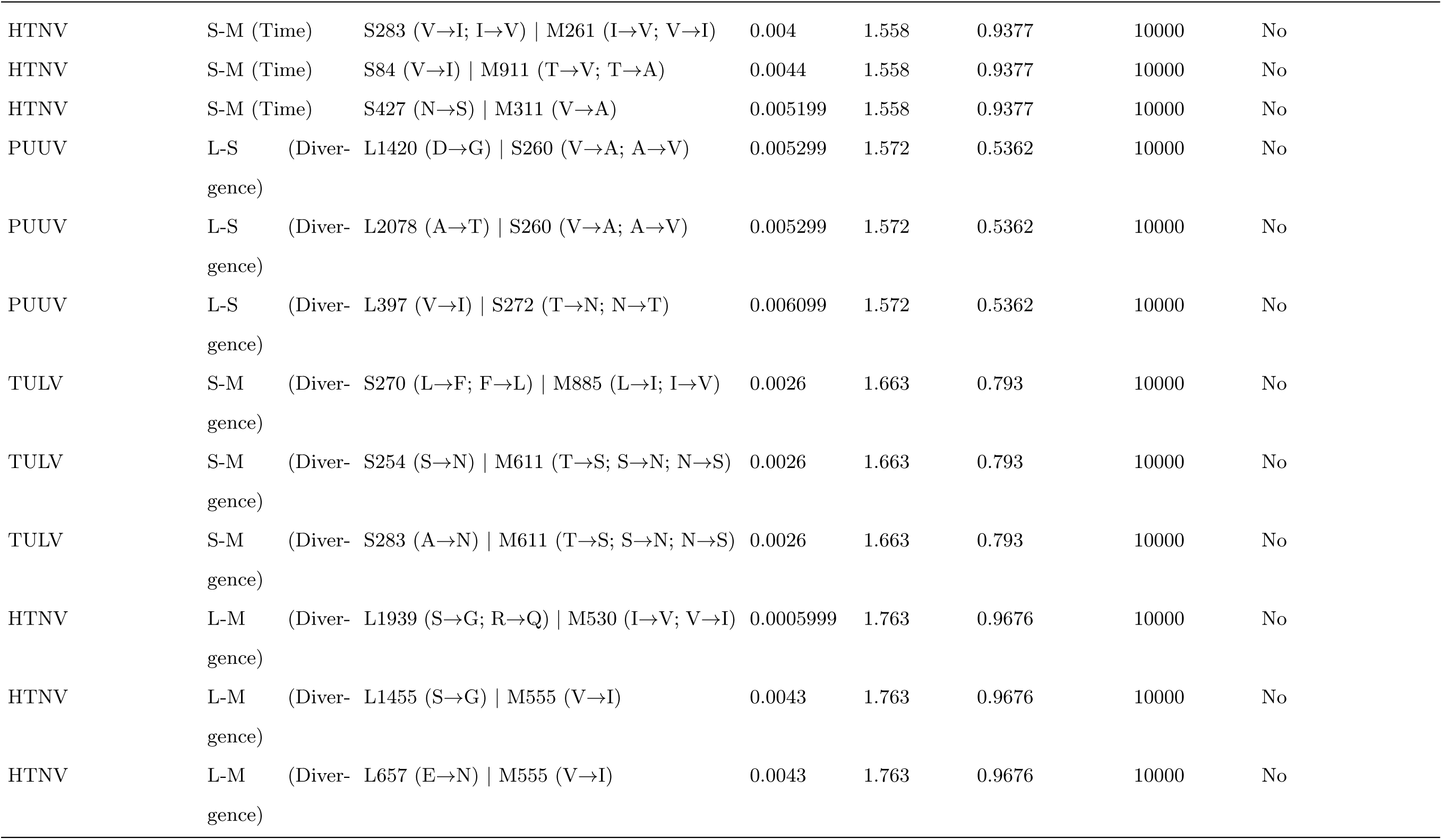
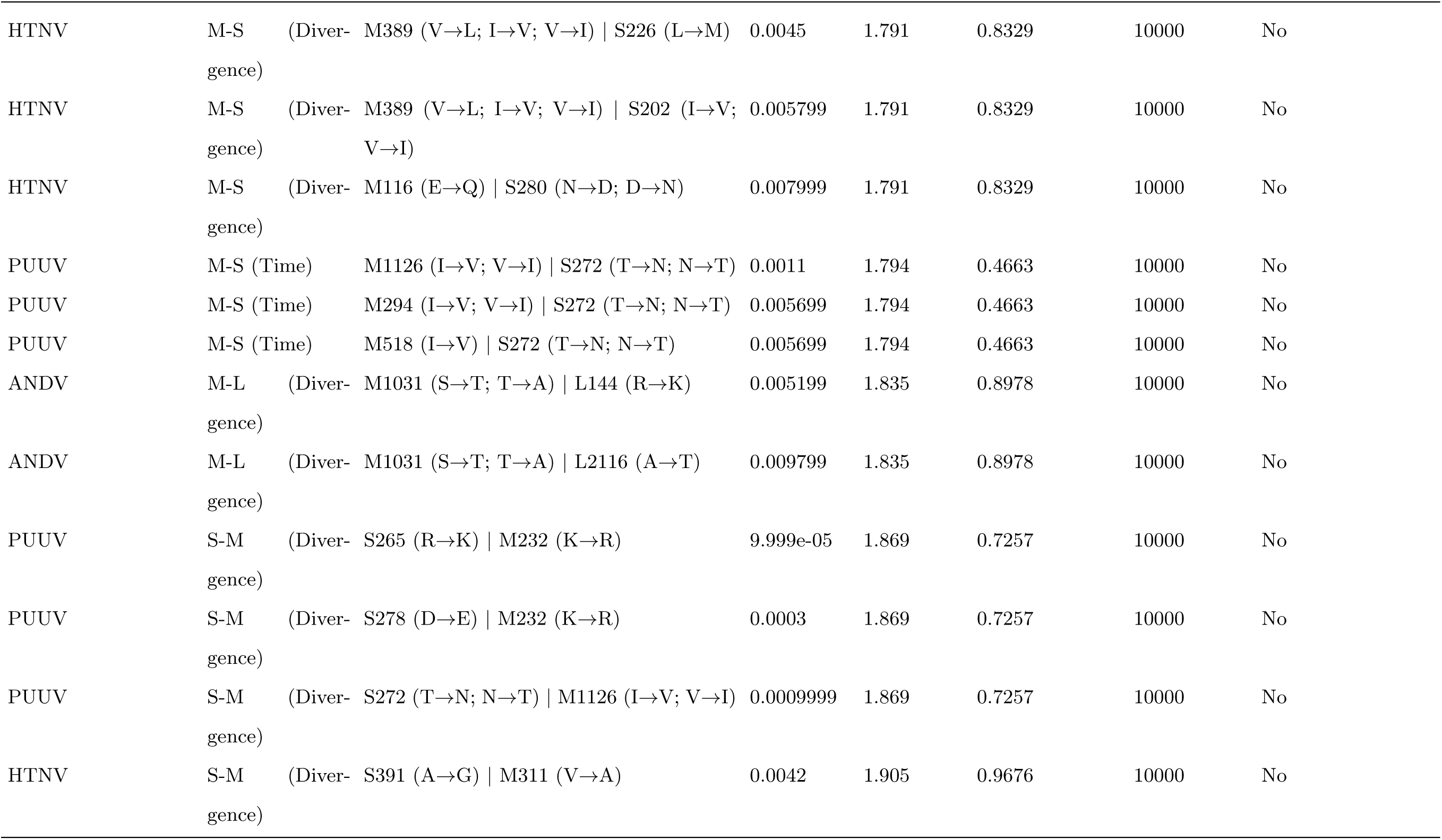
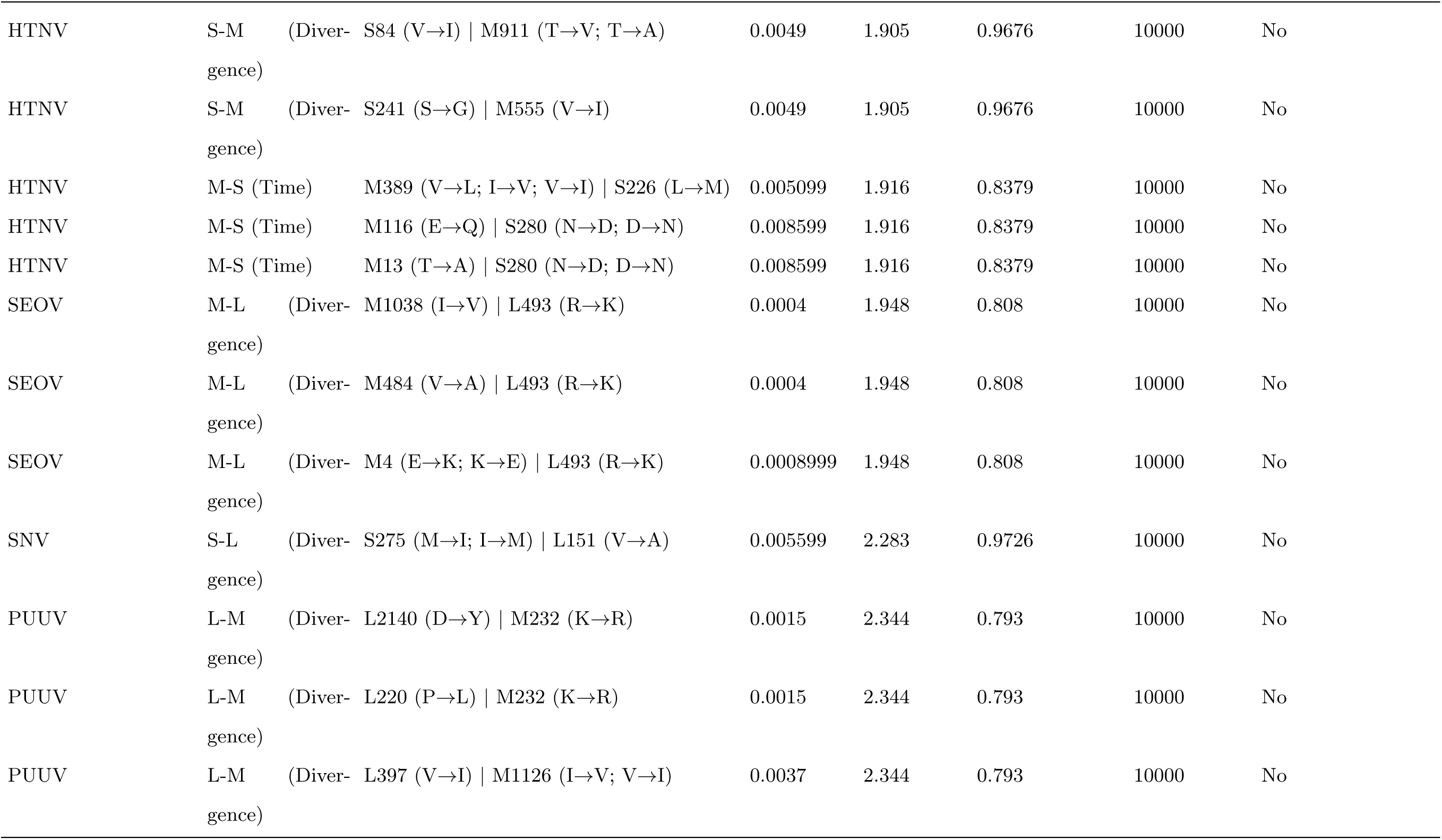
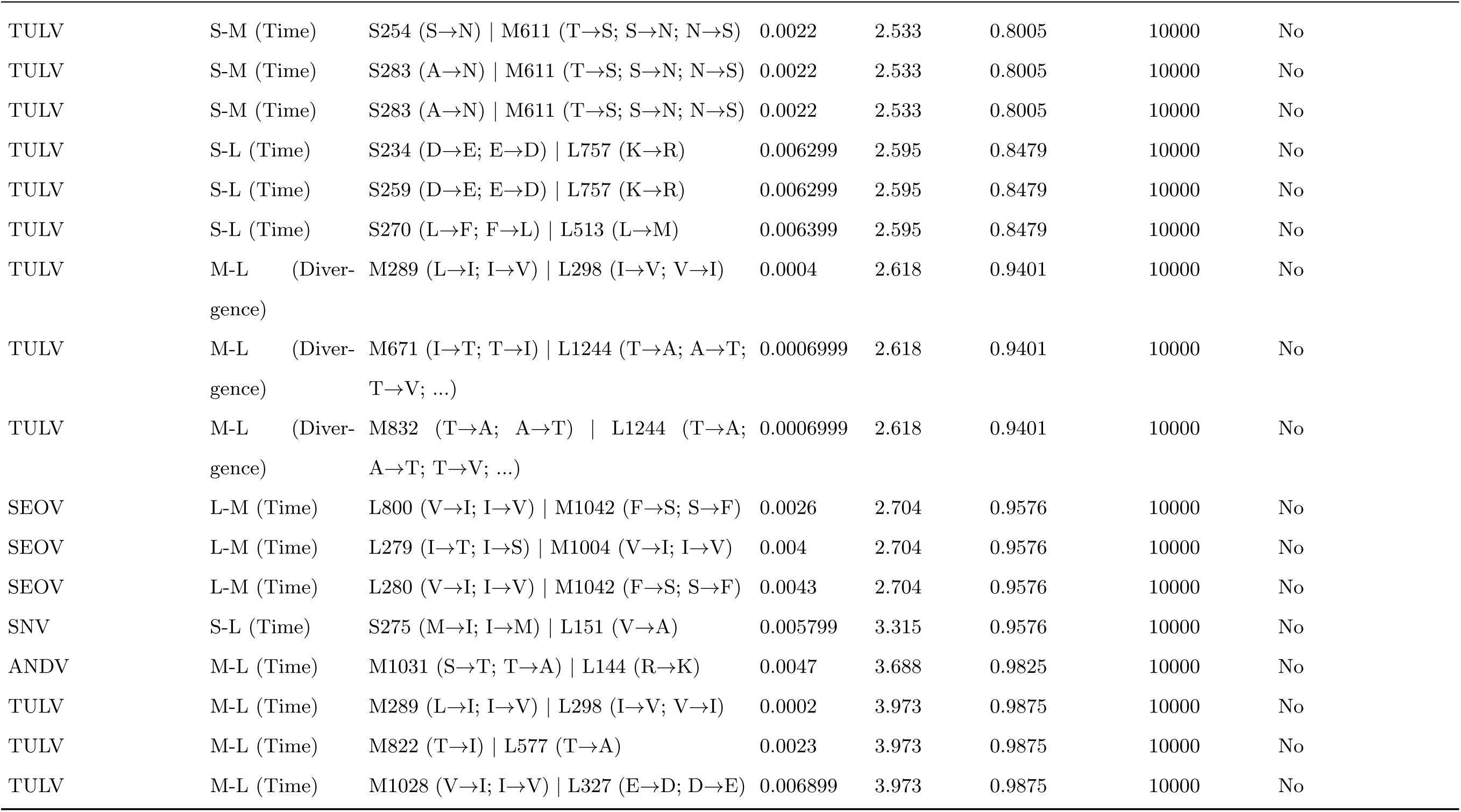
Top nominal epistatic-interaction candidates under the 10,000-permutation selected-interaction screen, with FDR, corrected p-values, and statistical support.

| Species | Segment pair | AA pair | p-value | FDR | Corrected p-value | Permutations | Statistical Support |
|---|---|---|---|---|---|---|---|
| DOBV | S-L (Diver-gence) | S56 (G→S) \| L875 (V→I; I→V) | 0.009899 | 0.235 | 0.1471 | 10000 | No |
| TULV | L-S (Diver-gence) | L1556 (F→L) \| S270 (L→F; F→L) | 0.0009999 | 0.3207 | 0.07731 | 10000 | No |
| TULV | L-S (Diver-gence) | L279 (K→R) \| S270 (L→F; F→L) | 0.0035 | 0.3207 | 0.07731 | 10000 | No |
| TULV | L-S (Diver-gence) | L280 (E→D) \| S270 (L→F; F→L) | 0.0035 | 0.3207 | 0.07731 | 10000 | No |
| TULV | L-S (Time) | L1556 (F→L) \| S270 (L→F; F→L) | 0.0012 | 0.3239 | 0.09476 | 10000 | No |
| TULV | L-S (Time) | L401 (I→V) \| S251 (G→S; S→G) | 0.0033 | 0.3239 | 0.09476 | 10000 | No |
| TULV | L-S (Time) | L279 (K→R) \| S270 (L→F; F→L) | 0.0036 | 0.3239 | 0.09476 | 10000 | No |
| PUUV | M-S (Diver-gence) | M1126 (I→V; V→I) \| S272 (T→N; N→T) | 0.0012 | 0.6017 | 0.1796 | 10000 | No |
| PUUV | M-S (Diver-gence) | M294 (I→V; V→I) \| S272 (T→N; N→T) | 0.005599 | 0.6017 | 0.1796 | 10000 | No |
| PUUV | M-S (Diver-gence) | M518 (I→V) \| S272 (T→N; N→T) | 0.005599 | 0.6017 | 0.1796 | 10000 | No |
| TULV | M-S (Time) | M885 (L→I; I→V) \| S270 (L→F; F→L) | 0.0036 | 0.6208 | 0.212 | 10000 | No |
| TULV | M-S (Time) | M1057 (A→T) \| S270 (L→F; F→L) | 0.0036 | 0.6208 | 0.212 | 10000 | No |
| TULV | M-S (Time) | M155 (V→I) \| S270 (L→F; F→L) | 0.0036 | 0.6208 | 0.212 | 10000 | No |
| TULV | M-S (Diver-gence) | M885 (L→I; I→V) \| S270 (L→F; F→L) | 0.003 | 0.6327 | 0.2145 | 10000 | No |
| TULV | M-S (Diver-gence) | M1057 (A→T) \| S270 (L→F; F→L) | 0.003 | 0.6327 | 0.2145 | 10000 | No |
| TULV | M-S (Diver-gence) | M155 (V→I) \| S270 (L→F; F→L) | 0.003 | 0.6327 | 0.2145 | 10000 | No |
| PUUV | L-S (Time) | L397 (V→I) \| S272 (T→N; N→T) | 0.0033 | 0.6939 | 0.2968 | 10000 | No |
| PUUV | L-S (Time) | L1982 (A→T; A→N; T→A) \| S34 (V→M; M→V) | 0.0034 | 0.6939 | 0.2968 | 10000 | No |
| PUUV | L-S (Time) | L777 (Q→L; R→Q; Q→R) \| S34 (V→M; M→V) | 0.0034 | 0.6939 | 0.2968 | 10000 | No |
| PUUV | L-M (Time) | L610 (L→F; F→L) \| M778 (L→P; P→L) | 0.0011 | 0.6978 | 0.2544 | 10000 | No |
| PUUV | L-M (Time) | L138 (L→I; L→Q) \| M1122 (I→V; V→I) | 0.004 | 0.6978 | 0.2544 | 10000 | No |
| PUUV | L-M (Time) | L759 (L→F) \| M89 (I→M) | 0.005499 | 0.6978 | 0.2544 | 10000 | No |
| TULV | L-M (Diver-gence) | L758 (A→L; D→N) \| M611 (T→S; S→N; N→S) | 0.0009999 | 0.8336 | 0.3067 | 10000 | No |
| TULV | L-M (Diver-gence) | L1651 (G→S; S→G) \| M611 (T→S; S→N; N→S) | 0.0042 | 0.8336 | 0.3067 | 10000 | No |
| TULV | L-M (Diver-gence) | L753 (S→N) \| M18 (I→A; A→D; D→G; ...) | 0.0044 | 0.8336 | 0.3067 | 10000 | No |
| HTNV | S-L (Time) | S234 (D→E) \| L153 (K→R; R→K) | 0.0002 | 0.8552 | 0.1895 | 10000 | No |
| HTNV | S-L (Time) | S275 (A→T; V→I) \| L153 (K→R; R→K) | 0.0002 | 0.8552 | 0.1895 | 10000 | No |
| HTNV | S-L (Time) | S275 (V→I) \| L153 (K→R; R→K) | 0.0002 | 0.8552 | 0.1895 | 10000 | No |
| SEOV | M-L (Time) | M1038 (I→V) \| L493 (R→K) | 0.0003 | 0.8667 | 0.3192 | 10000 | No |
| SEOV | M-L (Time) | M484 (V→A) \| L493 (R→K) | 0.0003 | 0.8667 | 0.3192 | 10000 | No |
| SEOV | M-L (Time) | M4 (E→K; K→E) \| L493 (R→K) | 0.006599 | 0.8667 | 0.3192 | 10000 | No |
| SNV | M-L (Time) | M308 (I→V) \| L273 (N→S) | 0.007799 | 0.8692 | 0.6608 | 10000 | No |
| SNV | M-L (Time) | M677 (I→V) \| L273 (N→S) | 0.007799 | 0.8692 | 0.6608 | 10000 | No |
| SNV | M-L (Time) | M677 (I→V) \| L273 (N→S) | 0.007799 | 0.8692 | 0.6608 | 10000 | No |
| SNV | M-L (Diver-gence) | M382 (R→K) \| L151 (V→A) | 0.006699 | 0.8867 | 0.6883 | 10000 | No |
| SNV | M-L (Diver-gence) | M308 (I→V) \| L273 (N→S) | 0.007899 | 0.8867 | 0.6883 | 10000 | No |
| SNV | M-L (Diver-gence) | M677 (I→V) \| L273 (N→S) | 0.007899 | 0.8867 | 0.6883 | 10000 | No |
| HTNV | L-S (Diver-gence) | L235 (L→S; L→*; W→L) \| S280 (N→D; D→N) | 0.0008999 | 0.9005 | 0.389 | 10000 | No |
| HTNV | L-S (Diver-gence) | L790 (V→I; I→V) \| S282 (R→G; G→R) | 0.0017 | 0.9005 | 0.389 | 10000 | No |
| HTNV | L-S (Diver-gence) | L1081 (N→G; N→S; T→N; ...) \| S202 (I→V; V→I) | 0.0036 | 0.9005 | 0.389 | 10000 | No |
| TULV | L-M (Time) | L758 (A→L; D→N) \| M611 (T→S; S→N; N→S) | 0.0013 | 0.9079 | 0.3317 | 10000 | No |
| TULV | L-M (Time) | L1651 (G→S; S→G) \| M611 (T→S; S→N; N→S) | 0.0042 | 0.9079 | 0.3317 | 10000 | No |
| TULV | L-M (Time) | L1438 (Q→H) \| M506 (V→I; I→L; F→L; ...) | 0.0045 | 0.9079 | 0.3317 | 10000 | No |
| HTNV | S-L (Diver-gence) | S234 (D→E) \| L153 (K→R; R→K) | 0.0004 | 0.9305 | 0.3441 | 10000 | No |
| HTNV | S-L (Diver-gence) | S275 (A→T; V→I) \| L153 (K→R; R→K) | 0.0004 | 0.9305 | 0.3441 | 10000 | No |
| HTNV | S-L (Diver-gence) | S275 (V→I) \| L153 (K→R; R→K) | 0.0004 | 0.9305 | 0.3441 | 10000 | No |
| PUUV | M-L (Diver-gence) | M1033 (V→I; I→V; M→V) \| L610 (L→F; F→L) | 9.999e-05 | 0.9642 | 0.4638 | 10000 | No |
| PUUV | M-L (Diver-gence) | M347 (V→I) \| L610 (L→F; F→L) | 9.999e-05 | 0.9642 | 0.4638 | 10000 | No |
| PUUV | M-L (Diver-gence) | M839 (V→I) \| L610 (L→F; F→L) | 9.999e-05 | 0.9642 | 0.4638 | 10000 | No |
| HTNV | M-L (Time) | M117 (S→L; L→S) \| L153 (K→R; R→K) | 0.0002 | 1.003 | 0.5062 | 10000 | No |
| HTNV | M-L (Time) | M961 (N→S; S→N) \| L153 (K→R; R→K) | 0.0002 | 1.003 | 0.5062 | 10000 | No |
| HTNV | M-L (Time) | M117 (L→S) \| L153 (K→R; R→K) | 0.0002 | 1.003 | 0.5062 | 10000 | No |
| SEOV | L-M (Diver-gence) | L279 (I→T; I→S) \| M1004 (V→I; I→V) | 0.0007999 | 1.007 | 0.5411 | 10000 | No |
| SEOV | L-M (Diver-gence) | L280 (V→I; I→V) \| M1004 (V→I; I→V) | 0.0024 | 1.007 | 0.5411 | 10000 | No |
| SEOV | L-M (Diver-gence) | L1847 (Y→H) \| M37 (R→K) | 0.007299 | 1.007 | 0.5411 | 10000 | No |
| HTNV | L-S (Time) | L235 (L→S; L→*; W→L) \| S280 (N→D; D→N) | 0.0008999 | 1.01 | 0.4988 | 10000 | No |
| HTNV | L-S (Time) | L790 (V→I; I→V) \| S282 (R→G; G→R) | 0.0012 | 1.01 | 0.4988 | 10000 | No |
| HTNV | L-S (Time) | L277 (N→T; T→I; N→S) \| S280 (N→D; D→N) | 0.005299 | 1.01 | 0.4988 | 10000 | No |
| PUUV | M-L (Time) | M1033 (V→I; I→V; M→V) \| L610 (L→F; F→L) | 9.999e-05 | 1.021 | 0.6234 | 10000 | No |
| PUUV | M-L (Time) | M544 (K→R; R→K) \| L610 (L→F; F→L) | 9.999e-05 | 1.021 | 0.6234 | 10000 | No |
| PUUV | M-L (Time) | M1122 (I→V; V→I) \| L610 (L→F; F→L) | 9.999e-05 | 1.021 | 0.6234 | 10000 | No |
| HTNV | M-L (Diver-gence) | M117 (S→L; L→S) \| L153 (K→R; R→K) | 0.0002 | 1.062 | 0.6259 | 10000 | No |
| HTNV | M-L (Diver-gence) | M961 (N→S; S→N) \| L153 (K→R; R→K) | 0.0002 | 1.062 | 0.6259 | 10000 | No |
| HTNV | M-L (Diver-gence) | M117 (L→S) \| L153 (K→R; R→K) | 0.0002 | 1.062 | 0.6259 | 10000 | No |
| HTNV | L-M (Time) | L1939 (S→G; R→Q) \| M530 (I→V; V→I) | 0.0009999 | 1.099 | 0.6633 | 10000 | No |
| HTNV | L-M (Time) | L201 (T→A) \| M911 (T→V; T→A) | 0.0032 | 1.099 | 0.6633 | 10000 | No |
| HTNV | L-M (Time) | L2095 (I→V; V→I) \| M911 (T→V; T→A) | 0.0032 | 1.099 | 0.6633 | 10000 | No |
| TULV | S-L (Diver-gence) | S126 (I→V) \| L327 (E→D; D→E) | 0.0039 | 1.137 | 0.5312 | 10000 | No |
| TULV | S-L (Diver-gence) | S270 (F→L) \| L327 (E→D; D→E) | 0.0039 | 1.137 | 0.5312 | 10000 | No |
| TULV | S-L (Diver-gence) | S279 (E→D) \| L327 (E→D; D→E) | 0.0039 | 1.137 | 0.5312 | 10000 | No |
| PUUV | S-L (Time) | S260 (V→A; A→V) \| L610 (L→F; F→L) | 0.0014 | 1.181 | 0.8603 | 10000 | No |
| PUUV | S-L (Time) | S235 (K→R; R→K) \| L610 (L→F; F→L) | 0.0014 | 1.181 | 0.8603 | 10000 | No |
| PUUV | S-L (Time) | S278 (D→E) \| L220 (P→L) | 0.0015 | 1.181 | 0.8603 | 10000 | No |
| PUUV | S-L (Diver-gence) | S260 (V→A; A→V) \| L610 (L→F; F→L) | 0.0002 | 1.302 | 0.9526 | 10000 | No |
| PUUV | S-L (Diver-gence) | S235 (K→R; R→K) \| L610 (L→F; F→L) | 0.0002 | 1.302 | 0.9526 | 10000 | No |
| PUUV | S-L (Diver-gence) | S242 (K→R) \| L431 (L→I; I→V; V→I) | 0.0014 | 1.302 | 0.9526 | 10000 | No |
| PUUV | S-M (Time) | S265 (R→K) \| M232 (K→R) | 9.999e-05 | 1.48 | 0.7581 | 10000 | No |
| PUUV | S-M (Time) | S272 (T→N; N→T) \| M1126 (I→V; V→I) | 0.0007999 | 1.48 | 0.7581 | 10000 | No |
| PUUV | S-M (Time) | S416 (S→A) \| M1126 (I→V; V→I) | 0.0027 | 1.48 | 0.7581 | 10000 | No |
| HTNV | S-M (Time) | S283 (V→I; I→V) \| M261 (I→V; V→I) | 0.004 | 1.558 | 0.9377 | 10000 | No |
| HTNV | S-M (Time) | S84 (V→I) \| M911 (T→V; T→A) | 0.0044 | 1.558 | 0.9377 | 10000 | No |
| HTNV | S-M (Time) | S427 (N→S) \| M311 (V→A) | 0.005199 | 1.558 | 0.9377 | 10000 | No |
| PUUV | L-S (Diver-gence) | L1420 (D→G) \| S260 (V→A; A→V) | 0.005299 | 1.572 | 0.5362 | 10000 | No |
| PUUV | L-S (Diver-gence) | L2078 (A→T) \| S260 (V→A; A→V) | 0.005299 | 1.572 | 0.5362 | 10000 | No |
| PUUV | L-S (Diver-gence) | L397 (V→I) \| S272 (T→N; N→T) | 0.006099 | 1.572 | 0.5362 | 10000 | No |
| TULV | S-M (Diver-gence) | S270 (L→F; F→L) \| M885 (L→I; I→V) | 0.0026 | 1.663 | 0.793 | 10000 | No |
| TULV | S-M (Diver-gence) | S254 (S→N) \| M611 (T→S; S→N; N→S) | 0.0026 | 1.663 | 0.793 | 10000 | No |
| TULV | S-M (Diver-gence) | S283 (A→N) \| M611 (T→S; S→N; N→S) | 0.0026 | 1.663 | 0.793 | 10000 | No |
| HTNV | L-M (Diver-gence) | L1939 (S→G; R→Q) \| M530 (I→V; V→I) | 0.0005999 | 1.763 | 0.9676 | 10000 | No |
| HTNV | L-M (Diver-gence) | L1455 (S→G) \| M555 (V→I) | 0.0043 | 1.763 | 0.9676 | 10000 | No |
| HTNV | L-M (Diver-gence) | L657 (E→N) \| M555 (V→I) | 0.0043 | 1.763 | 0.9676 | 10000 | No |
| HTNV | M-S (Diver-gence) | M389 (V→L; I→V; V→I) \| S226 (L→M) | 0.0045 | 1.791 | 0.8329 | 10000 | No |
| HTNV | M-S (Diver-gence) | M389 (V→L; I→V; V→I) \| S202 (I→V; V→I) | 0.005799 | 1.791 | 0.8329 | 10000 | No |
| HTNV | M-S (Diver-gence) | M116 (E→Q) \| S280 (N→D; D→N) | 0.007999 | 1.791 | 0.8329 | 10000 | No |
| PUUV | M-S (Time) | M1126 (I→V; V→I) \| S272 (T→N; N→T) | 0.0011 | 1.794 | 0.4663 | 10000 | No |
| PUUV | M-S (Time) | M294 (I→V; V→I) \| S272 (T→N; N→T) | 0.005699 | 1.794 | 0.4663 | 10000 | No |
| PUUV | M-S (Time) | M518 (I→V) \| S272 (T→N; N→T) | 0.005699 | 1.794 | 0.4663 | 10000 | No |
| ANDV | M-L (Diver-gence) | M1031 (S→T; T→A) \| L144 (R→K) | 0.005199 | 1.835 | 0.8978 | 10000 | No |
| ANDV | M-L (Diver-gence) | M1031 (S→T; T→A) \| L2116 (A→T) | 0.009799 | 1.835 | 0.8978 | 10000 | No |
| PUUV | S-M (Diver-gence) | S265 (R→K) \| M232 (K→R) | 9.999e-05 | 1.869 | 0.7257 | 10000 | No |
| PUUV | S-M (Diver-gence) | S278 (D→E) \| M232 (K→R) | 0.0003 | 1.869 | 0.7257 | 10000 | No |
| PUUV | S-M (Diver-gence) | S272 (T→N; N→T) \| M1126 (I→V; V→I) | 0.0009999 | 1.869 | 0.7257 | 10000 | No |
| HTNV | S-M (Diver-gence) | S391 (A→G) \| M311 (V→A) | 0.0042 | 1.905 | 0.9676 | 10000 | No |
| HTNV | S-M (Diver-gence) | S84 (V→I) \| M911 (T→V; T→A) | 0.0049 | 1.905 | 0.9676 | 10000 | No |
| HTNV | S-M (Diver-gence) | S241 (S→G) \| M555 (V→I) | 0.0049 | 1.905 | 0.9676 | 10000 | No |
| HTNV | M-S (Time) | M389 (V→L; I→V; V→I) \| S226 (L→M) | 0.005099 | 1.916 | 0.8379 | 10000 | No |
| HTNV | M-S (Time) | M116 (E→Q) \| S280 (N→D; D→N) | 0.008599 | 1.916 | 0.8379 | 10000 | No |
| HTNV | M-S (Time) | M13 (T→A) \| S280 (N→D; D→N) | 0.008599 | 1.916 | 0.8379 | 10000 | No |
| SEOV | M-L (Diver-gence) | M1038 (I→V) \| L493 (R→K) | 0.0004 | 1.948 | 0.808 | 10000 | No |
| SEOV | M-L (Diver-gence) | M484 (V→A) \| L493 (R→K) | 0.0004 | 1.948 | 0.808 | 10000 | No |
| SEOV | M-L (Diver-gence) | M4 (E→K; K→E) \| L493 (R→K) | 0.0008999 | 1.948 | 0.808 | 10000 | No |
| SNV | S-L (Diver-gence) | S275 (M→I; I→M) \| L151 (V→A) | 0.005599 | 2.283 | 0.9726 | 10000 | No |
| PUUV | L-M (Diver-gence) | L2140 (D→Y) \| M232 (K→R) | 0.0015 | 2.344 | 0.793 | 10000 | No |
| PUUV | L-M (Diver-gence) | L220 (P→L) \| M232 (K→R) | 0.0015 | 2.344 | 0.793 | 10000 | No |
| PUUV | L-M (Diver-gence) | L397 (V→I) \| M1126 (I→V; V→I) | 0.0037 | 2.344 | 0.793 | 10000 | No |
| TULV | S-M (Time) | S254 (S→N) \| M611 (T→S; S→N; N→S) | 0.0022 | 2.533 | 0.8005 | 10000 | No |
| TULV | S-M (Time) | S283 (A→N) \| M611 (T→S; S→N; N→S) | 0.0022 | 2.533 | 0.8005 | 10000 | No |
| TULV | S-M (Time) | S283 (A→N) \| M611 (T→S; S→N; N→S) | 0.0022 | 2.533 | 0.8005 | 10000 | No |
| TULV | S-L (Time) | S234 (D→E; E→D) \| L757 (K→R) | 0.006299 | 2.595 | 0.8479 | 10000 | No |
| TULV | S-L (Time) | S259 (D→E; E→D) \| L757 (K→R) | 0.006299 | 2.595 | 0.8479 | 10000 | No |
| TULV | S-L (Time) | S270 (L→F; F→L) \| L513 (L→M) | 0.006399 | 2.595 | 0.8479 | 10000 | No |
| TULV | M-L (Diver-gence) | M289 (L→I; I→V) \| L298 (I→V; V→I) | 0.0004 | 2.618 | 0.9401 | 10000 | No |
| TULV | M-L (Diver-gence) | M671 (I→T; T→I) \| L1244 (T→A; A→T; T→V; ...) | 0.0006999 | 2.618 | 0.9401 | 10000 | No |
| TULV | M-L (Diver-gence) | M832 (T→A; A→T) \| L1244 (T→A; A→T; T→V; ...) | 0.0006999 | 2.618 | 0.9401 | 10000 | No |
| SEOV | L-M (Time) | L800 (V→I; I→V) \| M1042 (F→S; S→F) | 0.0026 | 2.704 | 0.9576 | 10000 | No |
| SEOV | L-M (Time) | L279 (I→T; I→S) \| M1004 (V→I; I→V) | 0.004 | 2.704 | 0.9576 | 10000 | No |
| SEOV | L-M (Time) | L280 (V→I; I→V) \| M1042 (F→S; S→F) | 0.0043 | 2.704 | 0.9576 | 10000 | No |
| SNV | S-L (Time) | S275 (M→I; I→M) \| L151 (V→A) | 0.005799 | 3.315 | 0.9576 | 10000 | No |
| ANDV | M-L (Time) | M1031 (S→T; T→A) \| L144 (R→K) | 0.0047 | 3.688 | 0.9825 | 10000 | No |
| TULV | M-L (Time) | M289 (L→I; I→V) \| L298 (I→V; V→I) | 0.0002 | 3.973 | 0.9875 | 10000 | No |
| TULV | M-L (Time) | M822 (T→I) \| L577 (T→A) | 0.0023 | 3.973 | 0.9875 | 10000 | No |
| TULV | M-L (Time) | M1028 (V→I; I→V) \| L327 (E→D; D→E) | 0.006899 | 3.973 | 0.9875 | 10000 | No |

**Supplementary Table 8:** Branch-length-normalized and tip-normalized reassortment origin rates and rootward path distance threshold sensitivity across the seven focal orthohantavirus species (*n* = 553 complete genomes). Candidate and minimal (de-nested) origin clusters were extracted from reconciled Espalier ancestral recombination graphs (ARGs). Segment tree heights (*H*) and total branch lengths (*L*_tot_) are reported in substitutions per site for common-tip time-calibrated phylogenies (with ANDV S-segment metrics evaluated on the 59-tip common tree). Branch-length-normalized origin rates equal minimal origin clusters divided by mean segment total branch length (*L̄*_tot_), with exact 95% Poisson confidence intervals (one-sided 95% upper bound for ANDV). Columns *d* = 1, *d ≤* 2 (primary), *d ≤* 3, and *d >* 3*/d* = *∞* report positive tip counts under alternative rootward internal-node distance cutoffs to the nearest ancestral reassortment node.

| Species | Sampled genomes (*n*) | Primary *d* ≤ 2 tips (*k*, % [95% CI]) | Origins (cand./ minimal) | Tips per minimal origin | Tree height S / M / L (mean *H̄*) | Total branch length S / M / L (mean *L̄*<sub>tot</sub>) | Origins per subs./site [95% CI] | Origins per tip (orig./*n*) | Strict *d* = 1 (*k*, %) | Relaxed *d* ≤ 3 (*k*, %) | Deep *d* > 3 / no anc. (*d* = ∞) |
|---|---|---|---|---|---|---|---|---|---|---|---|
| Andes | 61 | 0 (0.0% [0.0–5.9]) | 0 / 0 | — | 0.081/0.047/0.052 (0.060) | 0.22/0.32/0.34 (0.30) | 0.00 [0.00–10.15] | 0.000 | 0 (0.0%) | 0 (0.0%) | 0 / 61 |
| Dobrava | 18 | 12 (66.7% [43.7–83.7]) | 7 / 4 | 3.00 | 0.085/0.306/0.220 (0.204) | 0.46/1.19/1.01 (0.88) | 4.52 [1.23–11.59] | 0.222 | 9 (50.0%) | 15 (83.3%) | 0 / 3 |
| Hantaan | 199 | 29 (14.6% [10.3–20.1]) | 20 / 14 | 2.07 | 0.161/0.193/0.147 (0.167) | 1.59/1.89/1.58 (1.69) | 8.31 [4.54–13.94] | 0.070 | 23 (11.6%) | 36 (18.1%) | 156 / 7 |
| Puumala | 69 | 28 (40.6% [29.8–52.4]) | 28 / 14 | 2.00 | 0.118/0.141/0.370 (0.210) | 0.89/0.83/2.91 (1.54) | 9.08 [4.96–15.23] | 0.203 | 17 (24.6%) | 32 (46.4%) | 26 / 11 |
| Seoul | 101 | 47 (46.5% [37.1–56.2]) | 38 / 18 | 2.61 | 0.087/0.212/0.144 (0.148) | 0.93/1.62/0.96 (1.17) | 15.37 [9.11–24.28] | 0.178 | 30 (29.7%) | 60 (59.4%) | 41 / 0 |
| Sin Nombre | 34 | 20 (58.8% [42.2–73.6]) | 13 / 9 | 2.22 | 0.371/0.339/0.353 (0.354) | 1.56/1.56/1.75 (1.62) | 5.55 [2.54–10.53] | 0.265 | 14 (41.2%) | 20 (58.8%) | 0 / 14 |
| Tula | 71 | 27 (38.0% [27.6–49.7]) | 25 / 14 | 1.93 | 0.216/0.248/0.330 (0.265) | 1.41/2.10/2.27 (1.93) | 7.26 [3.97–12.19] | 0.197 | 20 (28.2%) | 30 (42.3%) | 18 / 23 |
| Total / Pooled | 553 | 163 (29.5% [25.8–33.4]) | 131 / 73 | 2.23 | mean 0.201 | sum 9.13 | 8.00 [5.85–9.38] | 0.132 | 113 (20.4%) | 193 (34.9%) | 241 / 119 |

**Supplementary Table 9:**
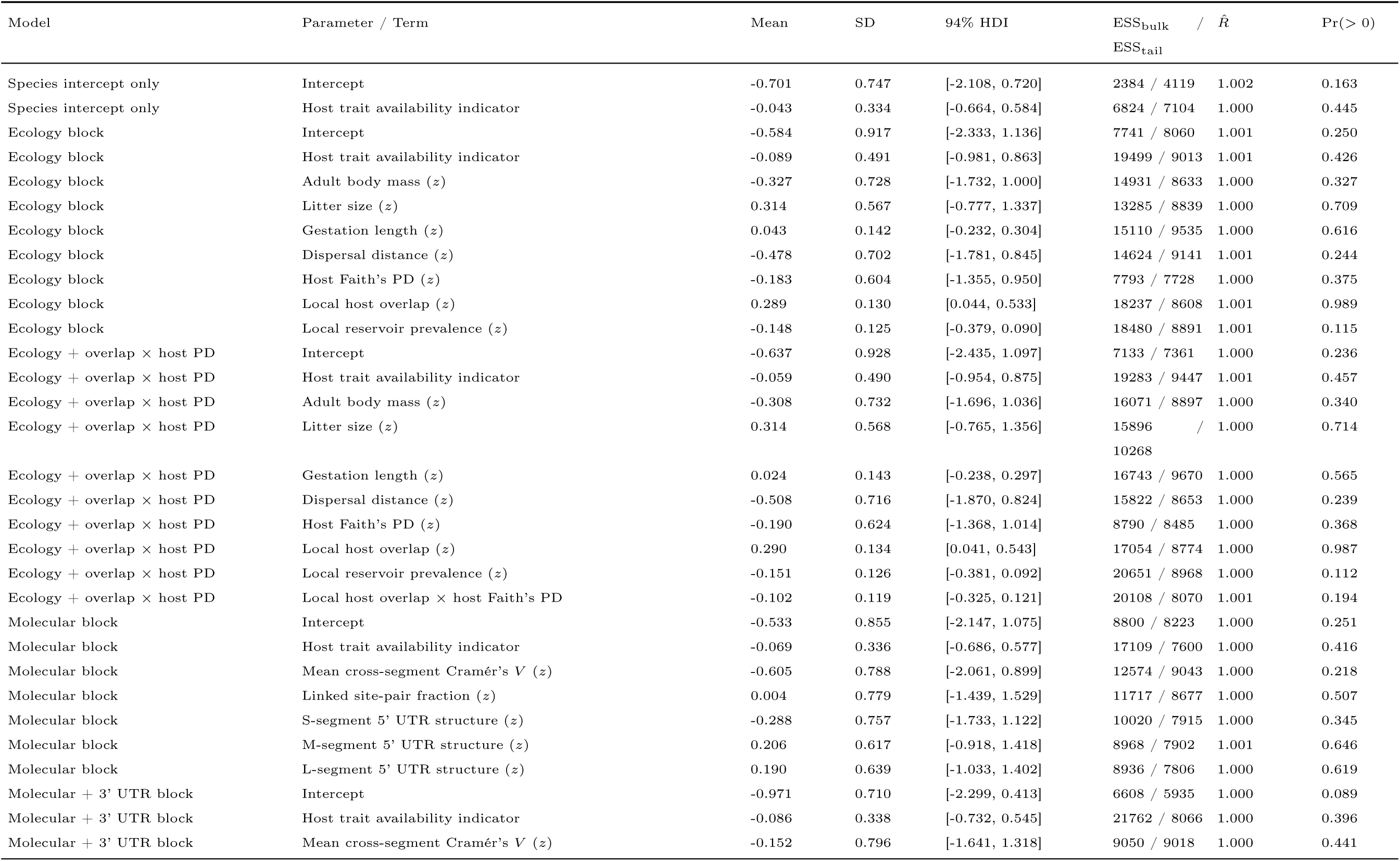
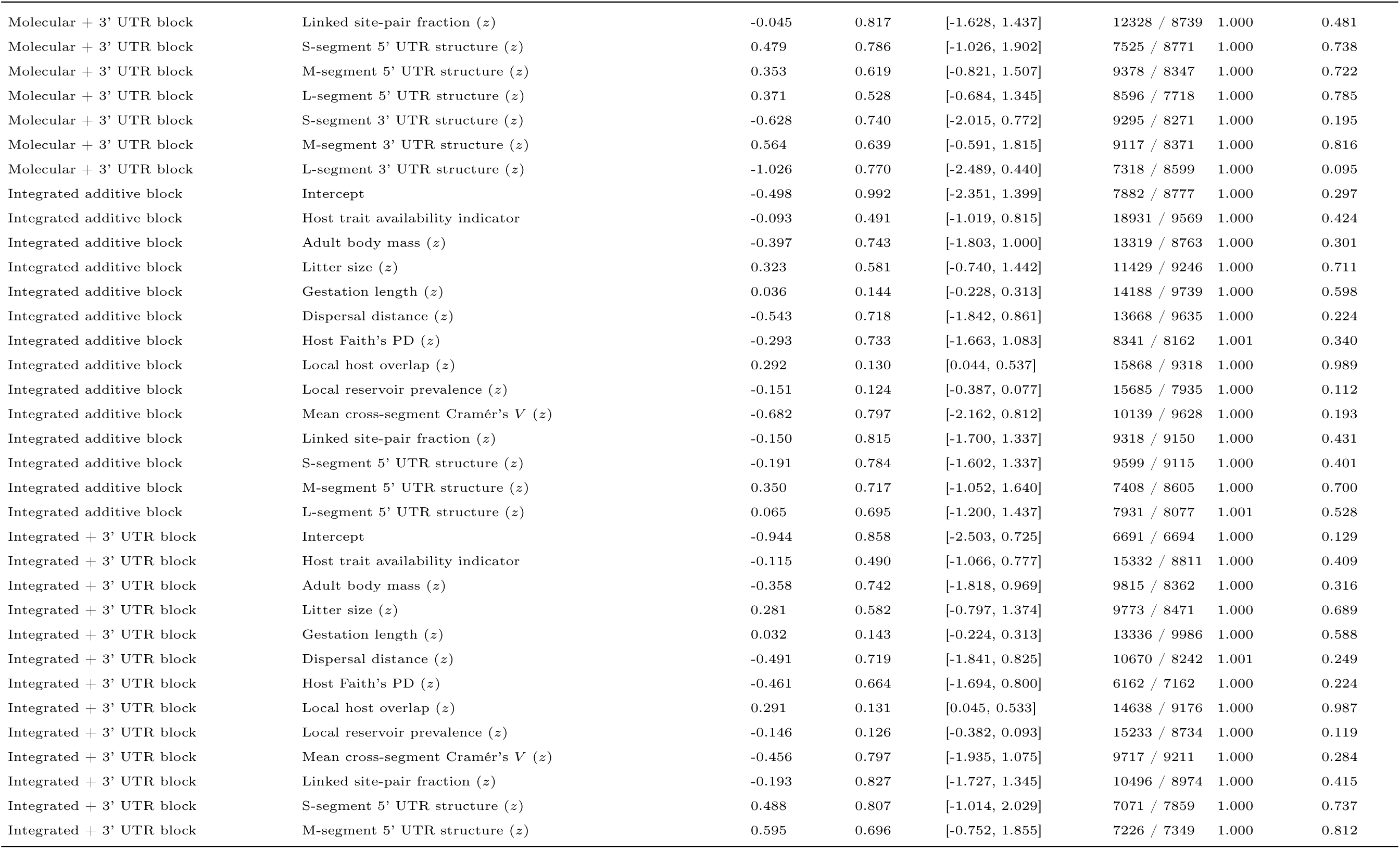
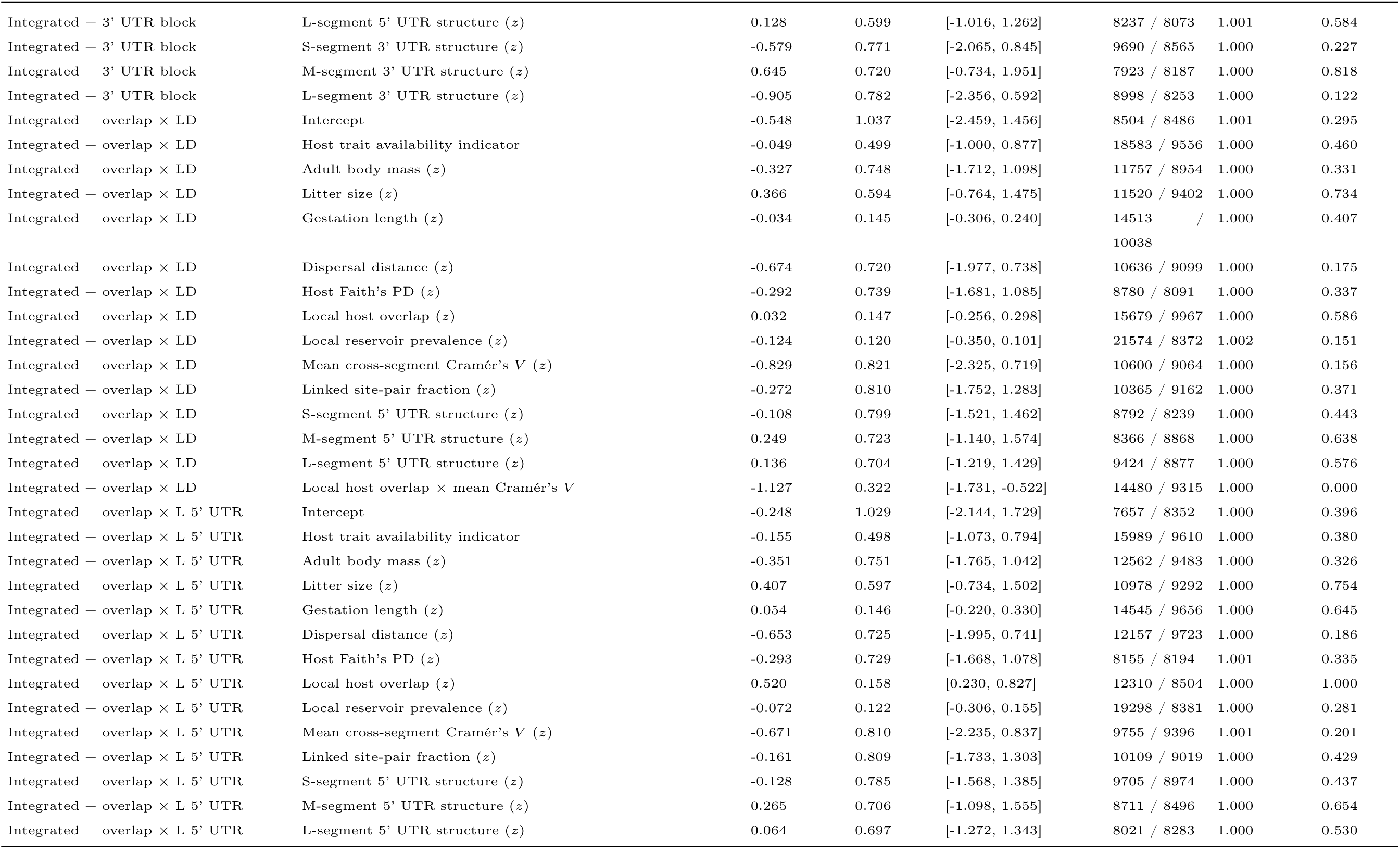
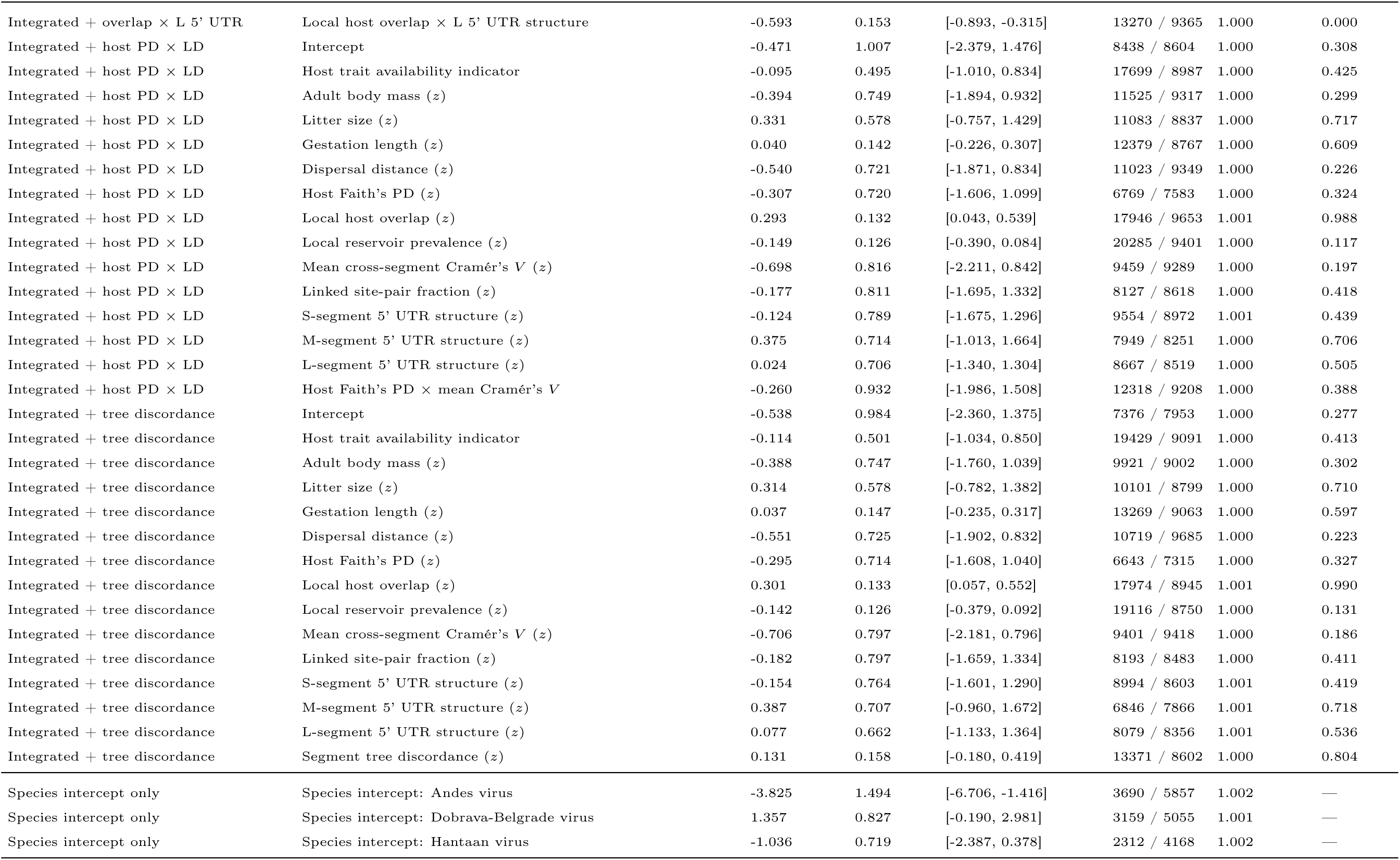
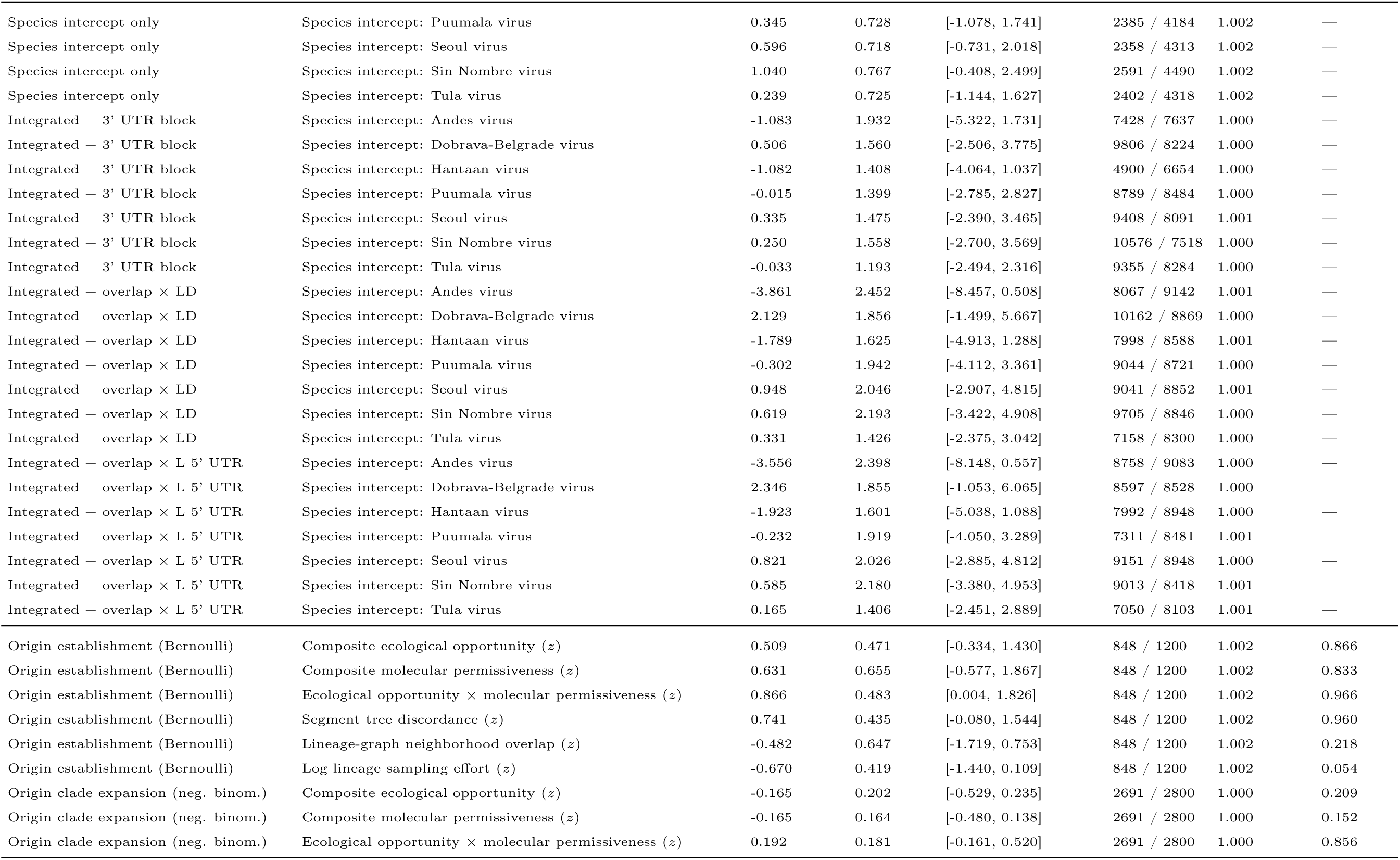
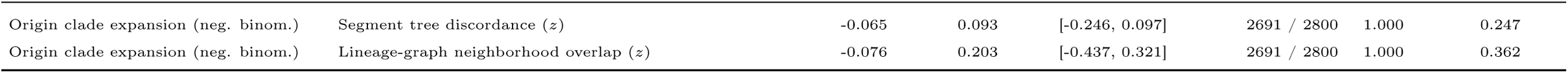
Complete posterior parameter estimates (posterior mean, standard deviation [SD], 94% highest-density interval [HDI], bulk and tail effective sample sizes [ESS], split-*R*^^^, and directional posterior probability Pr(*β >* 0)) for all 11 sequence-level Bayesian hierarchical logistic regression models (*n* = 553, 163 positives), key species random intercepts (*u_s_*), the origin-level Bayesian Bernoulli establishment model (*n* = 275: 73 minimal origins vs. 202 matched controls), and the origin-level negative-binomial clade-expansion count model (*n* = 73 minimal origins).

**Supplementary Table 10:**
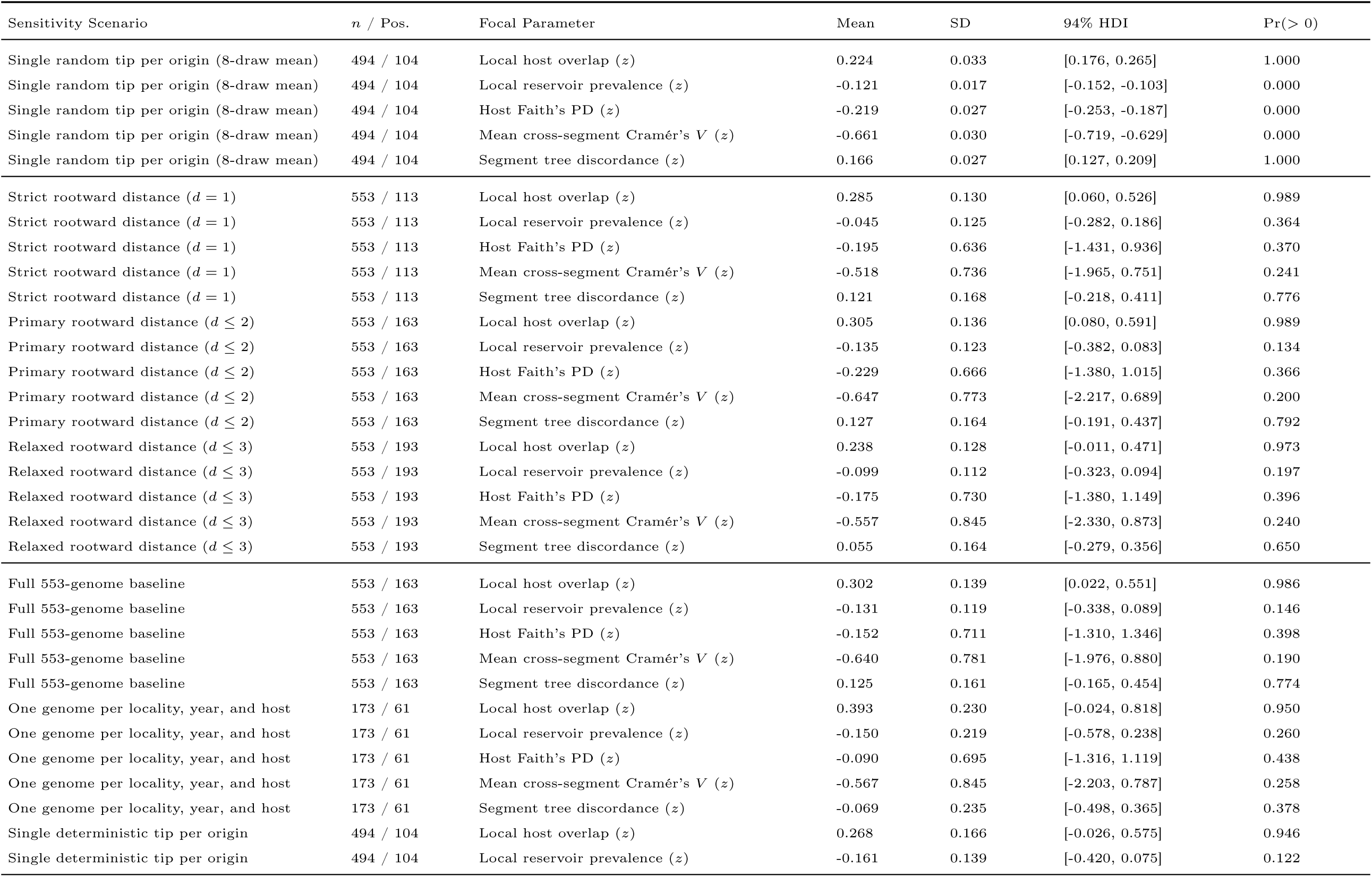
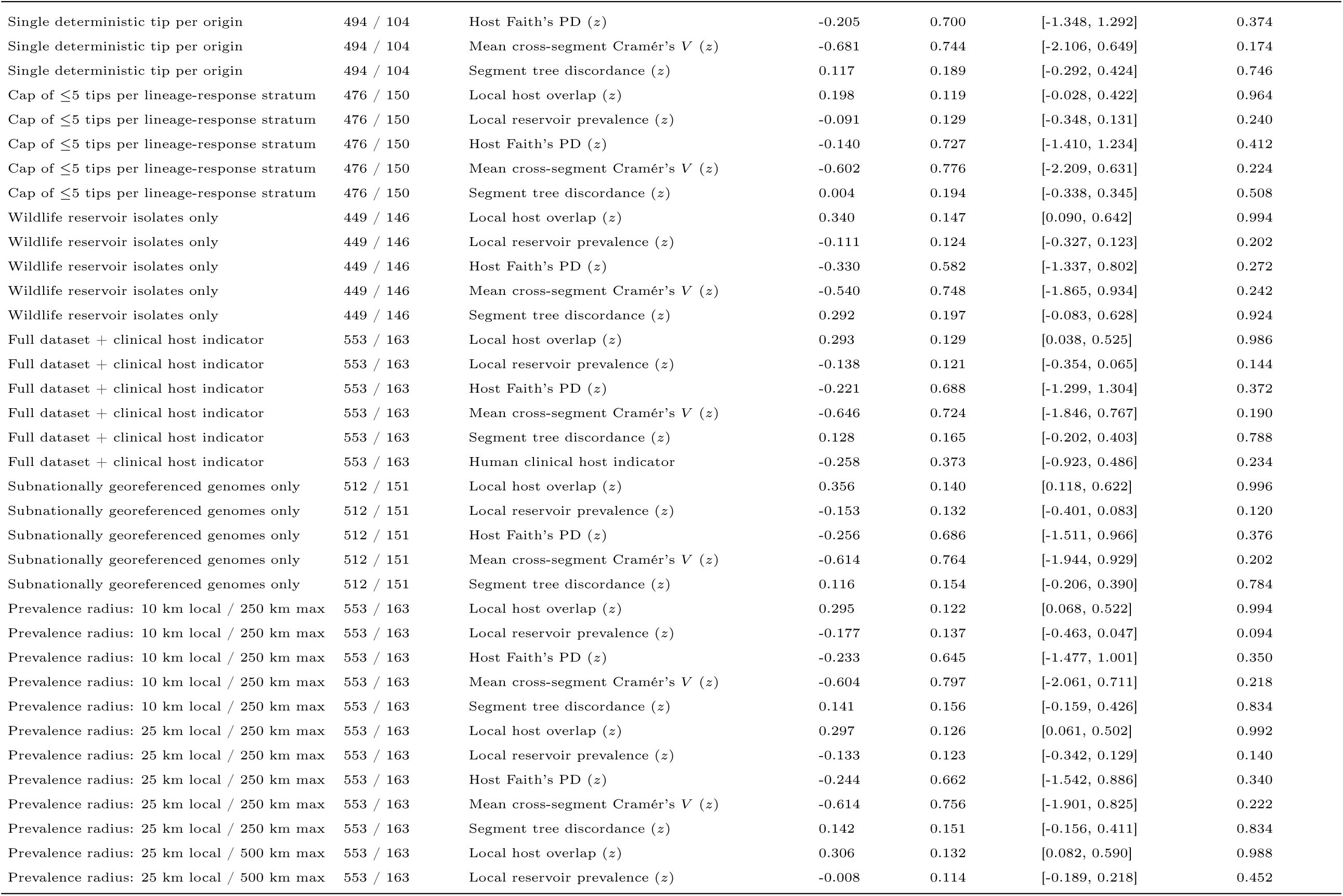
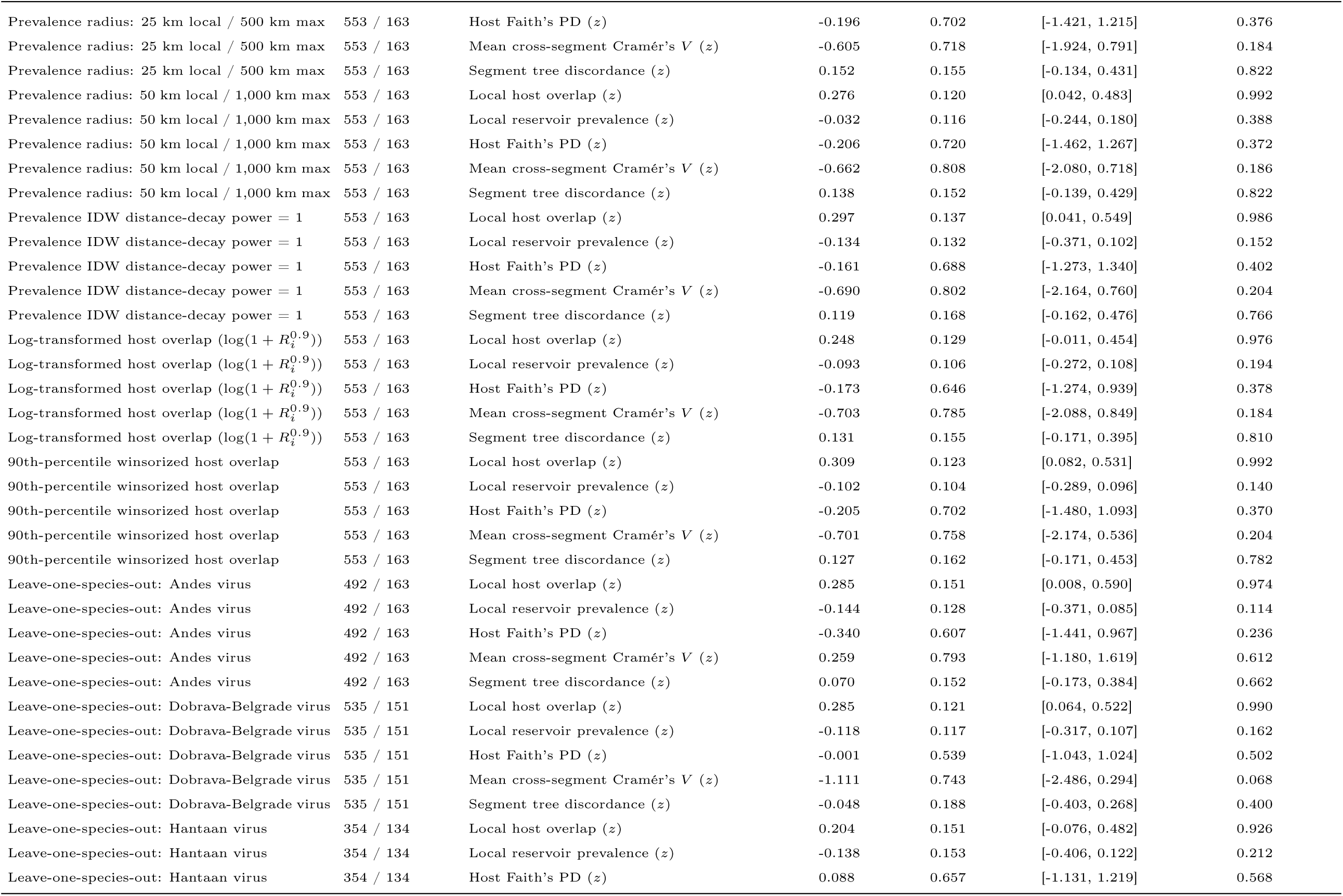
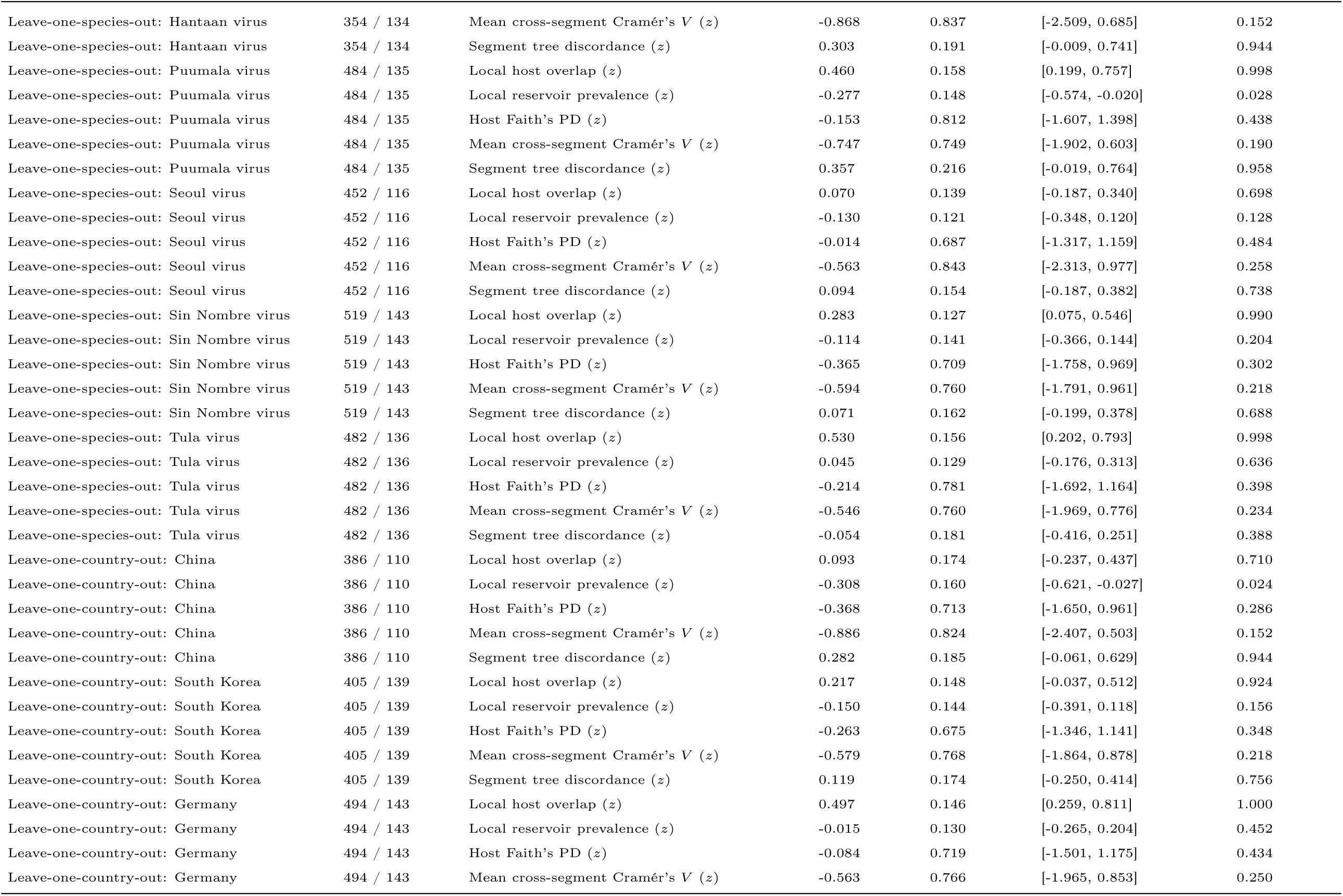
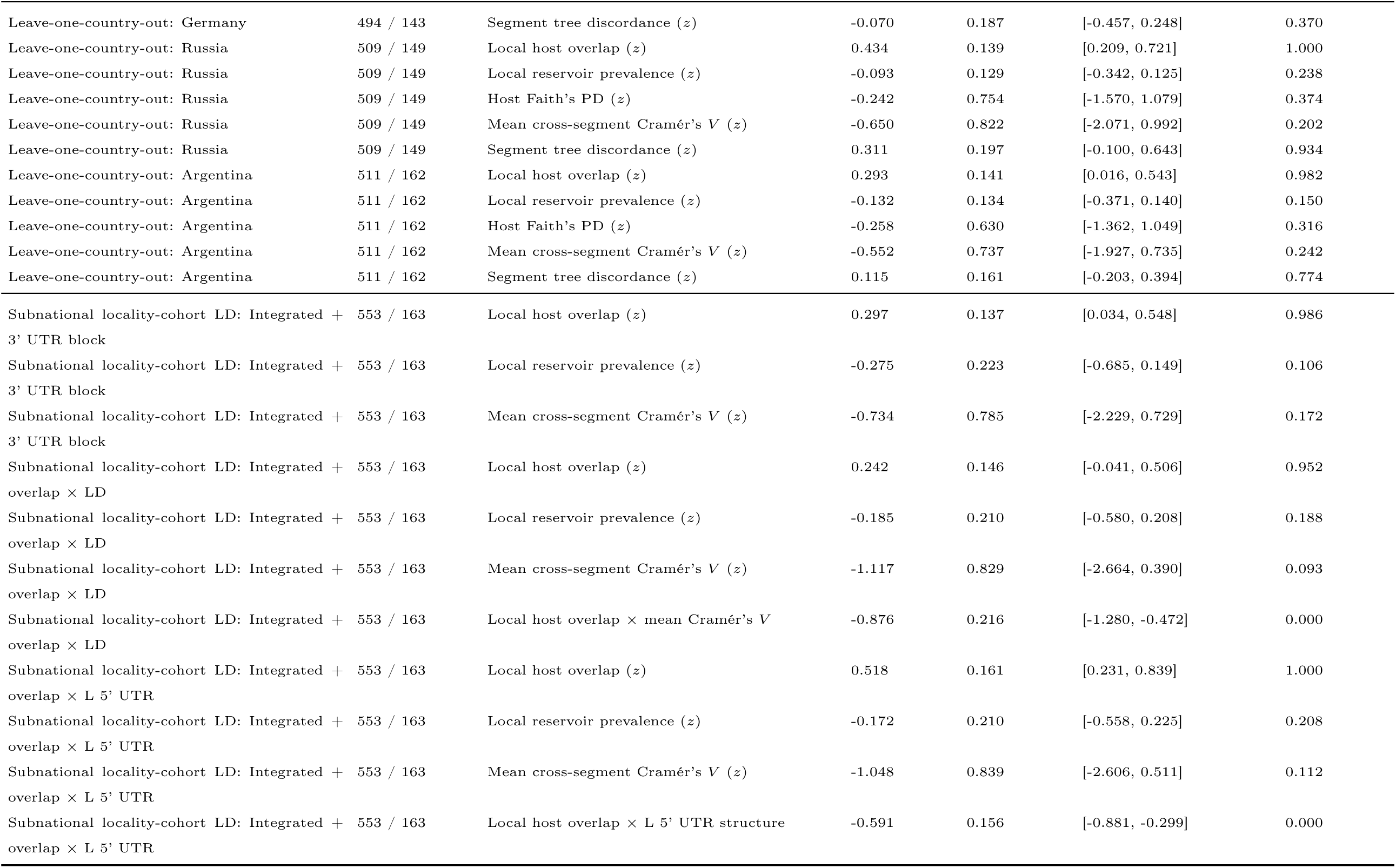
Sensitivity of Bayesian hierarchical regression coefficients across clonal tip-collapse, outbreak deduplication, rootward path distance thresholds (*d* = 1*, d ≤* 2*, d ≤* 3), non-human reservoir vs. clinical host provenance, subnational georeferencing, subnational locality-cohort rebuilt LD, local prevalence spatial interpolation radii (10–1,000 km and IDW power), overlap functional transformations, and leave-one-species-out / leave-one-country-out refits on the canonical *n* = 553 dataset.

## Notes

https://zenodo.org/records/20573223

https://github.com/RicardoRH96/hantavirus-reassortment

### Competing Interest Statement

The authors have declared no competing interest.

### Summary of Updates

This revision enhances the text's clarity and adds the sensitivity analyses requested during the journal's first round of revisions.

